# GPCRs show widespread differential mRNA expression and frequent mutation and copy number variation in solid tumors

**DOI:** 10.1101/546481

**Authors:** Krishna Sriram, Kevin Moyung, Ross Corriden, Hannah Carter, Paul A. Insel

**Affiliations:** Departments of Pharmacology, University of California, San Diego; Departments of Medicine, University of California, San Diego

## Abstract

G protein-coupled receptors (GPCRs) are the most widely targeted gene family for FDA-approved drugs. To assess possible roles for GPCRs in cancer, we analyzed The Cancer Genome Atlas (TCGA) data for mRNA expression, mutations, and copy number variation (CNV) in 20 categories/45 sub-types of solid tumors and quantified differential expression of GPCRs by comparing tumors against normal tissue from the GTEx database. GPCRs are over-represented among coding genes with elevated expression in solid tumors. This analysis reveals that most tumor types differentially express >50 GPCRs, including many targets for approved drugs, hitherto largely unrecognized as targets of interest in cancer. GPCR mRNA signatures characterize specific tumor types and correlate with expression of cancer-related pathways. Tumor GPCR mRNA signatures have prognostic relevance for survival and correlate with expression of numerous cancer-related genes and pathways. GPCR expression in tumors is largely independent of staging/grading/metastasis/driver mutations. GPCRs expressed in cancer cell lines largely parallels GPCR expression in tumors. Certain GPCRs are frequently mutated and appear to be hotspots, serving as bellwethers of accumulated genomic damage. CNV of GPCRs is common but does not generally correlate with mRNA expression. Our results suggest a previously under-appreciated role for GPCRs in cancer, perhaps as functional oncogenes, biomarkers, surface antigens and pharmacological targets.

**List of abbreviations/acronyms:** CAFs: Cancer associated fibroblasts; CCLE: Cancer Cell Line Encyclopedia; CNV/CNA: Copy number variation/ amplification; DE: Differential Expression; GPCR: G protein-coupled receptor; GTEx: Gene Tissue Expression Project [1]; GtoPdb: IUPHAR/BPS Guide to Pharmacology [2]; MDS: Multi-dimensional Scaling; Nmut: Number of genes with somatic non-silent mutations per tumor genome; OE: Overexpression; TCGA: The Cancer Genome Atlas; Table 1 lists abbreviated cancer names.

## Introduction

G protein-coupled receptors (GPCRs), the largest family of cell-surface receptors (>800 in the human genome), mediate the signaling of a wide variety of ligands, including hormones, neurotransmitters, proteases, lipids, and peptides. GPCRs regulate many functions (e.g., metabolism, migration, proliferation) and interactions of cells with their environment. GPCRs are also the largest family of targets for approved drugs [3,4], interacting with ~35% of FDA approved drugs, but are infrequently targeted in tumors other than endocrine cancers, even though a role for GPCRs has been implicated in features of the malignant phenotype [5,6]. One reason for their limited use is the notion that GPCRs are rarely mutated in cancer [7,8] although mutations occur in heterotrimeric GTP binding (G) proteins that GPCRs activate [8] and GPCRs regulate pathways, such as Wnt, MAPK and PI3K signaling, with mutations in cancer [9]. The biological relevance of GPCRs for the malignant phenotype and their high druggability imply that GPCRs might be an under-explored class of contributors to and targets in cancer.

To define the landscape of GPCRs in cancer, we undertook an integrated analysis of Differential Expression (DE), mutations, and CNV of GPCRs in 20 types of solid tumors (**Table 1, Supplementary Tables 1, 2**). Using RNA-seq data from TCGA and the GTEx database [1], we performed DE analysis of GPCRs in tumors compared to normal tissue, respectively, an analysis facilitated by the TOIL recompute project [10]. We studied GPCRs annotated by GtoPdb [2], including endoGPCRs (which respond to endogenous agonists) and taste receptors but not olfactory GPCRs for which such annotations are unavailable (**Supplement 2**). Our findings identify many differentially expressed GPCRs in solid tumors and corresponding cancer cell lines but a less important role for mutations and CNV. GPCRs with DE predict survival and are associated with expression of oncogenes and tumorigenic pathways. Overall, these results reveal a largely under-appreciated potential of GPCRs as contributors to cancer biology and as potential therapeutic targets. Our results from DE (N = 6224 individual tumors), mutation (N = 5103), and CNV analyses (N = 7545) are available as a resource at *<u>insellab.github.io</u>*.

**Table 1:** Tumors surveyed for Differential Expression (DE) analysis. TCGA cancer type and sub-classification, if applicable, for solid tumors with distinct histological classification are shown, along with the number of replicates and GPCRs with increased or decreased expression for each type of tumor.

|  | Cancer Type | Histology/Subtype | Replicates | # GPCRs ↑ | # GPCRs ↓ |
| --- | --- | --- | --- | --- | --- |
| 1 | Adrenocortical Cancer (ACC) | Adrenocortical carcinoma - Usual Type | 73 | 22 | 4 |
| 2 | Bladder Cancer (BLCA) | Papillary bladder cancer (BLCA_P) | 130 | 26 | 12 |
|  |  | Non-papillary bladder cancer (BLCA_NP) | 267 | 36 | 14 |
| 3 | Breast Cancer (BRCA) | Infiltrating ductal carcinoma (IDC), Her2 positive (BRCA_IDC_Her2+) | 48 | 46 | 25 |
|  |  | IDC, Hormone Receptor positive (BRCA_IDC_HR+) | 431 | 49 | 29 |
|  |  | IDC, Triple positive (BRCA_IDC_3pl+) | 54 | 51 | 25 |
|  |  | IDC, Triple negative (BRCA_IDC_3pl-) | 109 | 51 | 24 |
|  |  | Invasive Lobular Carcinoma (ILC), Hormone R positive (BRCA_Lob_HR+) | 57 | 50 | 31 |
| 4 | Cervical Cancer (CESC) | Cervical Squamous Cell Carcinoma (CESC_CervSq) | 252 | 40 | 18 |
|  |  | Endocervical Adenocarcinoma of the Usual Type (CESC_ECAD) | 21 | 38 | 23 |
|  |  | Mucinous Adenocarcinoma of Endocervical Type (CESC_Muc) | 17 | 38 | 24 |
| 5 | Colon Cancer (COAD) | Colon Adenocarcinoma in the sigmoid colon (COAD_Sig) | 71 | 41 | 28 |
|  |  | Colon Adenocarcinoma in the transverse colon (COAD_Trans) | 22 | 31 | 23 |
| 6 | Esophageal Cancer (ESCA) | Esophagus Adenocarcinoma (ESCA_AD) | 89 | 59 | 17 |
|  |  | Esophagus Squamous Cell Carcinoma (ESCA_SQC) | 92 | 43 | 9 |
| 7 | Kidney Papillary Cell Carcinoma (KIRP) | - | 288 | 28 | 14 |
| 8 | Kidney Clear Cell Carcinoma (KIRC) | - | 523 | 65 | 11 |
| 9 | Kidney Chromophobe (KICH) | - | 66 | 29 | 10 |
| 10 | Liver Cancer (LIHC) | Liver Hepatocellular Carcinoma (LIHC) | 360 | 11 | 7 |
| 11 | Lung adenocarcinoma (LUAD) | Lung Papillary Adenocarcinoma (LUAD_Pap) | 23 | 22 | 33 |
|  |  | Lung Bronchioloalveolar Carcinoma Non-Mucinous (LUAD_BCNM) | 19 | 34 | 36 |
|  |  | Lung Adenocarcinoma- Not Otherwise Specified (LUAD_NOS) | 308 | 33 | 33 |
|  |  | Lung Adenocarcinoma – Mixed (LUAD_Mixed) | 105 | 29 | 31 |
|  |  | Lung Acinar Adenocarcinoma (LUAD_Acinar) | 18 | 27 | 36 |
| 12 | Lung squamous cell carcinoma (LSQC) | Lung Squamous Cell Carcinoma- Not Otherwise Specified (LSQC_NOS) | 468 | 34 | 31 |
|  |  | Lung Basaloid Squamous Cell Carcinoma (LSQC_Basal) | 14 | 38 | 39 |
| 13 | Skin Cutaneous Melanoma (SKCM) | Primary melanomas (SKCM_Primary) | 100 | 34 | 18 |
|  |  | Distant metastases (SKCM_DMet) | 68 | 41 | 11 |
| 14 | Ovarian Cancer (OV) | Ovarian Serous Cystadenocarcinoma (OV) | 418 | 57 | 11 |
| 15 | Pancreatic Cancer (PAAD) | Pancreatic Ductal Adenocarcinoma (PDAC) | 147 | 68 | 11 |
| 16 | Prostate Cancer (PRAD) | Prostate Adenocarcinoma Acinar Type (PRAD) | 475 | 27 | 25 |
| 17 | Stomach Cancer (STAD) | Stomach, Adenocarcinoma, Diffuse Type (STAD_Diff) | 68 | 53 | 12 |
|  |  | Stomach, Adenocarcinoma, Not Otherwise Specified (STAD_NOS) | 154 | 48 | 14 |
|  |  | Stomach, Intestinal Adenocarcinoma, Mucinous Type (STAD_Muc) | 19 | 55 | 15 |
|  |  | Stomach, Intestinal Adenocarcinoma, Not Otherwise Specified (STAD_IntNOS) | 73 | 40 | 16 |
|  |  | Stomach, Intestinal Adenocarcinoma, Tubular Type (STAD_IntTub) | 76 | 39 | 12 |
|  |  | Stomach Adenocarcinoma, Signet Ring Type (STAD_Sig) | 12 | 37 | 10 |
| 18 | Testicular Cancer (TGCT) | Seminoma (TGCT_Sem) | 72 | 73 | 16 |
|  |  | Non-seminoma (TGCT_NonSem) | 65 | 76 | 9 |
| 19 | Thyroid Cancer (THCA) | Thyroid Papillary Carcinoma - Classical/usual (THCA_Usual) | 358 | 33 | 15 |
|  |  | Thyroid Papillary Carcinoma - Follicular (>= 99% follicular patterned) (THCA_fol) | 101 | 17 | 6 |
|  |  | Thyroid Papillary Carcinoma - Tall Cell (>= 50% tall cell features) (THCA_TC) | 36 | 33 | 17 |
| 20 | Uterine Carcinosarcoma (UCS) | Uterine Carcinosarcoma/Malignant Mixed Mullerian Tumor (MMMT): (UCS_NOS) | 24 | 43 | 19 |
|  |  | Uterine Carcinosarcoma/ MMMT: Heterologous Type (UCS_Het) | 20 | 44 | 22 |
|  |  | Uterine Carcinosarcoma/MMMT: Homologous Type (UCS_Homo) | 13 | 43 | 20 |

## Results

### Differential expression (DE) of GPCRs in solid tumors compared to normal tissues

We focused on GPCRs with both substantial DE and magnitude of expression in solid tumors, i.e., 1) > 2-fold increase/decrease in DE in tumors compared to normal tissue, 2) FDR < 0.05 and 3) median expression in tumors > 1 TPM. We used the latter threshold for median expression in order to identify GPCRs that may be useful as therapeutic targets, for which higher expression is preferable. For DE analysis, we divided the 20 TCGA tumor types into 45 tumor subtypes (**Table 1**), based on histological classification of tumors in TCGA metadata. We found that different tumor subtypes within the same TCGA tumor classification have distinct GPCR expression, e.g., subtypes of breast cancer (BRCA), thyroid cancer (THCA) and esophageal cancer (ESCA) (**Figs S1A-F**).

**Figure 1A** shows a heatmap with fold changes (where statistically significant) for mRNA expression of all GPCR genes in solid tumors, relative to their corresponding normal tissue. Hierarchical clustering was performed on the GPCR genes, revealing three main clusters of GPCRs: a) those with frequent increases in expression across multiple tumor types; b) those with relatively little change; and c) GPCRs frequently reduced in expression in solid tumors compared to normal tissue. A phylogenetic tree identifying the GPCRs in each cluster is shown in **Fig S1**. Among the most lethal forms of cancer (in terms of annual deaths), **Figure 1B** shows that >25 GPCRs have >2-fold increased expression relative to normal tissue (with median expression in tumors > 1 TPM), whereas >20 GPCRs have significantly down-regulated expression, while remaining expressed > 1 TPM in solid tumors. Thus, large numbers of GPCRs that are expressed in solid tumors show DE, including tumor types that are most lethal. **Figure 1C** shows the 20 GPCRs that have >2-fold increased expression (with median expression >1 TPM) in the largest number of tumor types. **Figure 1D** shows the same, but for GPCRs frequently reduced >2-fold in expression, but which are still detected at >1 TPM in those tumors. Among the GPCRs with frequently increased expression are receptors likely expressed in the tumor cells themselves (e.g., GPRC5A, [11,12]) and expressed in the tumor microenvironment, such as in fibroblasts (e.g., F2R, [13]) and immune cells (e.g., FPR3, CCR1, CCR5).

**Figure 1.**
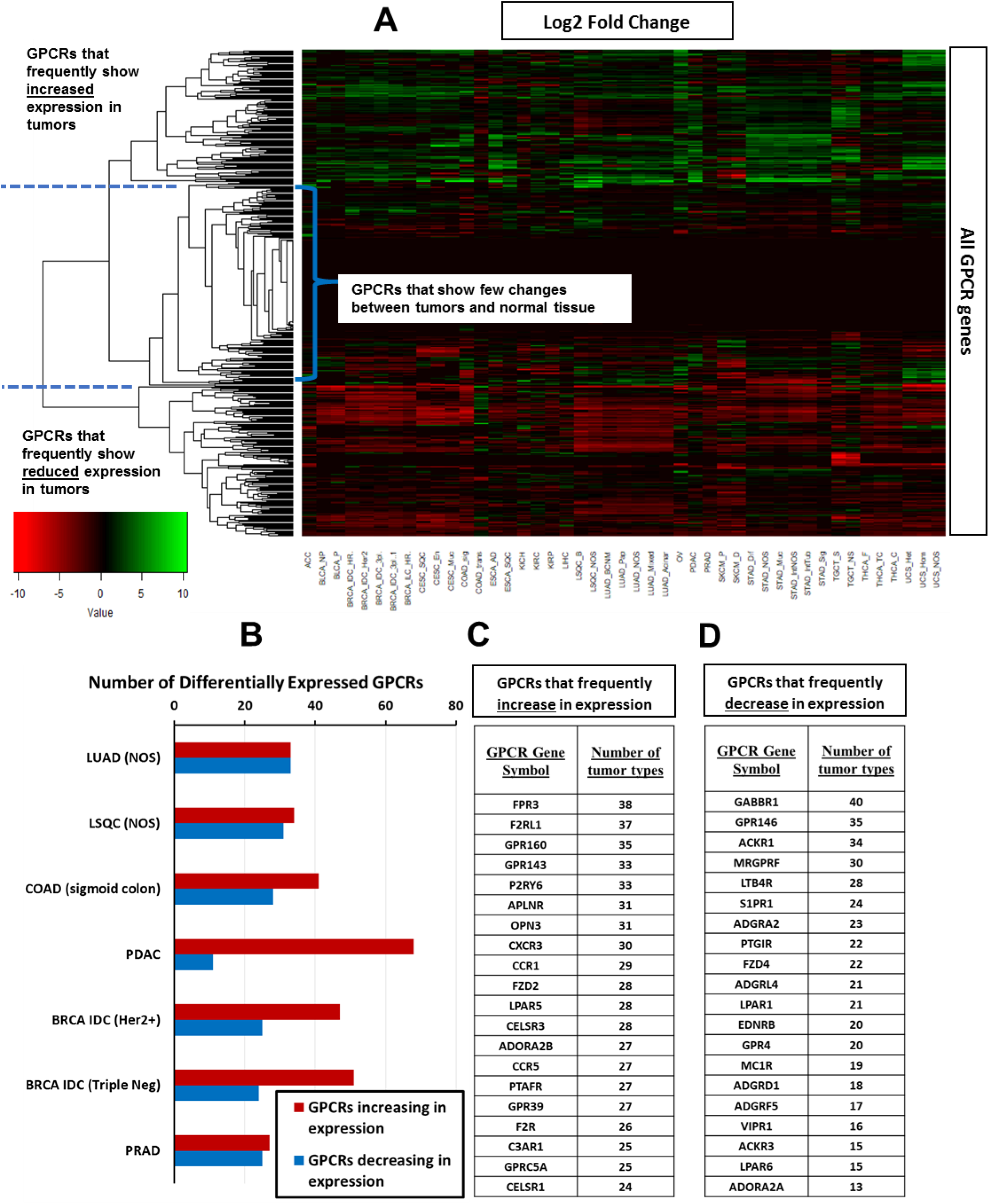
Heatmaps of GPCR expression and differential expression (DE) in solid tumors. (**A**) For all 45 tumor subtypes, a heatmap showing the Log2 Fold-Change of GPCR expression in tumors compared to normal tissue (positive values indicate higher expression in tumors), with hierarchical clustering of GPCR genes to reveal patterns of DE. (**B**) The number of GPCRs that show significant (FDR < 0.05) changes in expression compared to normal tissue among tumor types tested with large numbers of replicates (**Table 1**) and that correspond to the most lethal types of cancer. (**C-D**) The GPCRs that most frequently (i.e., in most tumor types) show increases (**C**) or decreases (**D**) in expression among the 45 tumor subtypes

**Figure S3A-C** shows DE for pancreatic ductal adenocarcinoma (PDAC) tumors (as an example) compared to normal pancreas. An MDS plot (**Fig S3A**) reveals clusters for tumors and normal tissue, implying distinct transcriptomic profiles. The more diffuse cluster of PDAC samples likely reflects their heterogeneity. Smear and volcano plots (**Figs S3B-C**) reveal many genes (>5000) with high, statistically significant DE (FDR <<0.05). **Figures S3D-F** show examples of genes (other than GPCRs) with high overexpression that prior studies implicated as having a role in PDAC. Multiple other tumor types also show expression of genes relevant to the malignant phenotype, cluster separately from their respective normal tissues, and have a large number of genes with DE, thus supporting the validity of our analysis.

Many GPCRs show DE in tumors, including those from each GPCR class: A (rhodopsin-like), B (secretin-like), C (metabotropic glutamate and others), frizzled and adhesion GPCRs. The highest expressed GPCRs in PDAC tumors (as an example, this finding is generalizable to other tumor types) are generally overexpressed compared to normal tissue and include orphan receptors (e.g., *GPRC5A* and *ADGRF4*/*GPR115*) and GPCRs with known agonists (e.g., *GPR68*) (**Fig 2A, C**). *GPRC5A*, the most highly expressed GPCR in PDAC, is 50-fold higher expressed; 95% of PDAC samples have >8-fold higher median *GPRC5A* expression than in normal pancreas (**Fig S3I**). Within a tumor type, a large majority of individual tumors express such overexpressed GPCRs at far higher levels than corresponding normal tissue (**Fig 3A**, e.g., GPRC5A); a subset of GPCRs are expressed in >90% of PDAC tumors at abundances greater than in any normal pancreas sample (**Fig 2F**). As discussed in subsequent sections, **Figures 3B, D-F** (and **Figs 7A-E**) indicate that GPCR expression is relatively consistent, ubiquitous among members of a given tumor cohort and largely independent of patient characteristics (e.g. tumor grade, pathological T and Sex, **Figs 3D-F**) that one might use to stratify patients. In addition, the GPCRs expressed couple to all major classes of Gα G protein signaling mechanisms (**Fig 3C**).

**Figure 2.**
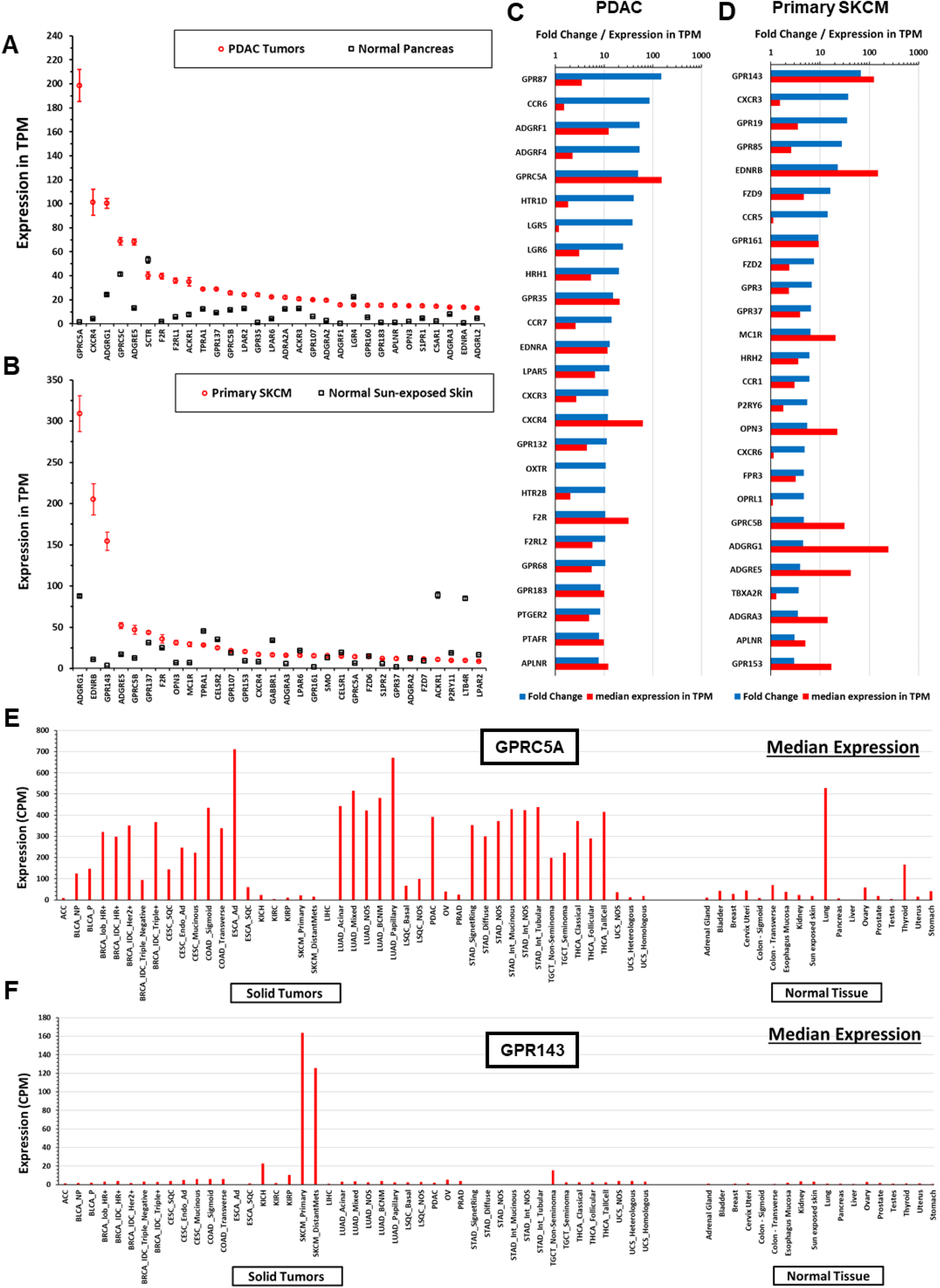
Solid tumors have large numbers of differentially expressed GPCRs compared to normal tissue. (**A-B**) The 30 highest expressed GPCRs in PDAC (**A**) or primary SKCM (**B**) and their corresponding expression in normal pancreatic (**A**) or skin (**B**) tissue. (**C-D**) The 30 GPCRs with the highest fold-increase in expression in tumors compared to normal tissue for (**C**) PDAC and (**D**) Primary SKCM, sorted by fold-increase in tumors compared to normal tissue. (**E-F**) For two highly expressed GPCRs in (**A-D**) as examples, the median expression of (**E**) GPRC5A and (**F**) GPR143 in all tumor types tested and corresponding normal tissue, normalized in CPM, allowing for comparison between tissue/tumor types. A lookup file that enables generation of similar plots (as well as upper and lower quartiles of expression) for any GPCR can be found at https://insellab.github.io/gpcr_tcga_exp.

**Figure 3.**
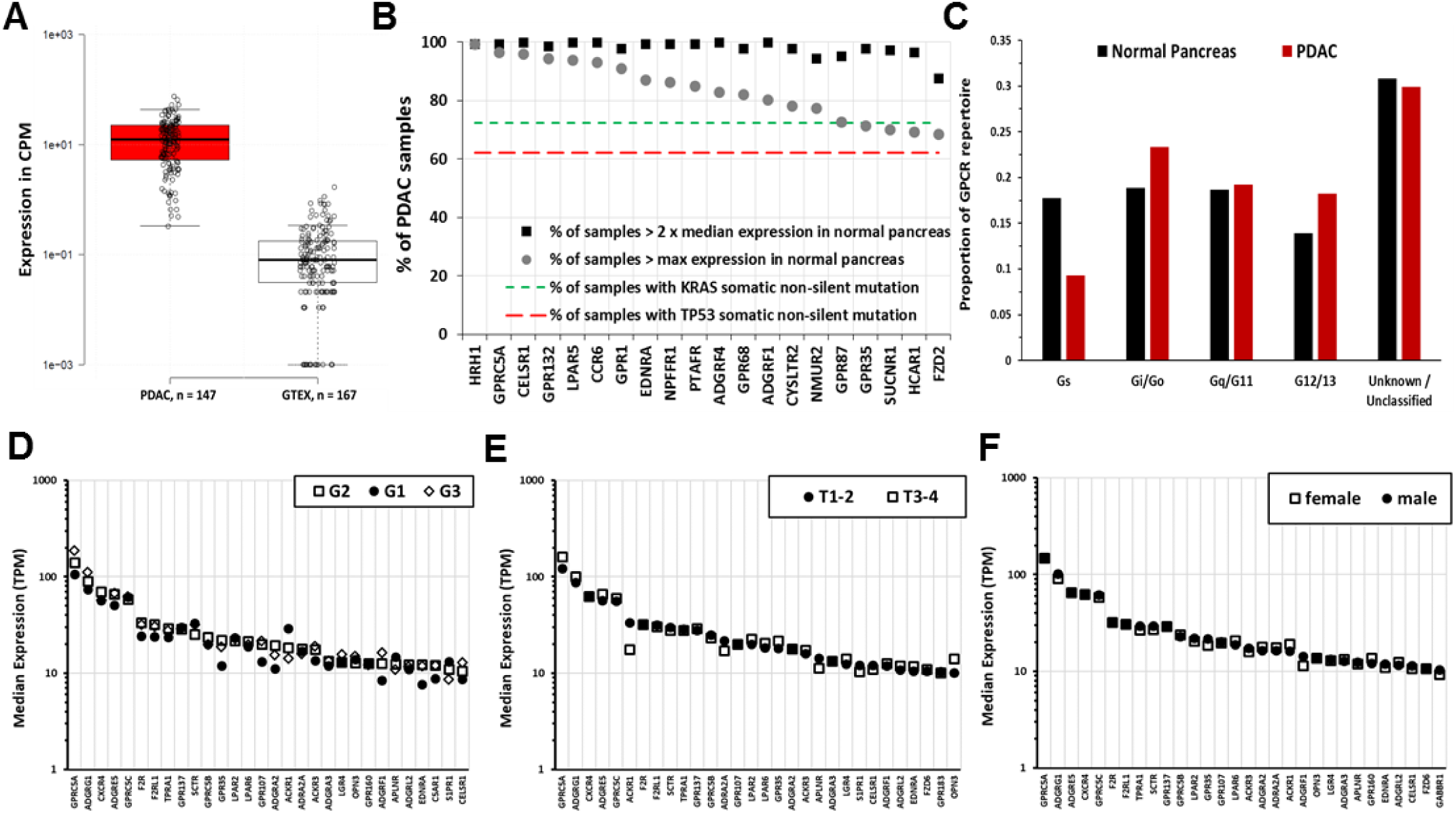
Increased expression of certain GPCRs in PDAC is more frequent than that of KRAS and TP53 mutations, is independent of tumor grade, pathological T, and patient sex and includes GPCRs that couple to each of the 4 major G protein families. (**A**) The expression of GPRC5A in all PDAC samples and normal pancreas tissues analyzed. (**B**) Frequency of 2-fold increase and percent of TCGA-PDAC samples with higher maximal expression compared to normal pancreas of the indicated GPCRs with comparison of the frequency of mutations of KRAS and TP53, the most frequent somatic, non-silent mutations in PDAC tumors in TCGA. (**C**) Changes in GPCR expression in PDAC alter the GPCR repertoire that couple to different G proteins. (**D-F**) Tumor grade (D), Pathological T (E) and patient sex (F) do not impact on GPCR expression. The 30 highest expressed GPCRs in PDAC tumors are shown for each case; no statistically significant differences occur between the groups.

Similar to PDAC, numerous GPCRs are highly, consistently overexpressed in other tumor types. For instance, in skin cutaneous melanoma (SKCM) (**Fig 2B, D**); *ADGRG1/GPR56*, *GPR143* and *EDNRB* are highly overexpressed and highly expressed in magnitude compared to normal skin in >90% of melanoma samples (**Figs S3G-I**). In general, such highly overexpressed GPCRs are expressed in the vast majority, typically >90%, of samples within a tumor subtype.

**Figure 2E** shows the median expression of GPRC5A, the highest expressed GPCR in PDAC (and highly differentially expressed, **Figs 2A, C**) and highly expressed in many adenocarcinomas. **Figure 2F** shows this analysis across all tumor types and normal tissue for GPR143, the highest up-regulated GPCR in SKCM. For many GPCRs, we observe similar patterns of expression with pronounced up-regulation in tumors, as shown in **Figure 1**. To facilitate exploration of these data, we provide a spreadsheet-based tool (downloadable at https://insellab.github.io/gpcr_tcga_exp) wherein users can generate similar plots (along with upper and lower quartiles of expression) as in **Figure 2E-F** for any GPCR gene of interest. Of note, overexpression of certain GPCRs tends to be more prevalent within specific tumor types/subtypes than are common mutations. For example, *KRAS* and *TP53* are the most frequently mutated genes in PDAC (>70% and >60% of TCGA samples, respectively) but increased expression of multiple GPCRs occurs with greater frequency (**Fig 3B**). Each GPCR shown in **Figure 3B** shows statistically elevated expression in tumors compared to normal tissue, with FDRs << 0.05. The magnitude of DE and corresponding FDR for each GPCR shown are provided in **Supplement 3** and the project website.

### A resource for exploring GPCR expression in tumors and normal tissue

We compiled a list of GPCRs overexpressed in solid tumors with fold-changes and FDR along with expression in TPM (for median expression and within-group comparisons of different genes) and CPM (for inter-group comparisons of the same gene). The analysis revealed that 35 of 45 tumor types/subtypes show increased expression of >30 GPCRs; 203 GPCRs are overexpressed in at least one type of cancer (**Supplementary Table 4; Supplement 3**), including 47 orphan GPCRs and >15 GPCRs that couple to each of the major G protein classes. Increased expression of 130 GPCRs occurs in ≥4 tumor subtypes (**Supplement 3**). A subset of GPCRs is overexpressed in many tumors, e.g., *FPR3* in 38 of the 45 tumor categories. **Supplementary Table 4** lists other examples along with GPCRs that have reduced expression compared to normal tissue. Importantly, of the 203 GPCRs with increased expression in one or more tumors, 77 are targets for approved drugs. These include *ADORA2B*, *CCR5* and *F2R*, which are overexpressed in 27, 27 and 26 tumor subtypes, respectively. **Table S4** and **Supplement 3** provide further details regarding such druggable GPCRs.

Data generated and mined in this study, including DE analysis, re-normalized GPCR expression data, expression of every GPCR in every individual tumor sample analyzed, accompanying CNV data and mutation data for GPCRs are provided in the online resource at https://insellab.github.io. This resource also includes plots of GPCR expression in tumors and corresponding normal tissue (such as those shown in **Figure 2A** and **2C**), MDS scaling plots showing the extent to which tumor and normal samples cluster together based on overall gene expression, the aforementioned lookup tool to plot GPCR expression and information on GPCR expression in cancer cell lines. In addition, high-resolution images of the heatmaps and phylogenetic trees in **Figures 1A, 4A-B** and **S1** are available for download. Thus, while we have shown examples of GPCR expression and DE in certain tumor types in this text, the resource website provides similar information for all GPCRs in all tumor types.

**Figure 4.**
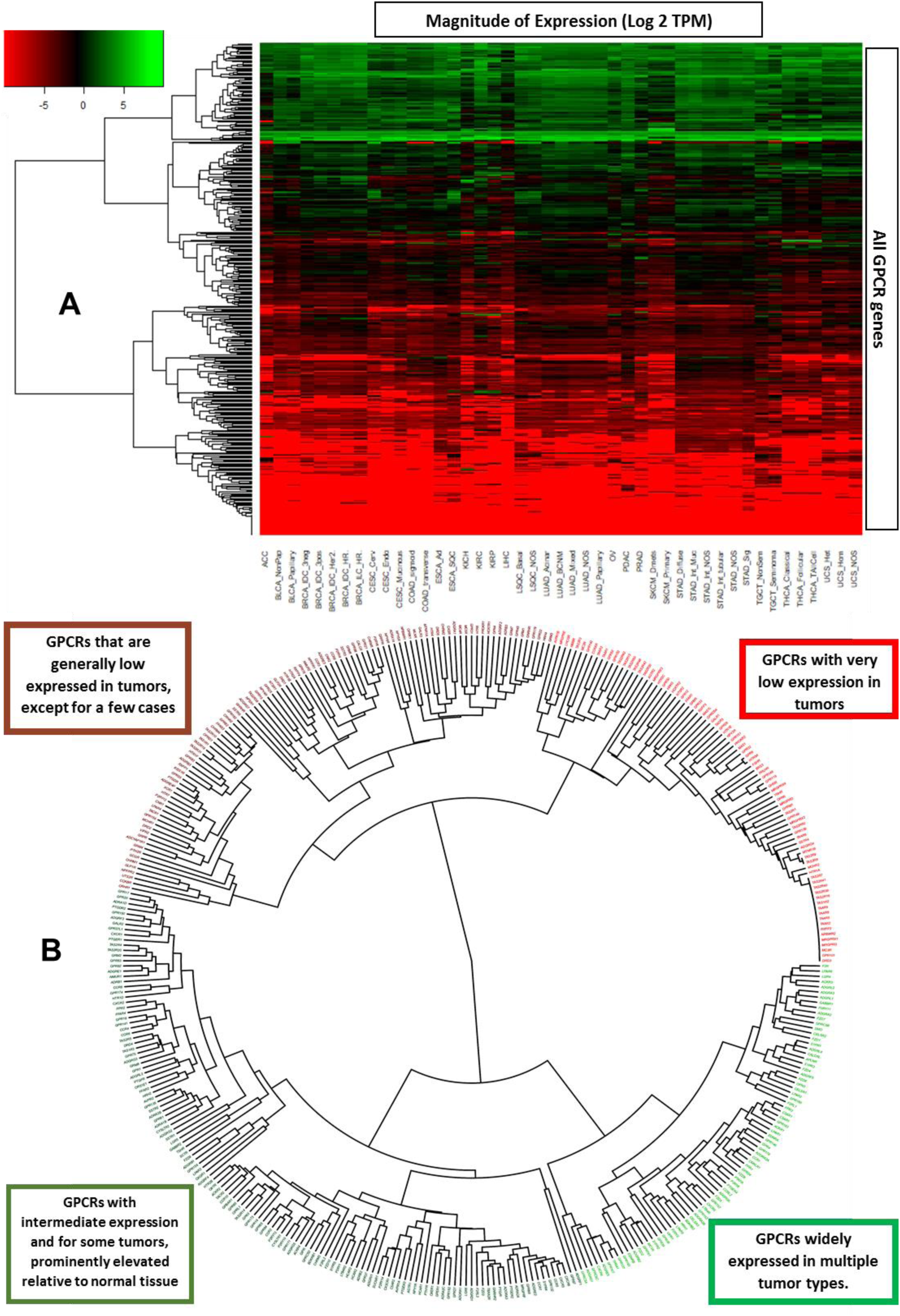
Grouping GPCRs and tumor types based on patterns of GPCR expression. (**A**) A phylogenetic tree showing the hierarchical clustering of GPCRs from Figure 1A reveals subsets of GPCRs that are either high- or low-expressed in solid tumors. (**B**) Hierarchical clustering of types of solid tumors, based on their expression (in CPM) of GPCRs reveals clusters of tumor types, characterized by expression of particular GPCRs.

### Patterns of GPCR expression across solid tumors

**Figure 4A** shows a heatmap that plots median GPCR expression in TPM across all tumor types, for all non-olfactory GPCRs and reveals that GPCRs can be divided into four groups: a) those widely expressed at high levels in all tumor types (often at >10 TPM) b) those with intermediate expression more broadly (~1 TPM), but with high expression in specific tumor types c) GPCRs with generally very low expression (~O(0.1) TPM or less) in most tumor types and d) GPCRs that are generally not detected in solid tumors. A phylogenetic (**Figure 4B**) tree shows the identities of these groups of GPCRs. In general, many of the GPCRs in **Figure 4** that are widely expressed in cancer are also widely differentially expressed (**Figure 1**).

We also performed hierarchical clustering on tumor types to explore whether GPCR expression is distinct in different subsets of solid tumors. **Figure 5** shows a phylogenetic tree of solid tumors, based on hierarchical clustering of median GPCR expression in CPM (thereby allowing for comparisons between groups) in each tumor type. Certain types of tumors cluster together, in a manner that corresponds more generally to their biology. For example, nearly all adenocarcinomas cluster together and express a common set of GPCRs (**Figure 5**). Similar findings are observed for the other clusters. All tumor types within each group had a median expression of these GPCRs ≥ 10 TPM, implying a common GPCR profile for adenocarcinomas compared to squamous cell carcinomas or other groups highlighted. Such widely expressed GPCRs may be potential drug targets. Several of these GPCRs commonly appear across multiple groups/clusters, but at different levels of expression. For example, GPRC5A is widely expressed in both adenocarcinomas and squamous cell carcinomas but is typically higher expressed in adenocarcinomas.

**Figure 5.**
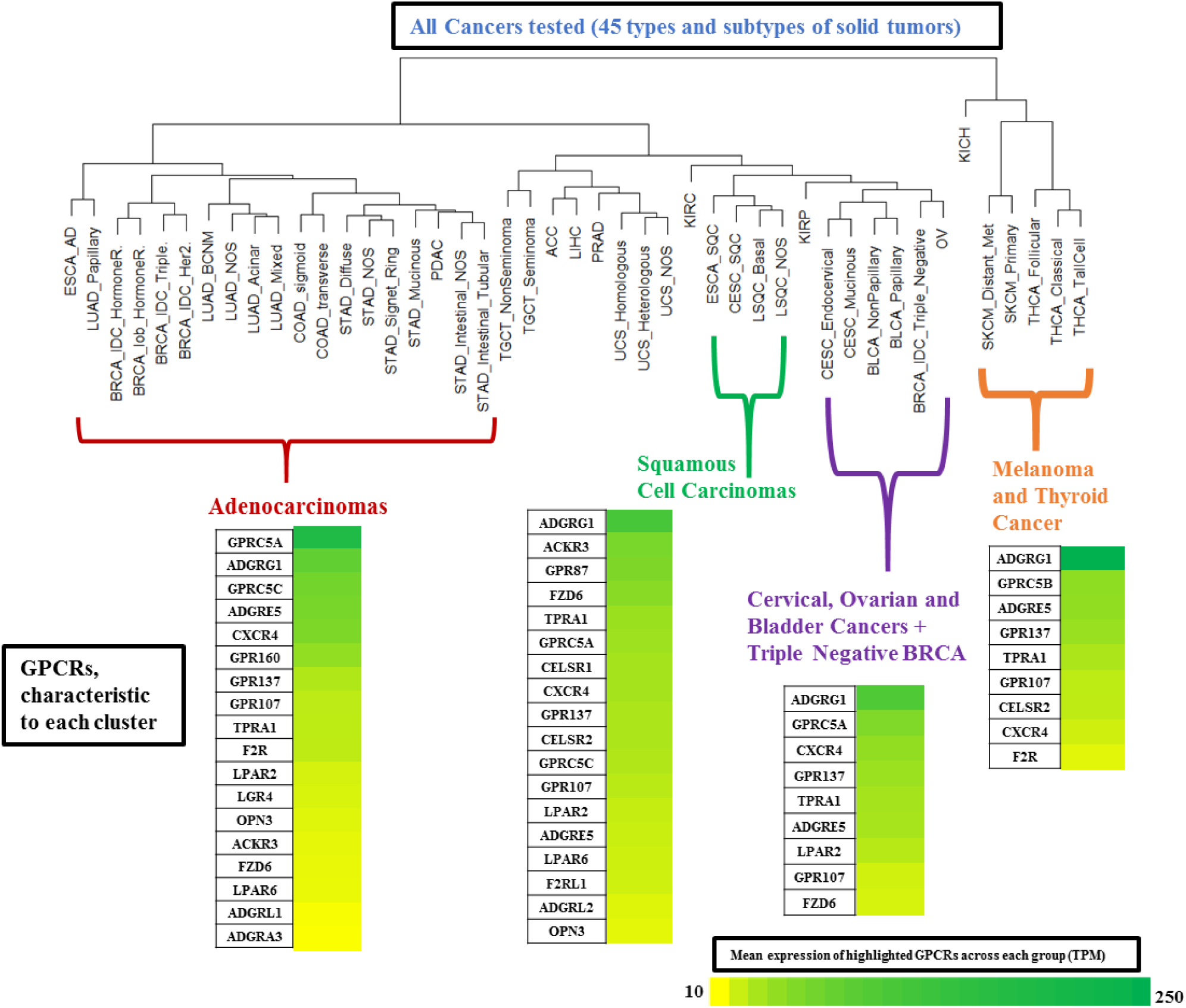
Hierarchical clustering of types of solid tumors, based on their expression (in CPM) of GPCRs reveals clusters of tumor types with expression of various GPCRs at characteristic levels of expression.

### GPCR expression is associated with cancer-related pathways and with survival: PDAC as an example

A subset of GPCRs in PDAC is highly overexpressed and prominently expressed in tumors and in PDAC cells (*vide infra*). Combining expression (normalized to median expression in PDAC) of 5 of the most highly DE GPCRs (ADGRF1, ADGRF4, GPRC5A, HRH1 and LPAR5) yields a composite ‘marker’ whose expression positively correlates with a subset of ~1200 genes (Benjamini-Hochberg adjusted p-values < 0.001) (**Fig S5A**). A reconstruction of the resulting network of genes using STRING [14] is shown in **Figure 6A**.

**Figure 6.**
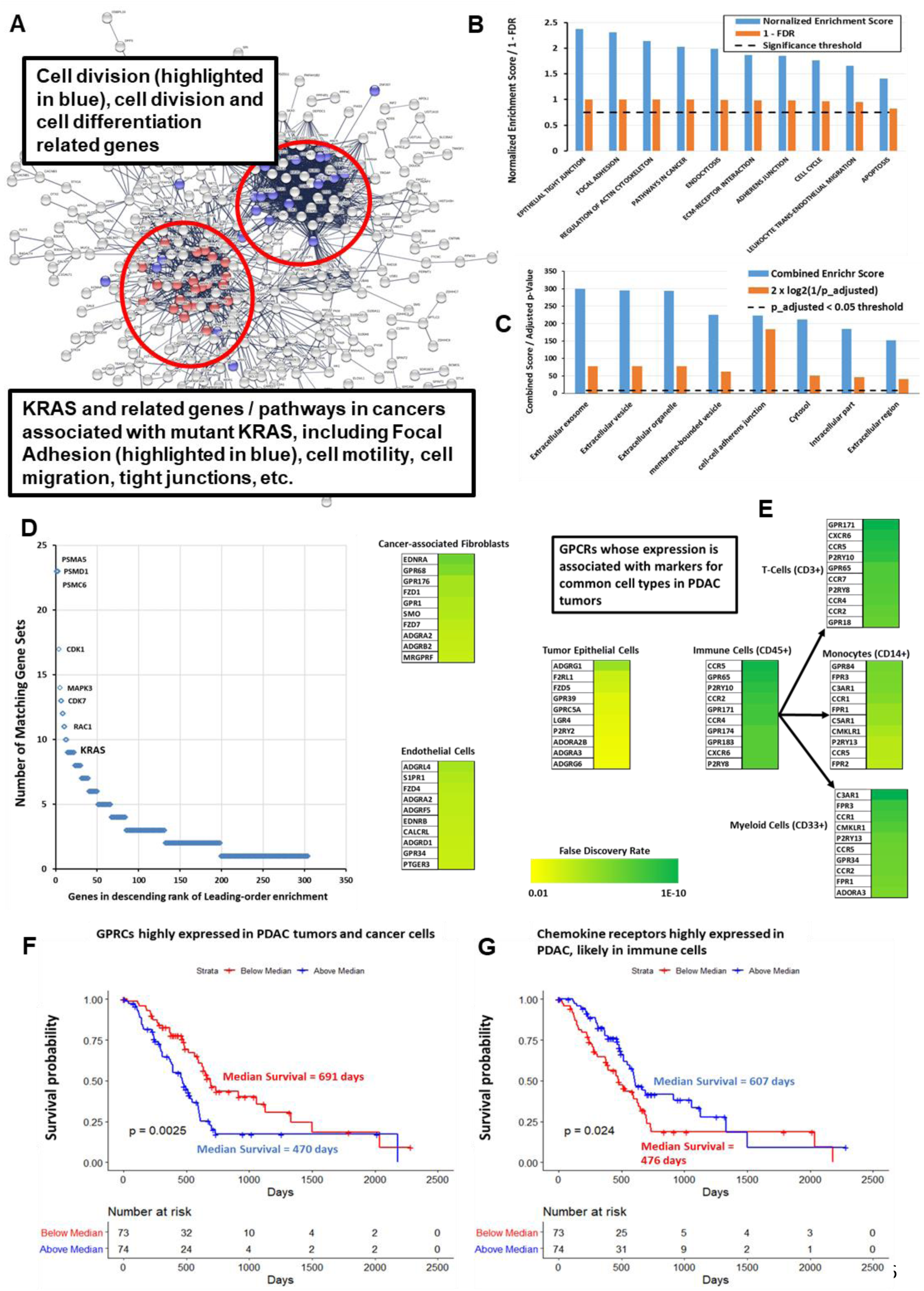
GPCRs expression correlates with cancer-related pathways and predicts survival in PDAC. (**A**) Network analysis via STRING [14] of the genes positively correlated with expression of GPCRs highly expressed in PDAC, with FDR < 0.001, revealing distinct clusters of genes associated with specific cancer-related pathways. (**B**) KEGG [16] gene sets with positive enrichment among genes most significantly positively or negatively correlated with expression of the identified subset of overexpressed GPCRs, based on GSEA pre-ranked analysis [15]. (**C**) Enrichr [19] analysis of the genes positively correlated in (**A**), with FDR < 0.001, based on their enrichment in cellular compartments, from the Jensen compartment database [20]. (**D**) Leading-edge analysis of GSEA results from (**B**), showing numerous cancer-related genes, including KRAS, which are common to multiple enriched KEGG genes sets. (**E**) GPCRs in PDAC show a positive, statistically significant association (FDR < 0.001) between their expression and that of markers for various cell types as shown. The top 10 GPCRs associated with each cell type are highlighted, along with the strength/significance of these correlations. (**F, G**) Kaplan-Meier curves for the impact of combined, normalized expression of subsets of GPCRs on PDAC patients. Total number of patients = 141; 59 patients were censored due to inadequate follow-up. (**F**) Impact of highly expressed GPCRs on median survival: 652 days (if below median expression) and 470 days (if above median expression) and for (**G**) Impact of highly expressed immune-related GPCRs on median survival: 460 days (below median expression) and 603 days (above median expression).

We conducted further analyses related to PDAC, including with GSEA [15] of the sets of negatively and positively associated genes (with respect to the five gene marker described above), pre-ranked/weighted by their FDRs and found an enrichment of a number of KEGG [16] pathways relevant to cancer (**Fig 6B**). The set of positively associated genes shows similar associations with cancer-related pathways when analyzed via GO [17,18] and Enrichr [19]. Enrichr also identifies, based on the Jensen compartment database [20], an enrichment of vesicle and exosome-related gene products among the set of positively correlated genes (**Fig 6C**). Network-analysis of the genes positively associated with the composite GPCR marker via STRING [14] provides an intuitive picture of this gene set (**Fig 6A**). Two ‘clusters’ of genes and pathways are evident; those associated with KRAS (including KRAS itself) and related processes (e.g., focal adhesion pathways) and a second group associated with regulation of cell cycle, cell division and differentiation. Expression of highly overexpressed GPCRs is positively correlated with one another and with expression of KRAS, implicating this GPCR subset as a PDAC signature. Leading edge analysis of the GSEA results confirmed that KRAS and other oncogenes are common elements in multiple enriched gene sets associated with this GPCR signature (**Fig 6D**). Survival analysis indicates that GPCR expression has prognostic relevance: patients with above-median expression of the five GPCRs had a ~200-day shorter survival compared to those with less than the median expression (**Fig 6F; Fig S3A**).

We tested whether we could identify associations between GPCR expression in PDAC tumors and expression of markers for cell types commonly found in these tumors, thereby perhaps serving as an indicator for the cell types in the tumor microenvironment that express certain GPCRs. The cell types and markers we explored were: Tumor epithelial cells (E-Cadherin as a marker), cancer-associated fibroblasts (CAFs, Collagen1A1), endothelial cells (*VWF*, Von Willebrand Factor), the immune compartment (*CD45*), T-Cells (*CD3G*), myeloid cells (*CD33*) and macrophages (*CD14*). Pearson correlations were calculated between expression of each GPCR and of these markers, from which p-values were then calculated, followed by adjusted p-values using the Benjamini-Hochberg method so as to identify GPCRs associated with markers for each cell type. **Figure 6E** lists the most strongly correlated GPCRs for each cell type. For CAFs and cancer cells, these data appear in excellent agreement with prior experimental results. For example, *GPR68*/*OGR1* is strongly associated with CAFs in our analysis, consistent with evidence that it is a novel functional receptor in PDAC-derived CAFs [13]. Similarly, epithelial-enriched GPCRs (**Figure 6E**) are expressed in cancer cells [21] and as shown below, in cancer cell lines.

PDAC tumors have a subset of highly overexpressed (and highly expressed in terms of magnitude) chemokine receptors (CCR6, CCR7, CXCR3 and CXCR4) not expressed in PDAC cancer cells but likely associated with immune cells and their activation. Genes that correlated with expression of these GPCRs are involved with immune-associated processes, especially T-cell and B-cell related pathways (**Fig S4B**). Combined expression of these GPCRs is a positive predictor of survival (**Fig 6G**). The observation that GPCR expression may be a marker for survival is not unique to PDAC (**Fig 9D-F**): In SKCM, ESCA and LIHC, expression of individual GPCRs is associated with survival but combinations of such GPCRs are even better predictors of survival. GPCR expression may thus serve as a prognostic indicator in multiple tumor types.

The finding that GPCRs with high expression and DE in tumors show an association with tumorigenic pathways appears generalizable. For example, GPCRs that are highly expressed/overexpressed in other adenocarcinomas (**Figure 5**) are positively correlated with expression of genes from pathways similar to ones shown in **Figure 6A-C** for PDAC; i.e., focal adhesion, cell motility, cell cycle and division. By contrast, GPCR expression of solid tumor types from different branches of the cancer/GPCR phylogenetic tree shows an association with different pathways from those observed in adenocarcinomas. For example, GPR143, EDNRB and ADGRG1 are highly expressed and differentially expressed in SKCM and are adverse indicators of survival (**Figs S2F-H and Fig 9D**). Pathways enriched among genes that correlate with expression of the GPCRs are ones implicated in metastatic SKCM (**Figs S5 A-C**), such as transferrin transport [22], melanosome organization [23] and insulin receptor signaling [24]. In general, highly expressed GPCRs in solid tumors show a positive correlation between GPCR expression and expression of tumorigenic pathways, implicating these GPCRs as potentially “functional oncogenes”.

### Functionality of overexpressed GPCRs

Evidence for functional roles in cancer cells of GPCRs that are highly expressed and overexpressed in solid tumors and cancer cells include findings for *PAR1/F2R* in breast cancer [25], gastric cancer [26], colon cancer [27] and melanoma [28] cells and for *PAR2/F2RL1* in melanoma [28], breast [29] and colon cancer cells [30]. Higher *PAR2* expression in ovarian cancer predicts poorer prognosis [31]. *EDNRB*, which is highly overexpressed in SKCM, promotes migration and transformation of melanocytes and melanoma cells and inhibition of *EDNRB* is pro-apoptotic [32,33]. *GPR143* promotes migration [34] and chemotherapeutic resistance [35] of melanoma cells. *GPR160* and *GPRC5A*, two frequently overexpressed GPCRs, are orphan receptors that influence the malignant phenotype. Knockdown of *GPR160* in prostate cancer cells increases apoptosis and growth arrest [36]. It has been suggested that *GPRC5A* is an oncogene that promotes proliferation, migration and colony formation of PDAC cells [11,12]. GPCRs with increased expression may thus be functional in cancer cells and activated by endogenous agonists or have constitutive activity that regulates signaling via heterotrimeric G proteins and/or β-arrestin [4]. At least certain of the many overexpressed GPCRs may thus serve as phenotypic drivers.

Incorporating omics analysis similar to what is presented here, our laboratory has recently shown [13] that *GPR68* (a proton-sensing GPCR) is highly overexpressed in PDAC tumors, in particular, in CAFs. We validated these data at the protein level and discovered that *GPR68* mediates symbiotic crosstalk between CAFs and PDAC cells and contributes to the tumor phenotype. Such findings provide an example as to how such omics data can identify overexpressed GPCRs with relevance to cancer cells themselves and to other cells in the tumor microenvironment.

### Driver mutations, patient sex and stage/grade of tumors does not impact on GPCR expression

GPCR expression and DE is largely independent of tumor stage and grade. **Figure 3D** shows the similarity in GPCR expression for Grades 1 to 3 (G1 to G3) PDAC tumors. Median expression of GPCRs was also similar in PDAC tumors with different pathological T (**Fig 3E**). Similarly, Stage I and Stage IIIA BRCA IDC HR+ (Hormone Receptor positive) tumors have comparable GPCR expression and DE (**Fig 7D**).

**Figure 7.**
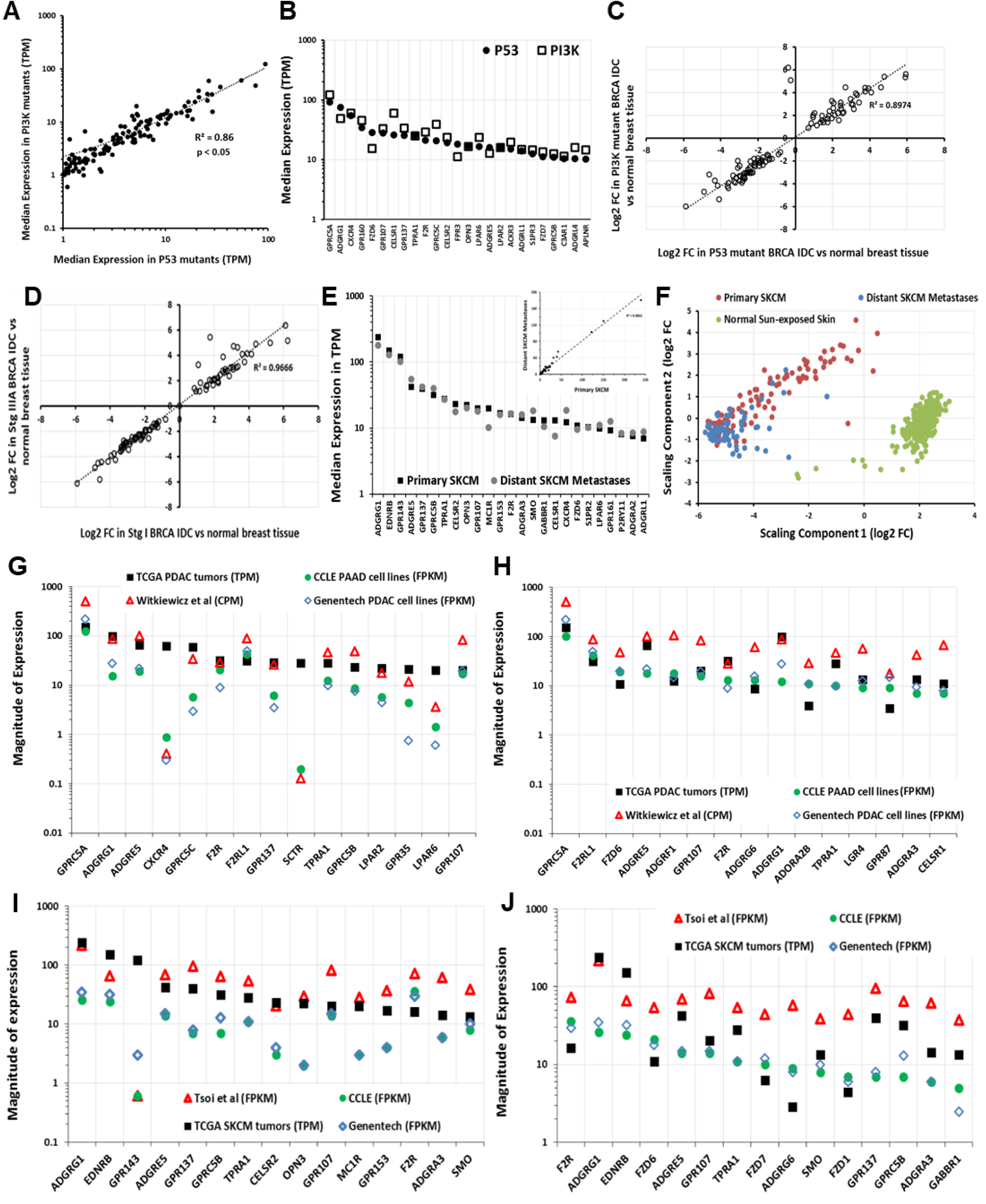
GPCR expression and presence of driver mutations and the similarity in GPCR expression of primary tumors, metastases and cancer cells derived from the tumors. (**A**) Correlation of median expression of GPCRs in P53 mutant and PI3K mutant HR-positive BRCA IDC tumors (**B**) Median expression of the 25 highest expressed GPCRs in P53-mutated tumors compared to expression in PI3K mutant HR-positive BRCA IDC tumors (**C**) Fold-changes of GPCRs in P53 mutant and PI3K mutation HR-positive BRCA IDC tumors compared to normal breast tissue. (**D**) Fold-changes of GPCRs in Stage 1 and Stage 3 ER-positive BRCA IDC tumors over normal breast tissue. (**E**) Expression of the 25 highest expressed GPCRs and (inset) correlation of median GPCR expression between primary and metastatic SKCM. (**F**) Gene expression of primary and distant metastatic SKCM tumors cluster differently and have large numbers of DE genes (**Supplement 3**) (**G**) Median expression of highest expressed GPCRs <u>in PDAC tumors</u> compared to cancer cells (CCLE [38], n = 33; Genentech [39], n = 16; Witkiewicz et al., [41], n = 72). (**H**) Median expression of highest expressed GPCRs <u>in CCLE PDAC cell lines</u> compared to other cell lines (as in panel **G**), primary cells and PDAC tumors. (**I**) Median expression of highest expressed GPCRs in <u>SKCM tumors</u> compared to cancer cells (CCLE [38], n = 45; Genentech [39], n = 44; Müller et al., [40], n = 29). (**J**) Median expression of highest expressed GPCRs in <u>CCLE SKCM cell lines</u> compared to other cell lines (as in panel **I**), primary cells and SKCM tumors.

GPCR expression appears largely independent of driver mutations, such as in BRCA HR+ IDC tumors with either *PI3KA* or *P53* mutations (**Fig 7A-C**); both groups have similar GPCR expression and DE of the same GPCRs compared to normal breast tissue. Similar results occur for LUAD and STAD that have or lack P53 mutations. Increased GPCR expression in solid tumors may thus not depend on specific driver mutations. Presence of highly overexpressed GPCRs may be a more ubiquitous feature of tumors than the presence of specific driver mutations, as exemplified by PDAC (**Fig 3B**) and in other tumor types with DE of numerous GPCRs (**Table 1**).

Moreover, GPCR expression also appears to be independent of a patient’s sex. **Figure 3F** shows this for PDAC as a representative example, with the 30 highest expressed GPCRs in males and females. This finding appears to be generalizable to other tumor types as well. Thus, the elevated expression of highly expressed GPCRs in tumors appears to occur in nearly every patient within a tumor type and is independent of attributes such as sex, tumor progression and the mutations present. As discussed below, GPCR expression tends to be independent of CNV. Each tumor type thus expresses a repertoire of GPCRs that is largely, conserved among all patients with that type of cancer. This finding suggests the potential relevance of such highly expressed GPCRs as cancer drug targets.

### GPCR expression is likely to be similar in metastatic and primary tumors

The SKCM gene expression dataset in TCGA has the most replicates of metastases. Primary and metastatic SKCM show similar expression and DE of GPCRs (e.g., *GPR143, EDNRB* and other highly expressed GPCRs) even though major differences occur in overall gene expression between primary and metastatic SKCM (**Fig 7E, F; S3F-H**). We found similar results for GPCR expression with primary and metastatic BRCA and THCA tumors and for recurrent and primary ovarian tumors (**Figs S6A-F**), though the number of replicates for each is small (<10). As new databases with large numbers of metastatic and primary tumor become available, we anticipate extending this analysis to strengthen this conclusion.

### GPCRs highly expressed in tumors are highly expressed in cancer cells

We assessed RNA-seq data for GPCR expression in cancer cell lines from the EBI portal generated via the iRAP analysis pipeline [37] for cell lines in CCLE [38] and from Genentech [39]. The use of a different analysis pipeline than that used for TCGA data does not allow direct statistical comparisons of the datasets but confirms that most GPCRs in TCGA tumors are present in cancer cells and vice versa. We also mined RNA-seq for primary melanoma cells [40] and PDAC cells [41] from the NCBI GEO database. The data from these sources (**Methods, Section 1**) allow an approximate comparison with data for tumors. **Supplement 3** shows GPCR expression in cancer cell lines.

As an example, GPCRs with the highest median expression in TCGA PDAC tumors are generally highly expressed in PAAD (pancreatic adenocarcinoma, most likely PDAC) cell lines and patient-derived PDAC cells [41] (**Fig 7G**). A few exceptions exist, perhaps from effects of cell culture or expression by non-cancer cells in tumors. Even so, highly expressed GPCRs in PAAD cells are highly expressed in PDAC tumors (**Fig 7H**), findings that also occur in other tumors, such as SKCM (**Fig 7I, J**). Thus, most highly expressed GPCRs in tumors are also highly expressed in cancer cells and vice versa.

### Most overexpressed GPCRs are rarely mutated

The most frequently mutated GPCRs in solid tumors are rarely overexpressed (**Figs 8A-B, 10H**) and conversely, highly overexpressed GPCRs in solid tumors are rarely mutated (**Fig 8A-B, Supplementary Table 5**). In SKCM, which has the highest mutation burden among TCGA tumor types, the most highly overexpressed GPCRs (*GPR143*, *EDNRB* and *GPR56*) are mutated in <2% of SKCM tumors whereas frequently mutated GPCRs (e.g., *GPR98*, mutated in nearly 40% of tumors) typically have low expression. The most frequently overexpressed GPCRs across all tumors (e.g., *FPR3*; **Table S5**) are mutated in <1% of all tumors surveyed, compared to frequently mutated GPCRs, e.g., *GPR98, GPR112*, which are mutated in >5% of all TCGA tumors surveyed. Thus, the frequency of GPCR mutations and likelihood of overexpression do not correlate (**Fig 8B**). The majority of GPCRs overexpressed in >20 tumor types/subtypes are mutated in < 50 samples out of > 5000 TCGA samples surveyed. Further, as discussed in following sections on GPCR mutation, mutations to these GPCRs are predicted to have no functional impact and are not enriched significantly for mutations at specific sites; thus, overexpressed GPCRs in tumors are not expected to be altered in their function by mutations.

**Figure 8.**
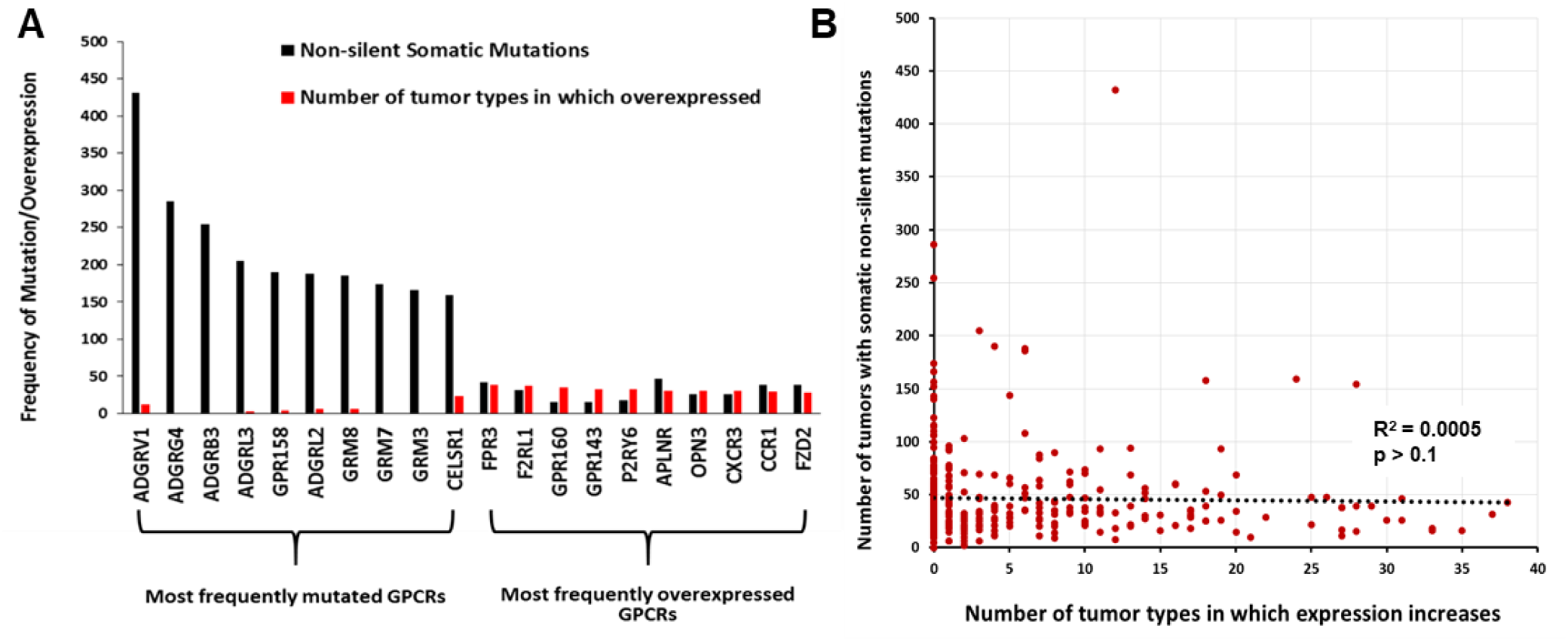
Frequent overexpression of GPCRs in tumors does not correlate with frequency of mutation. (**A**) The number of tumors with GPCR mutations and of tumor types/subtypes in which the same GPCR is overexpressed for the 10 most frequently mutated GPCRs in TCGA tumors surveyed and the 10 most frequently overexpressed GPCRs. (**B**) The frequency of overexpression compared to the frequency of mutations.

### The predicted association of expressed GPCRs with Gα G proteins

**Supplement 2** shows GPCRs annotated in the IUPHAR / BPS Guide to Pharmacology GPCR database [2], their signal transduction via G protein heterotrimers and if they are orphan GPCRs. Tissues and tumors typically express >150 GPCRs (at detection thresholds >0.1 TPM) that couple to the major types of G proteins (Gs, Gi/o, Gq/11, G12/13), most frequently Gi/Go and Gq/G11 (**Figs S6A-B**). We calculated the abundance of GPCRs that couple to each G protein by summing median GPCR expression (TPM), thereby yielding an expression ‘repertoire’ for each signaling mechanism (**Figs S6C-D**). Gs-coupled GPCRs typically account for the smallest GPCR expression repertoire for which such coupling is known. **Supplement 3** provides GPCR expression and G protein linkage data for normal tissues and solid tumors. Summing expression (TPM) of all GPCRs provides an estimate of the proportion of GPCRs among total mRNA (**Figs S6E-F**).

The GPCR expression repertoire varies among tissues in terms of total expression, number and identities of GPCRs. Tumors typically have a different GPCR repertoire than normal tissue, with increased or decreased expression of many GPCRs (**Fig 1C, Figs S6E-F**). Total GPCR expression and the number of GPCRs above expression thresholds increases in certain tumors (e.g., PDAC) but decreases in others (e.g., LIHC and SKCM) compared to normal tissue. Tumors also differ from normal tissue with respect to the abundance of GPCRs that couple to different G proteins (**Fig 1I, Figs S6C-D**), suggesting changes in signaling. For example, Gs-coupled GPCR expression decreases in many tumors (e.g., PDAC, **Fig 3C**), implying decreases in cAMP signaling.

### GPCRs as potential therapeutic targets in cancer

Among the >200 GPCRs overexpressed in at least one of the 45 tumor subtypes, 77 are targets for drugs approved by the FDA and/or EMA. (**Supplement 3**). Among these GPCR drug targets, >50% are overexpressed in 4 or more tumor subtypes (**Fig 9A**) and 15 GPCRs are increased in expression in 10 or more tumor subtypes. These results highlight the potential of GPCRs as targets in cancers and importantly, for the possible repurposing of drugs approved for other indications. Among the 77 GPCRs with increased expression in tumors, nearly two thirds link to either Gs- or Gi-coupled signaling and thus are predicted to regulate cAMP formation (**Fig 9B**).

**Figure 9.**
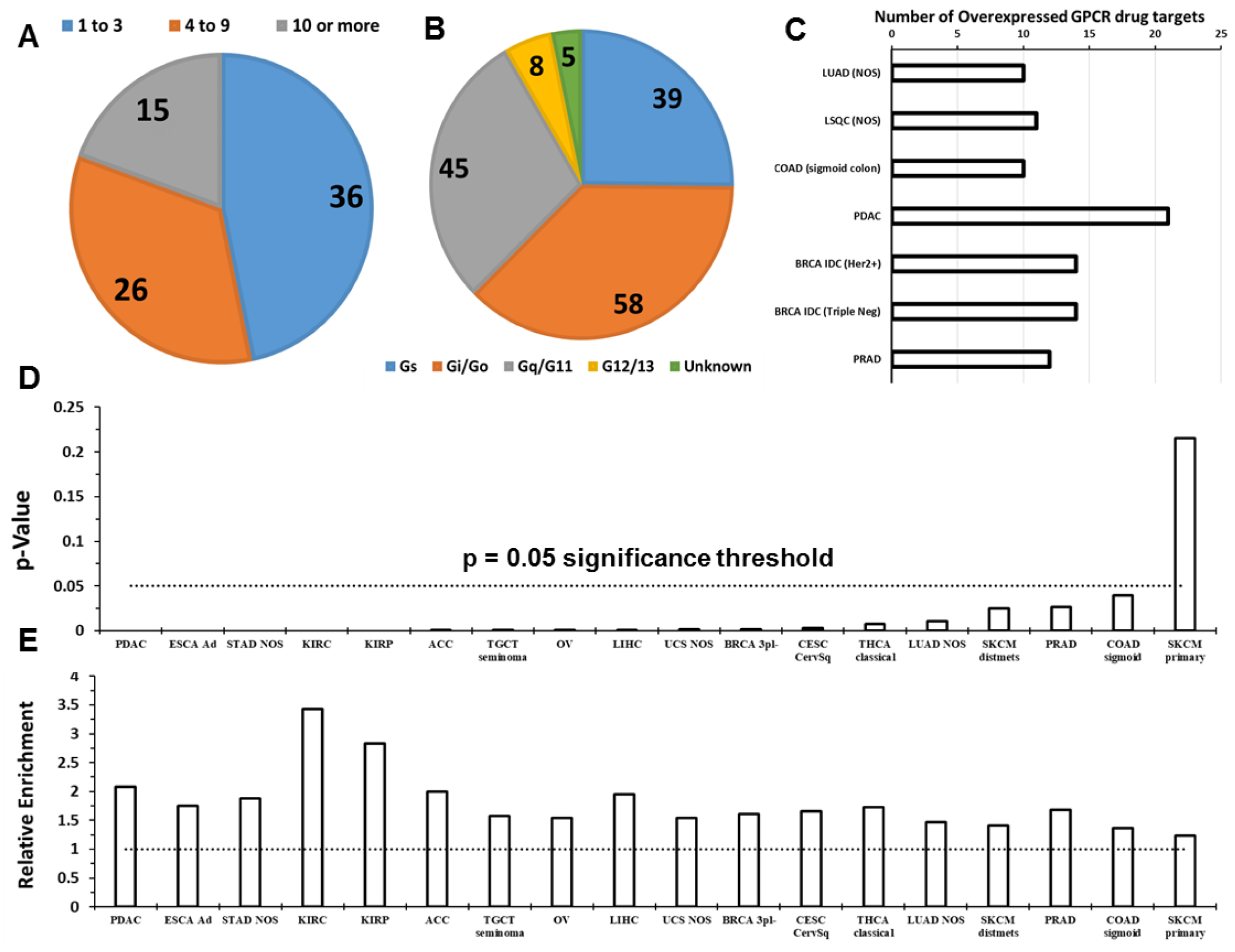
Solid tumor-expressed GPCRs with potential as drug targets. (**A**) The number of GPCRs that are targets for approved drugs and have increased expression in 1-3, 4-9 or ≥10 tumor subtypes. (**B**) The linkage to G proteins of the 77 GPCRs targeted by approved drugs and with increased expression in at least 1 tumor subtype. Note: multiple GPCRs couple to more than one G protein. (**C**) The number of GPCRs targeted by approved drugs that show increased expression in lung, colon, pancreatic, breast, and prostate cancers, the leading causes of cancer deaths in the U.S. (**D**) Overrepresentation of GPCRs among genes with >4-fold elevated expression (FDR < 0.05) for the indicated tumor types/subtypes with p-value calculated via Fischer’s exact test (**E**) The magnitude of overrepresentation (relative enrichment) of GPCRs corresponding to the p-value in (**H**)

Of the solid tumor types we analyzed, lung, colon, pancreatic, breast, and prostate cancers account for the largest annual number of deaths in the U.S. (https://www.cancer.gov/types/common-cancers). **Figure 9C** shows currently druggable GPCRs with increased expression in subtypes of those tumor types. Approved drugs target at least 10 GPCRs that have increased expression in those tumors. Hierarchical clustering of GPCR expression in different tumor types (based on their median expression of all GPCRs) reveals that GPCR expression distinguishes tumor types into groups that are consistent with other molecular/physiological traits (**Fig 5**). As discussed in sections above, GPCR expression appears to characterize categories of tumors and certain GPCRs may be targets across tumor classes/families.

As a protein-coding family of genes, GPCRs are disproportionately enriched among overexpressed genes in solid tumors, when compared to all protein-coding genes. Evidence for this was obtained as follows: In each tumor type indicated (**Fig 9G-H**), the ratio of number of coding genes with increased expression (above a prescribed threshold) over the total number of differentially expressed genes present was computed for a) GPCRs only; b) all coding genes. Fischer’s exact test was used to verify if overrepresentation of GPCRs among genes with increased expression is significant (p < 0.05). Data shown are for coding genes with >4 fold increased expression in solid tumors, i.e., highlighting genes with drastic increases in expression. We found that many tumor types have 2-fold or greater overrepresentation of GPCRs among coding genes with large increases in expression.

Moreover, extending from the data shown in fig (**6F-G**), we find that many GPCRs expressed by solid tumors may be prognostic markers. GPCR mRNA expression is associated with differences in survival in multiple tumor types (**Fig 10**). Using GPCR expression normalized in CPM, we performed survival analysis using the modified Peto-Peto test for every GPCR in tumor types where sufficient numbers of TCGA replicates were available (typically >60 replicates, allowing for >30 samples in each group for survival analysis). We omitted tumor types such as TGCT (Testicular cancer) for which almost no fatalities were recorded in the metadata. For certain tumor types with large numbers of replicates (e.g. OV, LUAD, LSQC etc), we have also provided alternative survival analysis on the related website omitting patients who are >75 years old at time of diagnosis, to reduce the risk of other sources of mortality confounding the analysis. This analysis yielded many GPCRs with a statistically significant impact on survival of each tumor type. Combined expression, i.e. a mean-normalized sum of expression, of GPCRs that individually predict survival and/or are highly differentially expressed yield stronger composite markers with prognostic relevance. **Figure 10A-C** shows examples of such combined markers in LSQC (**A**), LUAD (**B**), SKCM (**C**), which demonstrate strong adverse effects on survival. Certain tumor types, in particular KIRP, KIRC, LSQC and LUAD, express individual GPCRs that have statistically significant associations with survival (**Fig 10D**).

**Figure 10.**
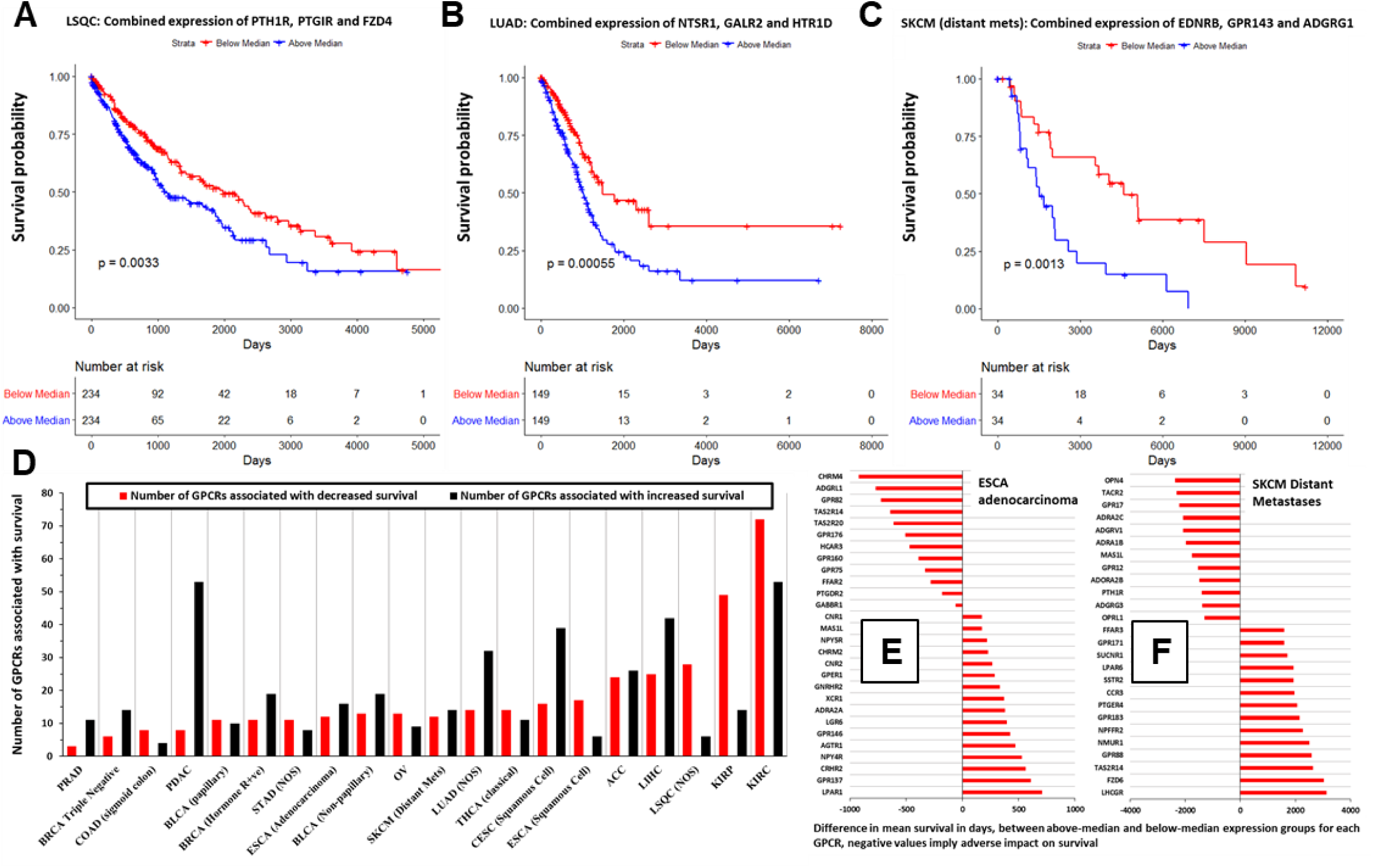
GPCR expression is associated with survival in many types of solid tumors. (**A-C**) Kaplan-Meier survival curves in the indicated tumor types, for weighted, combined expression of the indicated GPCRs (**D**) In the indicated tumor types, the number of GPCRs with significant (p < 0.05) association with survival (**E-F**) As examples, the impact on mean survival of GPCRs in (**E**) ESCA adenocarcinoma and (**F**) SKCM (distant metastases)

For two tumor types as examples (ESCA Adenocarcinoma and SKCM distant metastases, **Fig 10E-F**), we show the difference in mean survival times (in days) between patients with high expression (above median) and low expression (below median); negative values imply that elevated expression of these GPCRs is associated with adverse survival rates. GPCR expression is associated with both negative and positive survival effects, depending on the tumor type and GPCR. Several GPCRs are associated with differences in survival of >1 year. Some GPCRs have low median expression (<1 TPM) in the tumor population in general (e.g., taste receptors *TAS2R14* and *TAS2R20* in ESCA) but may be expressed in subsets of these populations, wherein they may contribute to differences in survival. Our analysis focusses on higher expressed GPCRs but we provide (at the accompanying website) expression and DE data for all GPCRs regardless of expression level, since in certain tumors very low-expressed GPCRs may have disease relevance in subsets of patients within a cancer cohort.

In total, we found that 301 GPCRs show statistically significant evidence (p <0.05) of an association with survival, in at least one tumor type/subtype (among 20 where such analysis was feasible). A subset of 32 GPCRs was significantly associated with survival in 5 or more of these tumor types/subtypes. All associations of GPCRs with survival in the tumor types tested are available at the accompanying website.

### Somatic mutations of GPCRs in solid tumors

Analysis of 5103 TCGA samples in 20 tumor types (**Table S1**; 21 tumor types if one divides ESCA into esophageal adenocarcinoma and squamous cell carcinomas) revealed many GPCRs with frequent non-silent mutations (**Fig 11A, S8A**), including a more frequently mutated subset (**Fig 11A, inset**). *GPR98*/*ADGRV1*, the most frequently mutated GPCR, occurs in >8% of TCGA samples. Tumor types with high mutation burdens have a high frequency of GPCR mutations (**Fig 11B**). SKCM has the highest frequency: ~40% SKCM tumors have *GPR98* mutations (**Fig 11C, H**). Approximately 65% of tumors have ≥1 non-silent GPCR mutation. Certain GPCRs are mutated in >10% of specific tumor types (**Fig 11C**).

**Figure 11:**
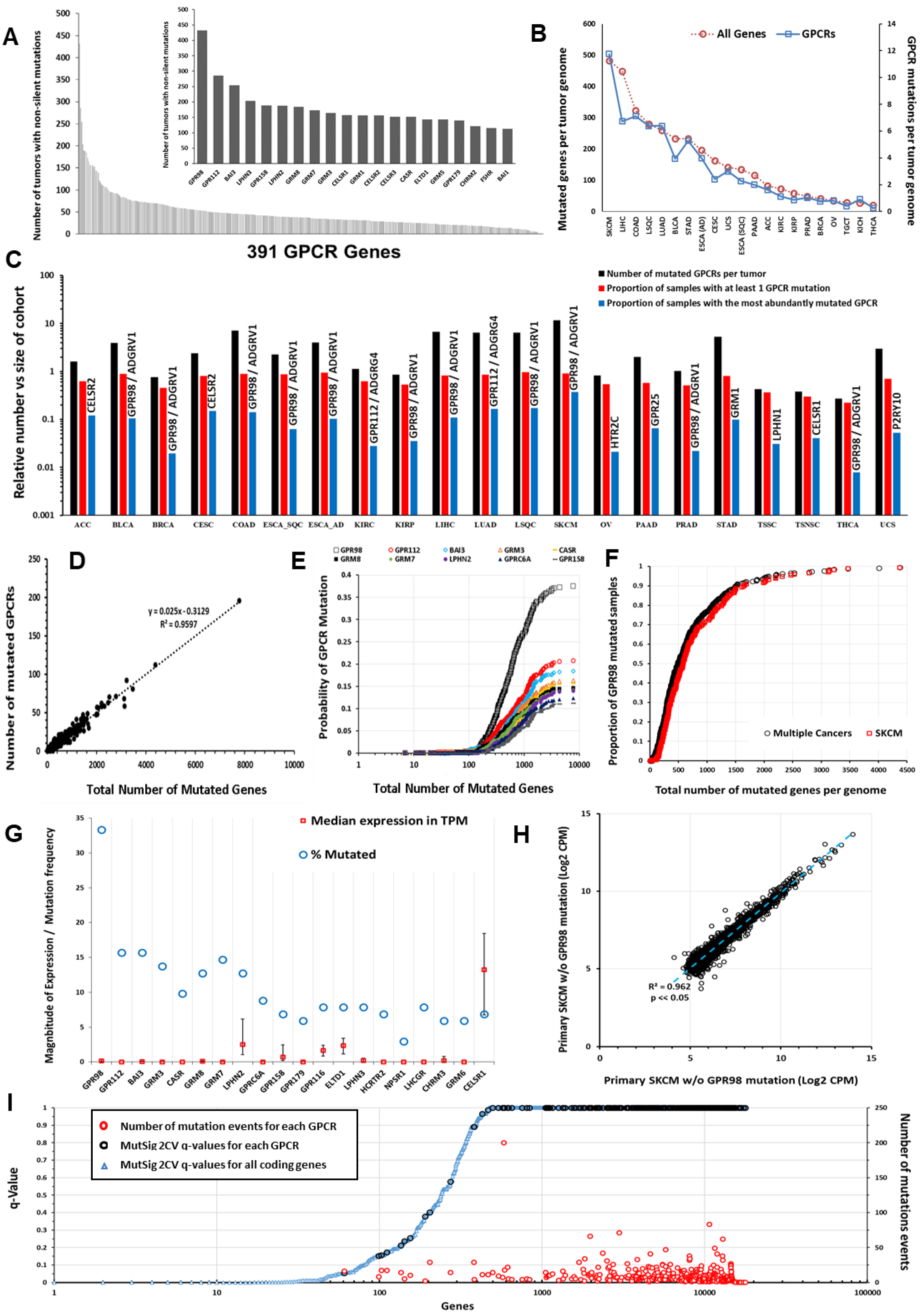
Somatic non-silent mutations of GPCRs. (**A**) Frequency of GPCR mutations in the TCGA cohort (n = 5103). Inset: 20 most frequently mutated GPCRs. (**B**) The average number of all genes (red) and GPCRs (blue) with somatic, non-silent mutations per tumor genome for the TCGA tumor types surveyed (**C**) The number of mutated GPCRs per tumor for several types of solid tumors in TCGA (black), the proportion of samples in each tumor type with at least one mutated GPCR (orange) and proportion with non-silent mutations for the most commonly mutated GPCR (gray) for each tumor type. (**D**) GPCR mutation frequency is linearly related to Nmut in SKCM. (**E**) Probability of GPCR mutation as Nmut increases in SKCM for the 10 most frequently mutated GPCRs (**F**) Normalized probability of GPR98 mutation as Nmut increases in SKCM, and the same for several cancers with high mutational burden and combined for BLCA, LUAD, LSQC, COAD, and SKCM. (**G**) The 20 most frequently mutated GPCRs in primary SKCM, with frequency of mutation and median (and upper and lower quartile) expression in TPM (**H**) Expression (CPM) of the 5000 most abundant genes in SKCM correlate closely in primary SKCM tumors that have or lack *GPR98* mutations. (**I**) MutSig 2CV v3.1 scores in SKCM obtained from https://gdac.broadinstitute.org/, showing the q-values for the significance of mutation scores for each annotated coding gene (blue); GPCR (black) and for those GPCRs, the number of mutation events (red) among the SKCM cohort.

Nmut, the number of genes with somatic non-silent mutations per tumor genome, and the number of mutated GPCRs scale linearly in individual tumors (**Figs 11D, S8E-G**). Frequently mutated GPCRs (e.g., *GPR98*, *GPR112*, *and BAI3*) are more likely to be mutated as Nmut increases (**Fig 11E**, SKCM as an example). The relationship between Nmut and likelihood of GPR98 mutation is similar in SKCM and other cancers (**Fig 11F**); this is also observed for other frequently mutated GPCRs. Hence, the likelihood of a GPCR being mutated appears to depend on the accumulation of genome damage and to be independent of the mechanisms for the mutations. The relationship between GPCR mutation rates and Nmut is identical in BLCA, LUAD and SKCM, although the factors driving DNA damage and oncogenesis are likely different. Mutations of certain GPCRs, such as GPR98, may thus serve as a bellwether for genome-wide DNA damage.

Missense mutations and in-frame deletions are the most frequent non-silent mutations in GPCR genes (**Fig S8C-D, Table S3**). Mutations in frequently mutated GPCRs occur at many sites (**Fig S9A**), which contrasts with the smaller number of such sites in common oncogenes, e.g., *KRAS* [9]. (**Fig S9B**). Certain GPCR genes (e.g., *GPR98*) may be in genomic regions vulnerable to dysregulation of DNA damage and repair and belong to a subset of mutated genes; *GPR98* mutations frequently occur alongside other frequently mutated genes such as *TTN* and *MUC16* (**Fig S10A-G**). *GPR98* is among the 25 most frequently mutated genes in all tumor types surveyed; its mutational frequency is similar to that of genes (e.g., *BAGE2* [**Fig S8B**]) that are mutational hotspots [42]. As the GPCR with the largest gene length (~19,000 bp), *GPR98* has more mutational events. Compared with other very long genes, e.g., genes > 10,000 bp, *GPR98* belongs to a subset of ~20 genes with high mutational frequencies (**Fig 12A**), implicating *GPR98* as a hotspot for both silent and non-silent somatic mutations. *GPR98* has a >4-fold increased density of mutational events (normalized for gene length) compared to the average of these very long genes. One obtains a similar result by assessing the density of somatic mutations across all genes, irrespective of length (**Fig 12B**). GPR98, GPR112 and BAI3 are among the top 5% of genes in number of mutations per unit gene length, highlighting these genes as chromosomal regions with higher than normal rates of somatic mutation.

**Figure 12:**
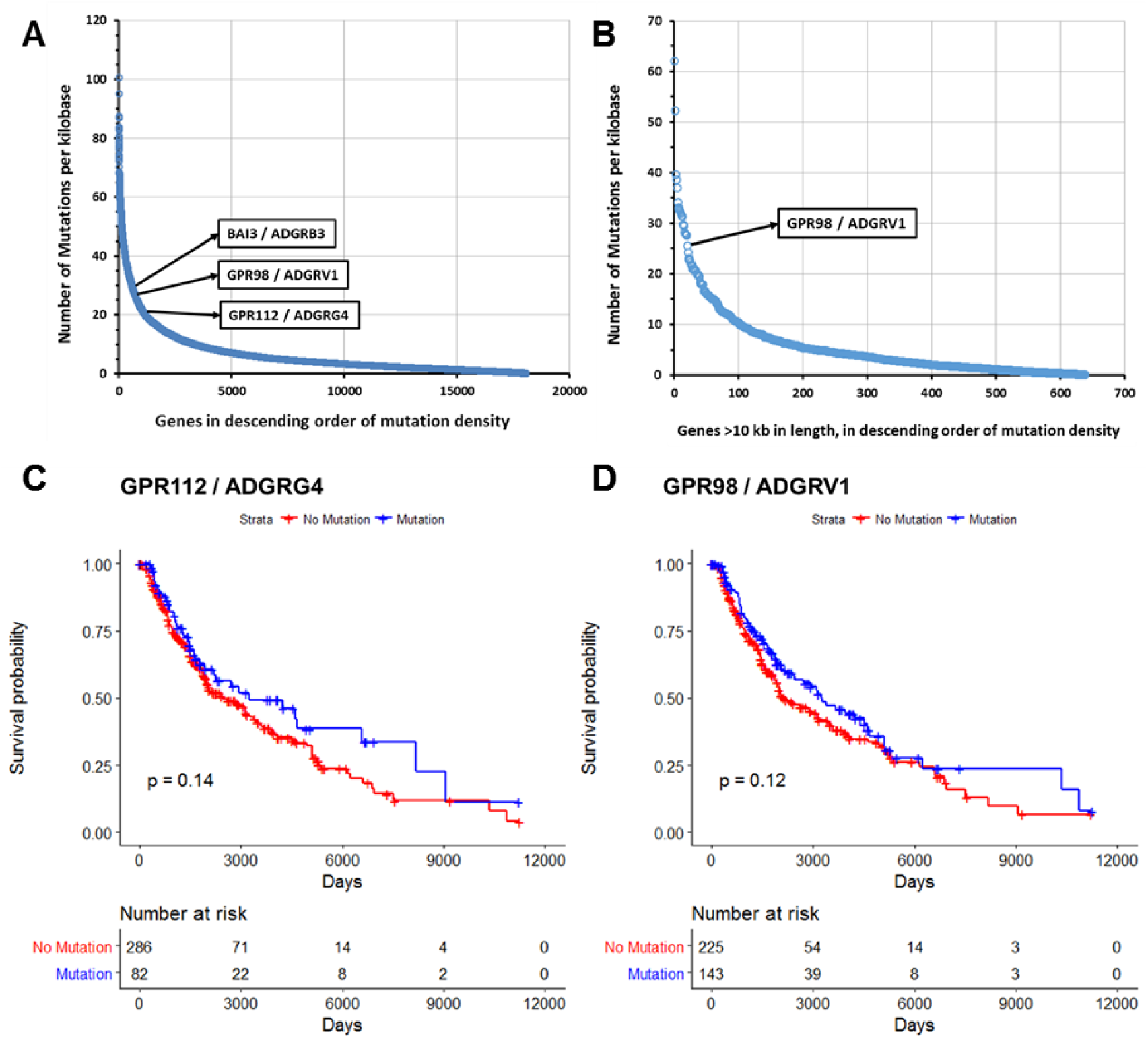
Mutated GPCRs show high density of mutations but do not impact survival. (**A-B**) Number of somatic (silent and non-silent) mutations per unit of gene length (total length of all exons per genes) for (**A**) all genes and (**B**) genes >10 kb in length in SKCM. (**C-D**) Survival analysis in metastatic SKCM for (**C**) GPR112 and (**D**) GPR98 mutations, with p-values calculated using the Peto & Peto modification of the Gehan-Wilcoxon test.

Survival analysis of metastatic SKCM samples was performed in order to evaluate the impact on tumors of somatic non-silent mutations to GPR98, GPR112, or other frequently mutated GPCRs. Analysis of Kaplan-Meier survival curves using the modified Peto-Peto method reveals that presence of somatic mutations in these GPCRs does not have a statistically significant impact on survival. **Fig 12C-D** shows this for GPR98 and GPR112, the two most frequently mutated GPCRs in SKCM. We find the same result in other tumor types as well and thus conclude that somatic non-silent mutations to GPCRs have no impact on patient survival.

Most mutated GPCR genes have low levels of mRNA expression (**Fig 8; 11G** [**SKCM as an example**]) so may not be functionally relevant but certain GPCR genes (e.g., *CELSR1* and *LPHN2*/*ADGRL2*) are frequently mutated and moderately/highly expressed. Several such GPCRs are orphan receptors (without known physiologic agonists or roles), for which it is unclear if they impact on cell function. As cell-surface receptors, frequently mutated, well expressed GPCRs may represent neo-antigens. For SKCM, which has the most GPCR mutations among tumors types surveyed, DE analysis of primary melanomas and distant metastases that have or lack GPCR mutations (e.g., *GPR98* and *LPHN2*) revealed little evidence that these mutations alter the tumor transcriptome, implying that such GPCR mutations are likely passenger, rather than driver, mutations (**Fig 11H**; **Fig S8C, D**). Conversely, previous work has suggested that for known oncogenes (e.g., for TP53, [43]), there are often widespread transcriptomic changes associated with specific mutations. We found similar behavior for other tumors (e.g., BLCA) that have frequent GPCR mutations.

As a further approach, we evaluated GPCR mutations, predicting the likelihood of functional consequences and site-specific enrichment of the mutations via MutSig 2CV v3.1 (*gdac.broadinstitute.org*). The majority of GPCRs frequently mutated (**Fig 11-I**, SKCM as example) show non-silent mutations that are non-significant in terms of enrichment (compared to the background mutation rate of silent mutations over the same regions) for individual mutation sites. These mutations are not predicted to be functional (calculated from estimations of functional impact of mutations based on whether mutated regions are highly evolutionarily conserved) by MutSig 2CV, consistent with the idea that the frequent GPCR mutations are likely passenger and not driver mutations. MutSig 2CV results for all GPCRs in all tumor types are available at the accompanying website.

### Copy-number variation (CNV) of GPCRs in solid tumors

CNV of certain GPCRs occurs frequently in TCGA solid tumor samples (**Fig 13A-B; Table S1**), with some GPCRs (e.g., *GPR160*) amplified in >5% of all TCGA samples surveyed. CNV data were obtained as GISTIC 2.0 [44] thresholded data, wherein values of −2, −1, 0, 1 and 2 respectively denote homozygous deletion, single copy deletion, diploid copy number, low level amplification (i.e. increase of 0.1 to 0.9 of copy number, expressed as a log2 ratio) and high-level amplification (amplification of >0.9 of the log2 ratio, i.e. >1.7 extra copies in a diploid cell) [45]. The distribution of amplification events among GPCRs parallels that of all genes (**Fig 13C**, SKCM as an example) but a subset of GPCRs is disproportionally amplified (**Fig 13D-F, and Supplement 3**). Most GPCRs have infrequent amplification (in 2% of tumors or less). Amplification does not predict high expression or overexpression of GPCRs (**Fig 13G-J**; OV as an example): frequently amplified GPCRs in tumors often have limited mRNA expression in those tumors while most highly expressed, overexpressed GPCRs are not amplified.

**Figure 13.**
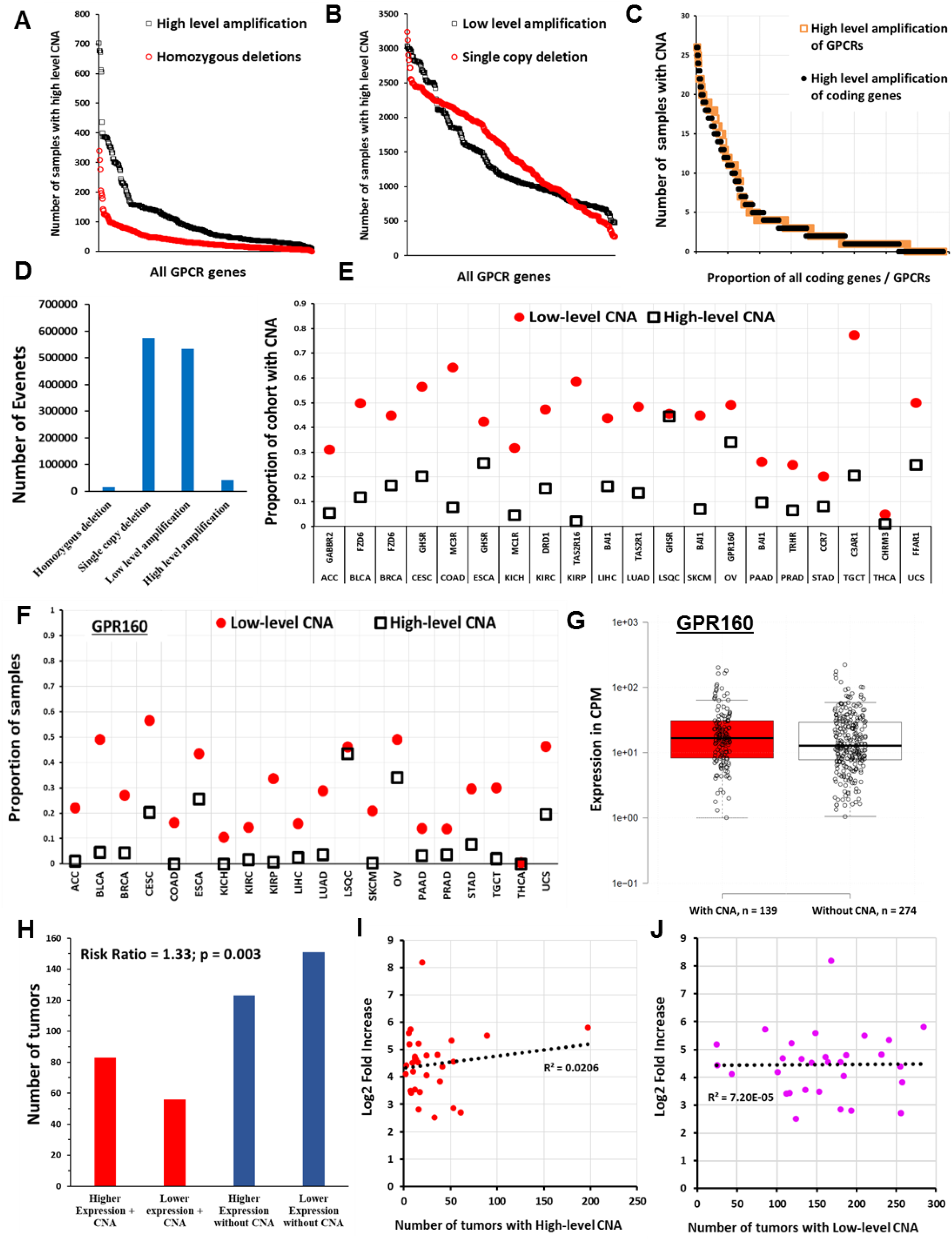
Copy number variations (CNVs) of GPCRs in solid tumors. (**A**) The number of solid tumors with CNV for each GPCR across all TCGA samples (plotted in descending order of frequency for high-level amplification and homozygous deletions; see text for definition of high- and low-level amplification) (**B**) The same as in (**A**) for low-level amplification and single-copy deletions (**C**) In SKCM tumors (n = 367), the distribution of high-level amplification of GPCRs (n = 390 genes) compared to that of all protein coding genes (n = 24,776) (**D**) The total number of homozygous deletions, single copy deletions, low-level and high-level amplifications for all GPCRs combined in the 7545 TCGA tumors surveyed for CNV (**E**) The most frequently amplified GPCR for each TCGA tumor type and the proportion of samples with High and Low-level amplification for that GPCR (**F**) Proportion of samples of various tumors types with High and Low-level amplification of *GPR160*, the most frequently amplified GPCR overall) (**G**) Ovarian cancer samples with and without High-level amplification of GPR160, with the median for each group indicated. The difference between groups was not statistically significant) (**H**) The risk ratio of elevated GPR160 expression (above the median value for OV) when GPR160 also shows high-level amplification; amplification of GPR160 increases the likelihood that GPR160 expression is elevated (**I**) For the 30 GPCRs in OV with the highest fold-increase in expression relative to normal ovarian tissue (with FDR < 0.05 and median expression in OV > 1TPM), the corresponding number (of 579) OV tumors with high-level amplification of those same GPCRs (**J**) The same as (**I**), but comparing fold-increase against low-level amplification

Single-copy/heterozygous deletions of GPCRs are widespread, whereas homozygous deletions are rare (**Fig 13A, D**). GPCR genes with single-copy deletions are generally not significantly expressed in tumors or normal tissues, implying that such deletions lack functional effects but exceptions exist. *PTH1R*, frequently deleted in KIRC (~77% samples have single-copy deletions), has ~10-fold reduced expression compared to normal kidney tissue. Similarly, *ADRA1A* is frequently deleted and has reduced expression in hepatocellular and prostate adenocarcinomas.

**Figure 13E** shows the identity/frequency of amplification of the most frequently amplified GPCR in each tumor type. Several cancers (e.g., OV and LSQC) have a high level of amplification of specific GPCRs in >25% and low-level amplification in >40% of samples. **Figure 13F** shows the CNA frequency of *GPR160*, the most frequently amplified GPCR overall, among all tumors surveyed. Except for *GPR160* and *FZD6*, most GPCRs with frequent amplification are rarely overexpressed (**Supplement 2**). CNA alone thus does not generally predict increased mRNA expression in tumors compared to normal tissue and highly expressed GPCRs in tumors are typically not amplified. We tested for correlation between high-level amplification and increased mRNA expression. **Figure 13G** shows GPR160 expression in OV tumors with and without high-level amplification; CNA is not a prerequisite for high mRNA expression and the small difference in median expression is not statistically significant, based on DE analysis via edgeR. However, tumors with amplification of GPR160 show a higher likelihood (~33%, **Fig 13H**) of expressing GPR160 at levels above the median for OV. We did not observe statistically significant risk ratios relating GPCR expression with amplification for other frequently amplified GPCRs. We conclude that CNA and GPCR mRNA expression are generally poorly correlated; hence, examination of amplified GPCR genes does not predict which GPCRs are highly and/or differentially expressed in a tumor (**Fig 13I, J**).

**Supplements 2 and 3** provide, respectively, the frequency of GPCR CNV and changes in expression of GPCRs in the tumors surveyed. The widespread CNV of certain GPCRs suggests that they (and/or neighboring genes that vary along with these GPCRs) contribute to the malignant phenotype and may be markers for malignancy.

## Discussion and Conclusions

In this study, we identified mutations, CNVs and alterations in mRNA expression of GPCRs in a range of solid tumors. The results reveal a broad landscape of changes, suggesting a functional role for this gene superfamily in such tumors and possible therapeutic utility of GPCRs in a large number of solid tumors.

Mutations of certain GPCRs have been implicated in cancer [8], but a comprehensive analysis of GPCR amplification, expression and DE has been lacking. The largest public datasets of normal (GTEx) [1] and cancer tissues (TCGA) provide RNA-seq data analyzed/normalized differently, making it difficult to compare datasets. For many tumors types, few replicates of ‘normal’ TCGA tissue are available and samples from matched non-tumor tissue from patients may not be representative of normal tissue (**Supp. Methods, Section 1**). The TOIL project enabled comparison of TCGA and GTEx data with analysis of both RNA-seq datasets via the same pipeline.

Our analysis identified frequently mutated GPCRs (e.g., *GPR98/ADGRV1* and *GPR112/ADGRG4*) in multiple cancers, especially melanoma (SKCM). GPCR mutations appear to reflect accumulation of DNA damage and mutations across the genome and may be tumor markers for this process. Multiple prior studies (e.g. [8,46]) have suggested that GPCRs that are frequently mutated (e.g., certain adhesion GPCRs) in tumors should be further studied for their role as drivers of tumor progression and as targets for novel therapeutics. Several observations lead us to question this idea:

1. The absence of enrichment of mutations in specific sites/regions on frequently mutated GPCRs
2. The analysis from MutSig2CV indicating the likelihood that such mutations are not enriched at specific residues at rates statistically elevated above the background mutation rate, i.e. the rate of non-synonymous mutations is not statistically elevated, adjusted for the rate of synonymous mutations.
3. The general low expression of such mutated GPCRs at the mRNA level
4. The lack of impact of GPCR mutations on survival
5. The lack of effect of GPCR mutations on gene expression

While GPCRs are of interest in cancer due to their high mRNA expression, using their frequency of mutation as a rationale to choose particular GPCRs as potentially functionally important or possible therapeutic targets is likely a flawed approach—a conclusion that contrasts with previous suggestions about mutated GPCRs. Our data also raise a cautionary point about mutations: frequency of mutation of a gene is insufficient evidence to suggest its importance as a potential oncogene or therapeutic target without additional analyses such as those noted above.

While we conclude that frequently mutated GPCRs are unlikely to be targets for therapeutic intervention in solid tumors, such GPCR genes may be useful for studies related to oncogenesis. For example, GPR98 is frequently mutated in a number of tumor types and is one of several genes with a high frequency of somatic mutations per kilobase of coding gene length, suggesting that the GPR98 gene lies on a chromosomal region particularly vulnerable to accumulation of DNA damage. Mutation to this gene seems to scale with mutational burden, in a manner that is independent of tumor type: in tumors such as melanoma and bladder cancer, in which DNA damage from environmental factors may occur via different mechanisms, the accumulation of damage at the GPR98 locus in response to genome-wide damage appears to occur at a constant rate. This rate is elevated relative to the rest of the coding genome in terms of mutation rate per kilobase. Understanding how and why this occurs may provide additional insight into the mechanisms that drive DNA damage.

Known driver mutations do not appear to influence GPCR expression in tumors but we excluded rare mutations. In order to ensure large numbers of replicates and high statistical significance, we analyzed tumors with high-frequency mutations (e.g., *P53* or *KRAS*) [9]. Highly expressed GPCRs are widely expressed among replicates of specific tumor types and are more prevalent in tumors than are common driver mutations. The overrepresentation of GPCRs among protein-coding genes with increased expression in solid tumors supports the hypothesis that the elevated expression of specific GPCRs is a hitherto underappreciated feature of solid tumors.

The finding that many GPCRs show altered mRNA expression in tumors at rates higher than occurs for coding genes in general raises the question: what causes GPCR-specific changes in expression? Regulation of GPCR mRNA expression is poorly understood; hence, exploration of potential mechanisms for such regulation/dysregulation is of interest. Studies of GPCRs with DE in tumors may shed light on such mechanisms and perhaps also have relevance for other disease settings with altered GPCR expression.

Given the widespread changes in copy number that occur in cancer, especially copy number amplification, which might explain the increased expression of particular GPCRs in tumors, we tested but failed to find that CNV can generally explain altered GPCR mRNA expression or DE. CNV is not stochastically distributed among the GPCR family, certain GPCRs are more frequently amplified (e.g., *GPR160*) or deleted (e.g., *PTH1R* in KIRC). Amplified and deleted GPCRs may have potential as biomarkers [47]. Our findings with respect to CNV and GPCRs raise a broader concern and cautionary note: changes in copy number of certain genes should not be taken as evidence of dysregulation of the same genes, unless one obtains evidence for corresponding changes at the mRNA level.

Numerous solid tumors have increased mRNA expression of large numbers of mostly non-mutated GPCRs: 72 GPCRs are overexpressed in >10 tumor subtypes, implying that common mechanisms may regulate GPCR expression in such tumors. Highly overexpressed GPCRs are potential candidates as drug targets. Of note, 77 such overexpressed GPCRs are targets of approved drugs that have the potential to be repurposed to treat tumors. The similarity of GPCR expression in primary tumors and metastases supports such therapeutic potential.

The data reveal that clusters of GPCRs may be prognostic indicators for survival and provide a molecular signature of the malignant phenotype. GPCRs whose expression adversely or positively predicts survival are candidates for antagonists or agonists, respectively as novel cancer drugs. Hierarchical clustering of tumor types based on GPCR expression identifies groups of tumors consistent with other molecular/phenotypic features of these tumors. Thus, the tumor GPCRome appears to be predictive of the broader molecular landscape of tumors.

Do GPCR mRNA data predict protein expression? Direct quantification of GPCR proteins is challenging, due to their generally low abundance and paucity of well-validated antibodies. However, mRNA expression of GPCRs, especially highly expressed GPCRs, generally predicts the presence of functionally active GPCRs in human and animal cells [48–52]. In contrast to earlier ideas, recent evidence supports the view that mRNA expression broadly predicts protein expression [53–57] (**Supplemental Note 1**). As an example, GPRC5a protein and mRNA abundance are concordant (**Supplemental Note 2; Fig S11**). GPCR detection via mass-spectrometry has been challenging; proteomics data (e.g., [58] indicate that at present, few GPCRs are detectable by such methods, likely due to the low abundance of GPCR proteins. As noted in **Results**, functional evidence is available for numerous GPCRs with DE in solid tumors. As cell-surface proteins enriched in tumors and cancer cells, certain GPCRs may represent novel tumor-associated proteins that might be targeted for diagnosis and/or treatment.

As a caveat to those ideas regarding mRNA and proteins expression, most analyses on their concordance cited above have used model organisms. Such concordance may be less evident in native mammalian cells, particularly in certain cell and tissue types, as observed in [54]. It is unclear whether GPCR mRNA expression predicts protein expression and functional effects in native cells/tissues. However, our recent work on Gq-coupled GPCRs in pancreatic cancer cells indicates a concentration-response relationship between GPCR mRNA expression and function, such that high GPCR expression corresponds to strong signaling and functional responses [59]. Previous work on GPCR mRNA expression in native cells has shown robust functional effects and/or protein expression in a range of settings, including cancer, for GPCRs identified as highly expressed, based on omics methods (reviewed in [50]). We thus suggest that highly expressed GPCRs (e.g. those expressed at ≥ 5-10 TPM from RNA-seq data) are very likely to be functional.

GPCR mutations, CNV and DE thus occur at a high frequency in solid tumors. Therefore, this receptor super-family may have unappreciated functional roles in such tumors, especially since GPCR expression appears to be largely independent of tumor grade/stage and mutations. Our results imply that new insights may derive from further studies of GPCRs regarding mechanisms of gene expression and phenotype in solid tumors and perhaps other cancers. Of particular and perhaps rapid, translational importance is the potential of GPCR-targeted drugs, including FDA/EMA-approved drugs that might be repurposed as therapeutics for a variety of solid tumors.

## Acknowledgements

This work was supported by the American Society of Pharmacology and Experimental Therapeutics (ASPET) David Lehr Award and Padres Pedal the Cause #PTC2017.

Results are in part based upon data from the Cancer Genome Atlas (TCGA) Research Network: http://cancergenome.nih.gov/.

## Author Contributions

KS and PAI conceived the project. KS designed and implemented the analysis approach and co-wrote the manuscript. PAI directed the project, interpreted results and co-wrote the manuscript. KM implemented analysis methods and maintained datasets/databases and the project website. RC co-wrote and edited the manuscript. HC interpreted results, designed analysis approach and co-wrote the manuscript.

## Supplemental Information

Supplement 1: Supplementary Figures 1-13, Supplementary Tables 1-5 and Supplementary Notes 1-3.

Supplement 2: Data file with GPCRs annotated by GtoPdb plus GPCR mutations and CNV

Supplement 3: Data file with GPCR expression in normal tissue, tumors and cancer cells and GPCRs with DE Additional downloadable content: *<u>insellab.github.io</u>*

## Methods

Software used and availability of data.

Software used and availability of data
| RESOURCE | SOURCE | LOCATION OF DATA |
| --- | --- | --- |
| **Software and Algorithms** |  |  |
| R v 3.3.2 | The Comprehensive R Archive Network | https://cran.r-project.org/ |
| edgeR | Bioconductor | http://bioconductor.org/packages/edgeR |
| **Deposited Data** |  |  |
| Numbers of GPCR mutations in each tumor type | This paper | Supplement 2 |
| Occurrence of CNV for GPCRs in each tumor type | This paper | Supplement 2 |
| GPCR expression and G protein linkages in tumors and healthy tissue | This paper | Supplement 1 |
| GPCRs with DE in tumors | This paper | Supplement 1 |
| GPCR expression in cancer cells | This paper | Supplement 1 |
| GPCRs annotated by IUPHAR | IUPHAR/BJP | Supplement 2 |
| Analyzed TCGA mutation data | TCGA/UCSC Xena | https://insellab.github.io/gpcr_mutations |
| Analyzed TCGA CNV data | TCGA/UCSC Xena | https://insellab.github.io/gpcr_cnv |
| Analyzed TCGA RNAseq data | TCGA/UCSC Xena | https://insellab.github.io/gpcr_tcga_exp |
| Analyzed GTEX RNAseq data | GTEx/UCSC Xena | https://insellab.github.io/gpcr_gtex_exp |
| Analyzed RNAseq data from cancer cells | Various cited sources | https://insellab.github.io/gpcr_cells_exp |

### 1. Contact for Resource Sharing

A website has been created for sharing all data at https://insellab.github.io/. Links to data files will be posted on this website following peer review. All data provided will be open access following peer review; information about how to cite data generated from this study is available at https://insellab.github.io/.

Contact for access to data:

Krishna Sriram

Department of Pharmacology, UC San Diego.

Tel: 858-534-2298

### 2. Details of Methods

#### 2.1. Differential Expression (DE) Analysis

Gene expression for the GTEX and TCGA datasets, assayed via RNAseq, was downloaded from the UCSC Xena Portal (xena.ucsc.edu). For DE analysis, RSEM expected counts were obtained, which were computed via the TOIL pipeline, described in [10] and available at the Xena Portal by the authors of the TOIL project (https://xenabrowser.net/datapages/?host=https://toil.xenahubs.net).

The analyzed data from the TOIL project were generated as follows. Merged FASTQ files were adapter-trimmed via CUTADAPT, followed by alignment via STAR [60]. Gene expression was then quantified using RSEM [61]. The HG38 reference genome, with Gencode V23 annotations, was used in the TOIL analysis. For this study, both RSEM estimated counts (for DE analysis) and RSEM TPMs (for evaluating magnitudes of expression) were used.

Files were accessed from (https://xenabrowser.net/datapages/?dataset=tcga_gene_expected_count&host=https://toil.xenahubs.net) for TCGA expected counts (version 2016-09-01) and (https://xenabrowser.net/datapages/?dataset=gtex_gene_expected_count&host=https://toil.xenahubs.net) for GTEX expected counts (version 2016-05-19).

Expression in TPMs for GPCRs was queried via https://xenabrowser.net/heatmap/ for both TCGA and GTEX.

Following the download of gene expression data, corresponding files on sample phenotype were obtained from the relevant links hosted at https://xenabrowser.net/datapages/?host=https://tcga.xenahubs.net for TCGA and at https://xenabrowser.net/datapages/?cohort=GTEX for GTEX samples, respectively. Samples were then grouped for DE analysis based on attributes such as tissue type and tumor type.

We used a table of estimated counts from tumor and normal tissue as input for analysis in edgeR [62], yielding normalized abundance in CPM (using TMM normalization) and DE, showing magnitude of fold change and statistical significance estimated by FDR (False Discovery Rate). We estimated fold-changes of gene expression in tumors compared to normal tissue via an exact test. Genes that changed with FDR <0.05 were considered statistically significant, however we focused attention on GPCRs that besides low FDRs are also expressed at >1 TPM in tumors, as high expressed GPCRs are likely of greater interest. The 20 TCGA tumor types were divided into 45 subtypes/categories (**Table 1**) based on histological classification. Different tumor subtypes show distinct GPCR expression, e.g., Classical vs. Follicular Thyroid Cancer (THCA), Triple negative vs. Her2 Breast Cancer Cancer, Infiltrating Ductal Carcinoma (BRCA IDC), and Esophageal (ESCA) squamous cell carcinoma vs. Adenocarcinoma (**Fig S1A-F**), hence requiring this subdivision into tumor subtypes.

In addition to the standard TMM approach in edgeR, we tested upper-quartile normalization before conducting DE analysis in edgeR. The two methods yielded nearly identical results (**Fig S11C, D).** Log2 fold-changes for all genes and GPCRs show that expression changes calculated by both methods are closely correlated, with nearly identical magnitude. We also evaluated DE via EBseq [63]. EBseq and edgeR yielded very similar results (**Fig S11A, B**), in particular for GPCRs, implying that assumptions implicit in the DE analysis via edgeR/TMM normalization do not skew or bias the results. As an empirical test, we verified that GPCRs that show large fold-changes between tumor and normal samples also show large differences in normalized gene expression in TPM, e.g., *EDNRB*, *GPR143* and *ADGRG1* in SKCM (**Fig S2F-H**).

#### 2.2. Database of normalized GPCR expression in tumors and normal tissue

For GPCR expression in tissue, expression as TPM (Transcripts Per Million, a normalization of gene abundance that corrects for effective length of genes) and CPM (Counts Per Million, number of times a gene is encountered per million reads, hence normalization for library size without length normalization) are provided in **Supplement 3**.

Data in TPM are provided for assessing the relative abundance of members of a gene family, such as GPCRs, *within* an individual sample or a set of biological replicates while data in CPM are provided for comparing a specific gene *between* multiple groups of dissimilar samples, where length normalization is problematic because dissimilar data sets may be normalized differently. We provide GPCR expression in both formats to facilitate different approaches for analysis.

This resource thus enables an estimate of GPCR abundance in more rigorous terms than comparing FPKM/TPM values for a particular gene across different tissue types. Further, this approach provides normalized gene expression estimates (in TPM and CPM) using the same units and analysis methods for both normal and cancer tissue, thus allowing for direct comparison.

Gene abundances in CPM were calculated via EdgeR. In several cases the same normal tissue dataset was used to compare multiple tumors (e.g., GTEX Kidney data was used for comparison with KIRP, KICH and KIRC). In these cases, the calculated CPM values for normal tissue from each analysis were (as expected) very similar but not equal, as EdgeR TMM normalization yields slightly different normalization factors in each case. The CPM values presented for these tissues are thus average values (e.g., for CPMs in normal Kidney tissue, the values provided are the average of the data obtained from comparisons of normal kidney with KIRP, KICH and KIRC respectively). The normal tissue types where this was performed are breast, lung and kidney.

#### 2.3. GPCR mutation and copy number analysis

For each cancer type, tables of somatic, non-silent mutations (gene-level) and somatic mutations (SNPs and small INDELs) and Gene-level GISTIC2 thresholded copy number variation were obtained using https://xenabrowser.net/datapages/?host=https://tcga.xenahubs.net and links within.

For mutation data, we used results obtained via the Broad Automated Pipeline, where available. In other cases, we used data from the Baylor College of Medicine sequencing center. The source of the mutation data is indicated on the respective downloadable files and **Table S1**. In all cases, the HG19 reference genome was used for calling mutations. Mutation data for genes coding for GPCRs were extracted as part of the present study; all GPCR mutation data are available for download as supplemental material.

TCGA copy number estimates were obtained using Affymetrix SNP 6.0 arrays. The data were analyzed via GISTIC 2.0 [44] to obtain gene-level estimates of copy number variation. The resulting ‘thresholded’ GISTIC 2.0 data yields values of −2,−1, 0, 1, 2, indicating homozygous/2 copy deletion, heterozygous/single copy deletions, no change, low level amplification, and high level amplification, respectively. Copy number variation for GPCR genes in each tumor type was extracted and is available as downloadable material.

#### 2.4. Which genes are included in this analysis?

We evaluated all GPCRs annotated by IUPHAR/*British Journal of Pharmacology* [2], accessible via (http://www.guidetopharmacology.org/GRAC/ReceptorFamiliesForward?type=GPCR). This list primarily focuses on endoGPCRs (GPCRs natively expressed in peripheral tissue, possess endogenous ligands and receptors primarily used as drug targets). The IUPHAR list includes taste and vision GPCRs but not olfactory receptors. We excluded several GPCRs annotated by IUPHAR but which are thought to be pseudogenes. The list of GPCRs in this analysis is provided in **Supplement 2**.

For these annotated GPCRs, we included information about their linkages to G proteins and their status as orphans or not. For non-orphans, an example of an endogenous ligand is provided. These data are almost entirely based on information available at the IUPHAR website cited above. In a few cases where such information is not provided by IUPHAR, we have used other literature sources for annotation.

#### 2.5 GPCR expression in cancer cells mined from other sources

GPCR expression in a range of cancer cell lines was queried via the EBI Expression Atlas (https://www.ebi.ac.uk/gxa/home) for cell line profiles part of CCLE [38] as well as by Genentech [39], profiled via RNA-seq. These data were analyzed as part of the EBI Expression Atlas via the iRAP bioinformatics analysis pipeline, described in detail in [37] wherein gene expression was computed in FPKM (Fragments Per Kilobase of exon, per Million reads), a length-normalized expression abundance estimate analogous to TPM units, used for TOIL TCGA data. Precisely, statistically relevant comparisons between gene abundances in TPM and FPKM are not feasible; however, empirical comparisons between such data sets are possible. In general, genes with high abundances in TPM or FPKM will be highly expressed relative to other genes within sets of samples; hence, our comparison of CCLE and other cell-based data vs. TOIL TCGA data serves as an empirical confirmation of the fact that GPCRs highly expressed in TCGA tumors are also present in cancer cells.

Normalized gene expression in cancer cells from other sources [40,41,64] were obtained via NCBI GEO, wherein analyzed RNA-seq data with quantification of gene expression were provided. As with the data from EBI above, such data allowed for empirical comparisons vs. TCGA TOIL data to confirm the presence of GPCRs in cancer cells, which were also detected in tumors.

#### 3. Quantification and Statistical Analysis

DE analysis was performed in the R software environment via EdgeR [62], as discussed above in section 2.1. We used the following criteria to evaluate GPCRs with significant DE:

1. FDR < 0.05. In the majority of cases, genes with a high fold change also showed FDRs << 0.05.
2. Magnitude of fold change > 2 fold (increase or decrease).
3. Magnitude of expression > 1 TPM median expression in tumors, as calculated by RSEM in the TOIL pipeline, discussed in section 2 above. We focused on genes with significant DE and high expression because our primary goal was to identify GPCRs that may be drug targets and/or biomarkers.

For compilation and distribution of data we assembled data files primarily in Microsoft Excel, with files stored in .xlsb format.

Plots of normalized expression in tumors and normal tissue, whether in TPM or CPM, show median expression for respective cohorts, along with upper and lower quartiles, as indicated in figure legends where applicable.

The numbers of replicates in each sample group/category of normal tissue and tumors are provided in **Tables S1 and S2 and Table 1**, respectively. Tumor types with 10 or more replicates were considered for DE analysis. A small number of samples (<< 1% of TCGA samples studied) were excluded because they were duplicated in downloaded databases from TOIL and showed discrepancies between expression in these downloaded data vs expression data queried via the visualization tool on the TOIL website. These discrepant samples are provided on a downloadable list at *<u>insellab.github.io</u>*.

DE results presented in this text are from comparisons between TCGA and GTEX samples, but we also include in our MDS analysis and in all downloadable counts files, data for TCGA-matched “normal” samples taken from tissue adjacent to tumors of TCGA patients. In general, normal TCGA and GTEX samples cluster closer together than do TCGA tumors and GTEX normal samples (**Figures S1G-J**).

The overlap, however, is not exact. In several cases, we found differences between TCGA normal and GTEX samples. It is unclear if these differences result from biological or technical factors; prior data show that tumors impact surrounding “normal” tissue and can also induce global changes [65–68]. Hence, we have not used batch-correction methods to account for these variations between TCGA normal and GTEX tissues. In general, DE of GPCRs is similar whether TCGA normal tissue or GTEX tissue is compared to TCGA tumor samples (e.g., **Figures S11E-F**) suggesting that such differences are unlikely to impact upon the general conclusions of this study.

For TCGA data, some recent efforts (e.g., http://bioinformatics.mdanderson.org/tcgambatch/) have been made to account for batch effects, though peer-reviewed studies have not yet established best practices for batch correction of TCGA data. Many TCGA datasets for individual tumor types contain numerous batches (with batches defined in terms of factors such as sequencing runs, or location of tissue collection), with small numbers of replicates in each batch. Given this, it is unclear if such batch corrections account for technical variation among batches or merely suppress biological variation, especially with the known heterogeneity among tumor samples. In light of this, we present all data from TCGA and GTEX without correcting for batch effects.

In nearly all cases, DE of GPCRs we highlight have large fold changes with high statistical significance (i.e., FDR << 0.05), such that minor technical variations ought not substantially impact our key findings. Moreover, the fact that TCGA tumors and GTEX normal tissues form distinct, separated clusters (and hence show a high degree of DE) is unlikely to be due to technical factors. In several cases (e.g., KICH matched normal vs GTEX kidney samples; **Fig S1G**), TCGA matched normal and GTEX normal tissues are in fact highly similar, whereas in other cases they are not (e.g., PRAD, **Fig S1J**). This suggests that technical factors between the two studies do not consistently skew/bias the two data sets vs. each other.

Analysis to identify dependence of GPCR expression on patient characteristics, such as sex or tumor stage was done as follows: GPCR expression for tumor samples normalized in CPM, output from edgeR was used to evaluate whether grouping samples together based on a specified attribute (e.g., sex) resulted in a statistically significant relative risk for that attribute tested for results in elevated GPCR expression in one group compared to the other. We tested this using Fisher’s exact test in R, yielding calculations of risk ratio and p-values, with the terms in the contingency table for Fisher’s test being the number of samples in each group with GPCR expression either above or below the population median. P-values were then adjusted for multiple testing using the p_adjust function in R, utilizing the Benjamini–Hochberg method. These adjusted p-values thus indicate whether any GPCRs have expression that is significantly associated with a given patient attribute. Specific attributes tested for were: sex, tumor stage, grade and pathological T (depending on availability of metadata for each tumor type and the number of available replicates). This method was also used to evaluate the significance of associations between expression of GPCRs and presence of specific driver mutations (e.g. presence or absence of mutations to TP53 or KRAS) and association between GPCR mRNA expression and the thresholded GISTIC 2.0 CNV call.

Survival analysis was performed in R using gene expression data normalized in CPM (i.e., units appropriate for comparisons <u>between samples</u>) via the “survival” and “survminer” packages. For each tumor type analyzed, samples were divided into two groups based on median expression of the gene being tested, and differences in survival between the two groups were calculated. The modified Peto-Peto (mPP) method was used to estimate the statistical significance of differences in survival rates. We noted cases where the effects of GPCR expression on survival were related/coupled (hence, the presence of composite markers of survival noted in the text). Thus, these statistical tests were not independent; given a lack of understanding about the nature of such dependencies (which to our knowledge were hitherto unknown), we did not adjust p-values for multiple testing. We also note that a) due to our use of relevant units for normalizing the gene expression data prior to performing survival analysis (CPM instead of units such as FPKM or RPKM) and b) the subdivision of broad TCGA categories into appropriate sub-types of tumors with consistent histological classification (e.g. dividing ESCA into adenocarcinomas and squamous cell carcinomas and analyzing each separately) our analysis yields associations of genes with survival that may differ in some cases from other sources that also provide survival analysis of TCGA data. We selected a significance threshold of p < 0.05, which is relatively lenient for survival analysis, as our initial priority was to minimize false negatives; we anticipate these analyses will need subsequent validation efforts in specific tumor types, with larger numbers of patients.

#### 4. Data and Software Availability

All data generated in this project are hosted at https://insellab.github.io/. These data are all open access. Relevant software such as R and edgeR are also freely available, refer key resources table.

