## Supplemental figures and text for "GPCRs show widespread differential mRNA expression and frequent mutation and copy number variation in solid tumors"

**Supplement 1: Supplementary Figures 1-13, Supplementary Tables 1-5 and Supplementary Notes 1-3.**

**Supplementary Figures legends.**

**Figure S1: Comparisons between GTEx normal samples and different TCGA tumor subtypes**

**(A-F) DE of GPCRs differs in different cancer subtypes within the same cancer category.**

(**A**) The repertoire of overexpressed (OE) GPCRs in Her2-positive and Triple-negative BRCA IDC (Breast Adenocarcinoma, Infiltrating Ductal Carcinoma). (**B**) For commonly OE GPCRs, the correlation of magnitude of fold-changes in expression in each tumor subtype compared to normal breast tissue.

(**C**) The repertoire of OE GPCRs in Classical and Follicular THCA (Thyroid Cancer). (**D**) For commonly OE GPCRs, the correlation of magnitude of fold-changes in expression in each tumor subtype compared to normal thyroid tissue.

(**E**) The repertoire of OE GPCRs in ESCA (Esophageal cancer) Adenocarcinoma and Squamous Cell Carcinoma. (**F**) For commonly OE GPCRs, the correlation of magnitude of fold-changes in expression in each tumor type compared to normal esophageal mucosal tissue.

BRCA IDC, either Her2-positive or Triple-negative, overexpresses a number of GPCRs. Several of these GPCRs are commonly overexpressed (**A**) but others are overexpressed (OE) in one type but not the other. In general, fold-changes of commonly overexpressed GPCRs correlated among cancer subtypes, but often with some scatter (**B**). Similar results are found in other tumors, (e.g. **C-D**), showing the degree of overlap of overexpressed GPCRs in Classical or Follicular THCA. Further, in tumors that occur in the same tissue, but with different precursor cells (e.g., squamous cell carcinomas vs adenocarcinomas), the repertoire of differentially expressed GPCRs is distinct. Panels **E-F** illustrate this for ESCA. Thus, in general, tumor types and subtypes with distinct histological classification possess distinct repertoires and changes in expression of GPCRs.

**(G-J) Differences between TCGA-matched "normal", GTEx normal tissue and tumors (KICH, LSQC (NOS).** MDS (multi-dimensional scaling) plots indicate that in some cases (**G, H**), TCGA-matched normal and GTEx normal tissue are similar, whereas in others (**I, J**) LSQC (NOS) and PRAD TCGA matched normal and GTEx normal samples differ, albeit these differences are smaller than the differences between normal tissue (from either source) and tumors. Differences in tumor biology with different tumor types influencing surrounding tissue to different degrees may explain the apparent differences in the "normal" tissue in the TCGA samples.

**Figure S2: Phylogenetic tree of GPCRs based on fold-change on solid tumors, for heatmap in figure 1A**

**Figure S3: Differential Expression (DE) of genes between PDAC tumors and normal pancreatic tissue**

**(A)** MDS (Multi-dimensional scaling) plot of gene expression in normal pancreatic tissue and PDAC tumors.
**(B)** Volcano plot showing significantly differentially expressed genes (FDR < 0.05) in red, with FDR plotted against fold-change.
**(C)** Smear plot showing genes with significant fold-change (red), with Fold change plotted against magnitude of gene expression in CPM.
**(D-F)** Expression of MMP11, S100A6 and LGALS3 in all samples for PDAC and normal pancreas, with medians (dashed lines) also indicated.
**(G-I)** Expression of EDNRB, ADGRG1 and GPR143 in all samples for primary and distant SKCM and normal skin, with medians (dashed lines) also indicated.
**(J)** The fraction of PDAC tumors which express GPRC5A above the indicated thresholds, compared to median expression in normal tissue.

**Figure S4: Differential Expression (DE) of genes between PDAC tumors and normal pancreatic tissue**

**(A)** The number of patients whose survival was tracked in the TCGA PDAC cohort at each time point, along with the rates of dropout and mortality.

**(B)** Network construction via STRING of the genes whose expression correlates with that of CCR6, CCR7, CXCR3 and CXCR4.

**Figure S5: Pathways enriched among genes associated with expression of GPCRs in SKCM**

**(A)** The combined, weighted expression of GPR143, ADGRG1 and EDNRB in SKCM shows positive correlation with expression of a subset of nearly 2000 genes.
**(B)** Network construction via STRING of the top 500 most strongly correlated genes from (**A**) illustrating the presence of genes related to the melanosome and to insulin response as examples of cancer-associated pathways in SKCM.
**(C)** Analysis of the 500 most strongly correlated genes via Enrichr shows enrichment of pathways such as transferring signaling, insulin response etc. among these positively correlated genes.

**Figure S6: Additional results on GPCR expression and DE: metastatic vs. primary tumors, primary vs. recurrent tumors and normal melanocytes vs. melanoma cell**

**(A-F). GPCR expression in metastatic and recurrent ovarian (OV), thyroid and breast cancer is similar to that in primary tumors.** Most TCGA tumor types have few replicates of metastases or recurrent tumors. However, for those with available data (SKCM, **Fig. 7**; BRCA, THCA and OV), we tested if GPCR expression is similar in primary tumors and metastases and in recurrent tumors. Panels **A-F** show that recurrent ovarian cancers, metastatic THCA (classical) tumors and BRCA IDC breast cancer, respectively, have similar GPCR expression, with identities of expressed GPCRs and magnitude of expression similar to primary tumors. One exception in BRCA IDC was *NPY1R*, which is more highly expressed in metastases than in primary tumors. All BRCA IDC subtypes were combined (e.g., Her2+, Triple Negative) for these analyses as there were only 6 metastases, precluding comparison of metastases in the different BRCA IDC subtypes.

**(A)** Correlation of expression of the 100 highest expressed genes in primary Ovarian Cancer (OV) with that of recurrent tumors. (**B**) Identity and relative expression of the 30 highest expressed GPCRs in primary and recurrent ovarian tumors. (**C**) Correlation of expression of the 100 highest expressed genes in primary THCA (classical) compared to that of metastatic tumors. (**D**) Identity and relative expression of the 30 highest expressed GPCRs in the primary and metastatic thyroid tumors.

**(E)** Correlation of expression of the 100 highest expressed genes in primary and metastatic BRCA (IDC, all types combined), excluding NPY1R which is much higher expressed in metastatic than primary tumors. (**F**) Identity and relative expression of the 30 highest GPCRs (including *NPY1R*) in primary and metastatic BRCA IDC tumors.

**(G, H). GPCR expression in non-diseased Melanocytes differs from that in Melanoma cells.** (**G**) GPCR expression in low-passage melanoma cancer cells (data mined from Müller et al., [40]) indicates that most highly expressed GPCRs in TCGA tumors are also highly expressed in melanoma cells. By comparison, non-diseased melanocytes [64] typically show much lower expression of these GPCRs than in melanoma cells. (**H**) GPCR expression (in FPKM) in low-passage melanoma cells and melanocytes is very poorly correlated.

**Figure S7: The GPCR expression 'repertoire' of normal tissues and solid tumors**

**(A, B)** The number of GPCRs, orphan GPCRs and GPCRs that couple to each G protein class in normal tissues **(A)** and solid tumors (**B**) and that have $\geq 0.1$ TPM median expression.

**(C, D)** GPCRs that couple to different G proteins compared to the total GPCR expression repertoire in normal tissue (**C**) and solid tumors (**D**).

**(E, F)** GPCR expression (TPM) in normal tissue (**E**) and solid tumors (**F**), and the number of GPCRs detected at different thresholds of expression. GPCRs typically account for $< 0.1\%$ of the tissue and tumor transcriptomes.

**Figure S8: GPCR mutation events and their relationship to genome-wide mutations**

**(A, B) Missense Mutations are the most frequent type of non-silent mutation of GPCRs in TCGA tumors, with *GPR98/ADGRV1* the most frequently mutated GPCR.** (**A**) The number of each type of somatic mutation for

GPCRs in TCGA tumors surveyed (n=5103 tumors with 32,727 somatic mutational events in GPCRs. **(B)** The same analysis for SKCM, which has the highest number (11,348) of GPCR mutational events, i.e., more than one third of all somatic mutation events but only ~9% of TCGA samples. **Table S3** lists the total number of mutation events for the most commonly mutated GPCRs; a complete list is provided in **Supplement 3**. GPCRs with the most frequent mutation events are also mutated in the largest number of tumors (**Fig 10A**). In the 5103 TCGA tumors surveyed, missense mutations are the most frequent type of non-silent GPCR mutation, occurring ~10-fold more frequently than frameshift deletions, the next most common type of non-silent mutation. Missense mutations were the most common type of mutational event for GPCRs in all cancer types except LIHC, which had a high frequency of frameshift deletions (n = 1875 events), with *GPR98* the most frequently so mutated (n = 79 events).

**(C) The number of somatic mutation events in all TCGA tumors for all annotated GPCRs. Inset**: the number of mutation events for the 10 most frequently mutated GPCRs. These data mirror those in **Figure 10** that show the number of tumors in TCGA that possess somatic, non-silent mutations to GPCRs.

**(D) The most frequently mutated GPCR and non-GPCR genes in solid tumors**: the number of tumors across all tumor types surveyed with somatic non-silent mutations for the genes indicated.

**(E-G) The number of mutated GPCRs in a tumor scales linearly with Nmut, the number of mutated genes per tumor genome.** For SKCM, LUAD and LSQC, their number of mutated GPCRs increases linearly with Nmut. This linear relationship is found with other tumor types and is nearly identical among tumor types, implying a general pattern of accumulation of genome-wide mutations and mutations in the GPCR super-family.

#### Figure S9: Additional results on the nature of non-silent GPCR mutations compared to certain other non-GPCR mutations

**(A, B) Location of silent and non-silent mutations of *GPR98*/*ADGRV1* in TCGA SKCM tumors (A) and *KRAS* mutations in PAAD tumors (B) accessed via Xena (xena.ucsc.edu).** Data are shown for 204/472 SKCM samples in which *GPR98* has silent or non-silent mutations. Introns are not included; hence the figure shows exonic locations of mutations. 132/186 PAAD samples had somatic *KRAS* mutations. Vertical gray bars indicate exons. Mutations of *GPR98*/*ADGRV1* are distributed along the length of the gene and are not enriched at specific locations or exons. Thus, large portions of *GPR98* represent mutational hotspots, findings that contrast with what occurs in driver mutations such as *KRAS*, in which mutations at specific sites and specific exons result in gain-of-function or loss-of-function respectively. **Figure S9B** shows that virtually all somatic (almost exclusively missense) mutations occur at exon 2 (resulting in an oncogenic *KRAS*) in PAAD, in contrast with the range of mutations in SKCM for *GPR98*, *GPR112* and other GPCR genes. The distribution of mutations along gene length is a feature of GPCR mutations in other cancers as well (e.g., BLCA, LUAD).

**(C, D) For primary (C) and distant metastatic (D) SKCM samples, relatively few genes show DE if one compares samples with or without LPHN2/ADGRL2 mutations**. Fewer than 100 genes increase or decrease >2-fold (with FDR<0.05 and >1 TPM median expression). Lists of DE genes for each case are provided in **Supplement 3***.* Red dots correspond to DE genes with FDR < 0.05. Similar data occur for other frequently mutated GPCRs (e.g., *GPR98*) in SKCM and in LUAD and BLCA.

**(E, G)** Correlation of GPCR expression (n = 100 highest expressed GPCRs) in primary and distant metastatic SKCM tumors respectively, with *GPR98* somatic non-silent mutations compared to tumors without *GPR98* mutation.

**(F, H)** The identities and median expression (TPM) of the 30 highest expressed GPCRs in primary and distant metastatic SKCM tumors respectively, for tumors with or without *GPR98* mutations.

#### Figure S10: GPR98 is part of a Tumor mutation profile

**(A-D) Mutations in GPR98 are frequently accompanied by mutations in other genes.** The proportion of tumors possessing somatic, non-silent mutations in GPR98 correlates with that of several other frequently mutated genes. Tumor types that show a high frequency of mutations to genes such as *TTN* and *MUC16* also typically show a high frequency of *GPR98* mutations and vice-versa for tumors with infrequent mutations to these genes. *TTN* and *MUC16* form a group of genes that, along with *GPR98*, are frequently mutated across a range of tumors. (**A**) The proportion of tumors in each TCGA tumor type that show mutations in *TTN*, *MUC16* and *GPR98*. (**B**) The correlation between the proportions of tumor samples possessing *TTN* mutations and *GPR98* mutations for the 20 TCGA tumor types shown above. (**C**) The same data for mutations in *MUC16* vs *GPR98* mutations.

**(E, F) Tumors with *GPR98* mutations frequently have mutations in other frequently-mutated genes.** SKCM tumors (n = 472) have a high frequency of *TTN*, *MUC16* and *DNAH5* somatic, non-silent mutations. Most SKCM tumors with somatic, non-silent mutations to *GPR98* also have mutations in these other genes.

**Figure S11: Concordance between protein and mRNA expression for GPRC5A (refer Supplemental note 3)**

**(A) *GPRC5A* mRNA expression in normal tissue.**
Median normalized mRNA expression from GTEx, in CPM of GPRC5A in normal tissues profiled by RNAseq
**(B) *GPRC5A* protein expression in normal tissue.**
Protein abundance (by immunohistochemistry from the human protein atlas; Uhlen et al., {2017}) of GPRC5A in normal tissue
**(C) *GPRC5A* mRNA expression in tumors.**
Median normalized mRNA expression (in CPM in TCGA) of GPRC5A in tumors profiled by RNAseq. For TCGA tumor types with multiple subtypes (e.g., LUAD), values are the median for all subtypes of the tumor type to facilitate comparison with protein atlas data, in which TCGA tumor types are not separated into subtypes.
**(D) *GPRC5A* protein expression in tumor tissue.**
Protein abundance (by immunohistochemistry from the human protein atlas; proteinatlas.org) of GPRC5A in a range of tumor types. Data are provided for multiple replicates at https:/www.proteinatlas.org/ENSG00000013588-GPRC5A/pathology; the proportion of samples staining at different intensity levels are shown.

**Figure S12: Supplemental results validating DE analysis methods**

**(A, B) EBseq and EdgeR yield similar results for DE analysis.** (**A**) Comparison of fold-changes determined via EdgeR for the 10,000 genes with lowest FDRs (i.e., the most significantly altered genes), compared to the fold-changes for the same genes determined via Ebseq, using data for PDAC tumors compared to normal pancreatic tissue. (**B**) The same comparison (between EBseq and EdgeR) for the 100 GPCRs with lowest FDRs calculated via EdgeR. Dashed lines indicate linear fits. The two methods make different statistical assumptions but yield similar results, especially for GPCRs. We chose to use EdgeR rather than Ebseq based on the much lower processing times in EdgeR for the large files we generated. These results, along with the similarity of DE analysis between upper-quartile and TMM normalized data helps allay concerns regarding the validity of this analysis stemming from the large numbers of DE genes found when comparing tumor and normal samples.

**(C, D) TMM and Upper-quartile normalization in EdgeR yield similar results for DE analysis. (C)** Comparison of fold-changes determined via EdgeR for the 10,000 genes with lowest FDRs (i.e., the most significantly altered genes), compared to fold-changes for the same genes determined via Ebseq, using data for PDAC tumors compared to normal pancreatic tissue. (**D**) The same comparison for the 100 GPCRs with lowest FDRs calculated via EdgeR. Dashed lines indicate linear fits.

**(E, F)** DE analysis of fold-changes for GPCRs identified as meaningfully increased or decreased expression in KICH (**E**) and LSQC (NOS) (**F**) is similar (especially for highly expressed GPCRs with high-fold-change) whether one compares these tumors with GTEx normal tissue or TCGA-matched normal samples.

**Figure S13: Differential Expression of GPCRs with and without corrections for batch effects**

**(A, B)** Correlation of median (**a**) and average (**b**) expression of GPCRs in TCGA PDAC samples, with and without batch corrections performed for TCGA plate ID.
**(C, D)** Correlation of median (**c**) and average (**d**) expression of GPCRs in OV (TCGA Ovarian Cancer) samples, with and without batch corrections performed for TCGA plate ID.

#### Supplementary Tables

**Table S1**. **The types of cancers and number of replicates from TCGA surveyed for GPCR mutations and CNV**. Mutation and CNV data were obtained from *xena.ucsc.edu*. For mutations, data were generated at the Broad institute sequencing center, except for * Baylor College of Medicine Sequencing Center and # Washington University Sequencing Center. Mutation data were generated via automated pipelines from the respective sources, hosted at *xena.ucsc.edu*.

| Cancer Type | Replicates for Somatic Mutation Analysis | Replicates for Copy Number Variation analysis |
|---|---|---|
| Adrenocortical Cancer (ACC) * | 91 | 90 |
| Bladder Cancer (BLCA) | 396 | 408 |
| Breast Cancer (BRCA) # | 771 | 1080 |
| Cervical Squamous Cell Carcinoma (CESC) | 40 | 295 |
| Colon Cancer (COAD) * | 217 | 451 |
| Esophageal Cancer (ESCA) | 184 | 184 |
| Kidney Chromophobe (KICH) | 66 | 66 |
| Kidney Clear cell carcinoma (KIRC) | 213 | 528 |
| Kidney Papillary cell carcinoma (KIRP) | 113 | 288 |
| Liver hepatocellular carcinoma (LIHC) | 202 | 370 |
| Lung adenocarcinoma (LUAD) | 543 | 516 |
| Lung squamus cell carcinoma (LSQC) | 178 | 501 |
| Ovarian carcinoma (OV) | 142 | 367 |
| Pancreatic adenocarcinoma (PAAD) | 185 | 579 |
| Prostate adenocarcinoma (PRAD) | 499 | 184 |
| Skin cutaneous melanoma (SKCM) | 472 | 492 |
| Stomach adenocarcinoma (STAD) | 91 | 441 |
| Testicular carcinoma (TGCT) | 139 | 150 |
| Thyroid carcinoma (THCA) | 504 | 499 |
| Uterine carcinomasarcoma (UCS) | 57 | 56 |
| Total | 5103 | 7545 |

**Table S2**. **The normal tissue types and number of replicates (GTEx database) used for differential expression (DE) analysis of RNA-seq data in normal tissue compared to tumors (TCGA)**. Colonic adenocarcinoma (COAD) tumors in the sigmoid and transverse colon were compared with normal tissue from those regions. For esophageal tumors (adenocarcinomas and squamous cell carcinomas), DE analysis was compared to esophageal mucosal tissue; data in GTEx for esophageal muscularis tissue were not used in this analysis. Data for sun-exposed skin were compared to melanomas; similar DE results were found if non-sun exposed skin (in GTEx) was used. Only 9 of 952 TCGA BRCA samples for which gender was recorded were from males, hence we only used GTEx breast tissue from females as reference normal tissue for DE analysis of BRCA.

| Normal Tissue Type | Number of replicates |
|---|---|
| Adrenal Gland | 125 |
| Bladder | 9 |
| Breast (Female) | 79 |
| Cervix | 10 |
| Colon (Sigmoid) | 140 |
| Colon (Transverse) | 163 |
| Esophagus Mucosa | 269 |
| Kidney | 27 |
| Liver | 110 |
| Lung | 287 |
| Skin (sun exposed) | 323 |
| Ovary | 88 |
| Pancreas | 165 |
| Prostate | 100 |
| Stomach | 172 |
| Testicle | 165 |
| Thyroid | 278 |
| Uterus | 78 |

**Table S3. The total number of different types of somatic mutational events for the 30 most frequently mutated GPCRs in the TCGA tumors surveyed**. A complete list is provided as downloadable supplemental material at *insellab.github.io*. In addition, a breakdown of numbers of mutation events for each GPCR in each tumor type is provided in **Supplement 2**.

| Gene | Frame_Shift_Del | Frame_Shift_Ins | In_Frame_Del | In_Frame_Ins | Missense_Mutation | Nonsense_Mutation | Nonstop_Mutation | RNA | Silent | Splice_Site | Translation_Start_Site | Total |
|---|---|---|---|---|---|---|---|---|---|---|---|---|
| GPR98 | 96 | 7 | 1 | 0 | 607 | 53 | 0 | 0 | 213 | 23 | 0 | 1000 |
| GPR112 | 28 | 8 | 0 | 0 | 349 | 27 | 0 | 0 | 150 | 6 | 0 | 568 |
| BAI3 | 21 | 1 | 0 | 0 | 292 | 21 | 0 | 0 | 86 | 19 | 1 | 441 |
| GRM8 | 22 | 7 | 0 | 0 | 198 | 15 | 0 | 1 | 95 | 3 | 0 | 341 |
| CASR | 19 | 0 | 1 | 0 | 185 | 6 | 0 | 0 | 127 | 0 | 0 | 338 |
| LPHN2 | 17 | 0 | 1 | 0 | 209 | 17 | 0 | 0 | 79 | 8 | 0 | 331 |
| LPHN3 | 19 | 2 | 0 | 0 | 207 | 12 | 0 | 0 | 79 | 11 | 1 | 331 |
| GPR158 | 12 | 2 | 0 | 0 | 203 | 14 | 0 | 0 | 90 | 8 | 0 | 329 |
| GRM7 | 18 | 2 | 0 | 0 | 190 | 16 | 0 | 0 | 94 | 6 | 0 | 326 |
| CELSR3 | 38 | 3 | 0 | 0 | 159 | 18 | 0 | 0 | 96 | 3 | 0 | 317 |
| GRM3 | 5 | 1 | 0 | 0 | 189 | 14 | 0 | 0 | 94 | 5 | 2 | 310 |
| CELSR2 | 47 | 0 | 1 | 21 | 156 | 7 | 0 | 0 | 70 | 3 | 0 | 305 |
| CELSR1 | 27 | 3 | 0 | 0 | 140 | 10 | 0 | 0 | 77 | 12 | 0 | 269 |
| GPR179 | 36 | 3 | 1 | 0 | 151 | 11 | 0 | 0 | 51 | 3 | 0 | 256 |
| GRM1 | 15 | 2 | 1 | 0 | 162 | 8 | 0 | 0 | 53 | 2 | 1 | 244 |
| GRM5 | 13 | 2 | 1 | 0 | 147 | 10 | 0 | 0 | 59 | 0 | 1 | 233 |
| ELTD1 | 11 | 1 | 0 | 0 | 132 | 16 | 0 | 0 | 43 | 13 | 0 | 216 |
| GRM6 | 6 | 6 | 0 | 0 | 115 | 6 | 0 | 0 | 77 | 2 | 0 | 212 |
| CHRM2 | 13 | 3 | 0 | 0 | 131 | 6 | 0 | 2 | 55 | 0 | 0 | 210 |
| GPR116 | 16 | 1 | 0 | 0 | 113 | 4 | 0 | 0 | 72 | 4 | 0 | 210 |
| EMR1 | 8 | 0 | 0 | 0 | 113 | 5 | 0 | 0 | 72 | 4 | 0 | 202 |
| FSHR | 16 | 1 | 0 | 0 | 111 | 11 | 0 | 0 | 58 | 4 | 1 | 202 |
| BAI1 | 22 | 2 | 3 | 0 | 102 | 5 | 0 | 0 | 59 | 7 | 0 | 200 |
| GPRC6A | 6 | 4 | 0 | 2 | 117 | 9 | 0 | 0 | 58 | 0 | 0 | 196 |
| DRD5 | 3 | 0 | 0 | 0 | 107 | 2 | 0 | 0 | 74 | 0 | 0 | 186 |
| TAS1R2 | 8 | 0 | 0 | 0 | 104 | 6 | 0 | 0 | 66 | 1 | 0 | 185 |
| CHRM3 | 10 | 0 | 0 | 0 | 110 | 9 | 1 | 0 | 53 | 0 | 0 | 183 |
| GPR133 | 17 | 0 | 1 | 0 | 85 | 6 | 0 | 0 | 50 | 11 | 1 | 171 |
| HCRTR2 | 9 | 10 | 0 | 0 | 93 | 8 | 0 | 0 | 40 | 9 | 0 | 169 |
| BAI2 | 26 | 1 | 2 | 0 | 85 | 3 | 0 | 0 | 50 | 2 | 0 | 169 |

**Table S4. GPCRs showing DE in multiple tumor types.** (**Left**) **Multiple GPCRs have increased expression in multiple types of cancers**. The 25 most widely/commonly overexpressed (OE) GPCRs are listed along with the number of cancer types in which they are OE. (**Center) The same tabulation for the 25 most commonly OE GPCRs that are targets for approved drugs**. (**Right**) **The 25 GPCRs most frequently reduced in expression in tumors compared to normal tissue**. GPCRs were considered to have DE if fold-changes were >2, FDR <0.05 and median expression in tumors was >1 TPM. Details of the tumor types where such DE occurs, the size of fold-changes etc. are available in **Supplement 3** and at *insellab.github.io*.

A list of GPCRs that are targets for approved drugs was obtained by querying the IUPHAR and CHEMBL databases for lists of approved drugs and the genes that they target. Most such GPCRs are targets for drugs approved by the FDA. The resulting list of 'druggable' GPCRs (and the approving agency) is provided in **Supplement 3**.

**Overexpressed GPCRs**

| Gene Symbol | Number of cancer types |
|---|---|
| FPR3 | 38 |
| F2RL1 | 37 |
| GPR160 | 35 |
| GPR143 | 33 |
| P2RY6 | 33 |
| APLNR | 31 |
| OPN3 | 31 |
| CXCR3 | 30 |
| CCR1 | 29 |
| FZD2 | 28 |
| LPAR5 | 28 |
| CELSR3 | 28 |
| ADORA2B | 27 |
| CCR5 | 27 |
| PTAFR | 27 |
| GPR39 | 27 |
| F2R | 26 |
| C3AR1 | 25 |
| GPRC5A | 25 |
| CELSR1 | 24 |
| CXCR4 | 22 |
| CCRL2 | 21 |
| ADGRG1 | 20 |
| ADGRG6 | 20 |
| CXCR6 | 20 |

**Overexpressed GPCR drug targets**

| Gene Symbol | Number of cancer types |
|---|---|
| GPR143 | 33 |
| P2RY6 | 33 |
| ADORA2B | 27 |
| CCR5 | 27 |
| F2R | 26 |
| CXCR4 | 22 |
| ADORA1 | 17 |
| P2RY13 | 17 |
| P2RY2 | 17 |
| HRH1 | 14 |
| GPR35 | 14 |
| HTR1D | 11 |
| CYSLTR1 | 10 |
| GPR68 | 10 |
| HCAR1 | 10 |
| ADORA3 | 9 |
| ADRA2A | 9 |
| S1PR5 | 9 |
| SMO | 9 |
| ADORA2A | 8 |
| BDKRB2 | 8 |
| ACKR3 | 8 |
| EDNRA | 7 |
| S1PR3 | 7 |
| CCR4 | 7 |

**Down-regulated GPCRs**

| Gene Symbol | Number of cancer types |
|---|---|
| GABBR1 | 40 |
| GPR146 | 35 |
| ACKR1 | 34 |
| MRGPRF | 30 |
| LTB4R | 28 |
| S1PR1 | 24 |
| ADGRA2 | 23 |
| PTGIR | 22 |
| FZD4 | 22 |
| ADGRL4 | 21 |
| LPAR1 | 21 |
| EDNRB | 20 |
| GPR4 | 20 |
| MC1R | 19 |
| ADGRD1 | 18 |
| ADGRF5 | 17 |
| VIPR1 | 16 |
| ACKR3 | 15 |
| LPAR6 | 15 |
| ADORA2A | 13 |
| PTH1R | 13 |
| F2RL3 | 13 |
| ADRA2C | 12 |
| CALCRL | 12 |
| GPR162 | 12 |

**Table S5. GPCRs that show frequent overexpression typically show infrequent mutation and vice versa.** (**Left**) The 30 most frequently overexpressed GPCRs across the 45 different types/subtypes of cancer profiled, along with the number of TCGA tumors (of 5103 total) in which they have somatic, non-silent mutations. (**Right**) The same data, sorted for the 30 GPCRs mutated in the highest number of tumors. Among the most frequently mutated GPCRs, the *CELSR* genes are also frequently overexpressed, but most other frequently mutated genes are not.

| Gene Symbol as per IUPHAR | Hg19 Synonym, used in analysis of mutations | Overexpressed in how many cancers types? | Number of Tumors with a mutation |
|---|---|---|---|
| FPR3 | FPR3 | 38 | 42.5 |
| F2RL1 | F2RL1 | 37 | 31.5 |
| GPR160 | GPR160 | 35 | 16 |
| GPR143 | GPR143 | 33 | 16 |
| P2RY6 | P2RY6 | 33 | 18 |
| APLNR | APLNR | 31 | 46.5 |
| OPN3 | OPN3 | 31 | 26 |
| CXCR3 | CXCR3 | 30 | 26 |
| CCR1 | CCR1 | 29 | 39 |
| CELSR3 | CELSR3 | 28 | 154.5 |
| FZD2 | FZD2 | 28 | 39 |
| LPAR5 | LPAR5 | 28 | 15.5 |
| ADORA2B | ADORA2B | 27 | 11 |
| CCR5 | CCR5 | 27 | 38 |
| GPR39 | GPR39 | 27 | 38 |
| PTAFR | PTAFR | 27 | 17 |
| F2R | F2R | 26 | 48 |
| C3AR1 | C3AR1 | 25 | 47.5 |
| GPRC5A | GPRC5A | 25 | 22 |
| CELSR1 | CELSR1 | 24 | 159 |
| CXCR4 | CXCR4 | 22 | 29 |
| CCRL2 | CCRL2 | 21 | 10 |
| ADGRG1 | GPR56 | 20 | 34 |
| ADGRG6 | GPR126 | 20 | 68.5 |
| CXCR6 | CXCR6 | 20 | 15 |
| ADGRB2 | BAI2 | 19 | 93.5 |
| GPR132 | GPR132 | 19 | 26 |
| GPRC5C | GPRC5C | 19 | 50 |
| CCR7 | CCR7 | 18 | 25 |
| CELSR2 | CELSR2 | 18 | 157.5 |

| Gene Symbol as per IUPHAR | Hg19 Synonym, used in analysis of mutations | Overexpressed in how many cancers types? | Number of Tumors with a mutation |
|---|---|---|---|
| ADGRV1 | GPR98 | 12 | 432 |
| ADGRG4 | GPR112 | 0 | 286 |
| ADGRB3 | BAI3 | 0 | 254.5 |
| ADGRL3 | LPHN3 | 3 | 205 |
| GPR158 | GPR158 | 4 | 190 |
| ADGRL2 | LPHN2 | 6 | 188 |
| GRM8 | GRM8 | 6 | 185.5 |
| GRM7 | GRM7 | 0 | 174 |
| GRM3 | GRM3 | 0 | 166.5 |
| CELSR1 | CELSR1 | 24 | 159 |
| CELSR2 | CELSR2 | 18 | 157.5 |
| GRM1 | GRM1 | 0 | 156.5 |
| CELSR3 | CELSR3 | 28 | 154.5 |
| CASR | CASR | 0 | 152.5 |
| ADGRL4 | ELTD1 | 5 | 143.5 |
| GRM5 | GRM5 | 0 | 143 |
| GPR179 | GPR179 | 0 | 140.5 |
| CHRM2 | CHRM2 | 0 | 122.5 |
| FSHR | FSHR | 0 | 115.5 |
| ADGRB1 | BAI1 | 0 | 114.5 |
| GRM6 | GRM6 | 0 | 110 |
| CHRM3 | CHRM3 | 6 | 108 |
| HCRTR2 | HCRTR2 | 0 | 106.5 |
| LHCGR | LHCGR | 0 | 106 |
| ADGRF5 | GPR116 | 2 | 103 |
| TAS1R2 | TAS1R2 | 0 | 99.5 |
| ADGRE1 | EMR1 | 1 | 96 |
| CALCR | CALCR | 0 | 95 |
| LGR5 | LGR5 | 13 | 94 |
| ADGRB2 | BAI2 | 19 | 93.5 |

#### Supplementary Notes

##### Supplementary Note 1: Concordance of mRNA and protein expression

Multiple recent studies, exemplified by citations below, support the notion that mRNA expression broadly predicts protein expression and that mRNA expression accounts for the majority of protein expression, compared to other factors such as translational regulation.

Using a yeast model, Csardi et al. [53] have suggested that mRNA expression is highly predictive of steady-state protein abundance. Applying noise-correction techniques to normalize mRNA and protein expression data, this study reported that mRNA accounts for ~85% of steady-state protein abundance. Similarly, [69] showed that in a frog egg model, if dynamics of protein decay and synthesis are accounted for, mRNA expression is highly predictive of protein expression. Data in *E. coli* [55] also suggest that mRNA expression is the primary determinant of protein expression, explaining a larger degree of protein expression than variations in translation.

Further, a study using a drug-treated ovarian cancer mouse xenograft model [56] showed that differentially expressed genes show concordance with changes in protein abundance. Accounting for gene-specific corrections, [54] showed that mRNA abundance is highly predictive of protein abundance in human tissue. Abundances of individual genes and their corresponding proteins correlated with one another by gene-specific proportionality coefficients that were largely independent of tissue and cell type. This result implies a linear, proportional relationship between mRNA and protein, such that changes in mRNA expression will (in general) be expected to predict similar changes in protein expression.

Factors such as those above likely account for much of the discrepancy between gene and protein expression noted in prior studies (e.g., [70]). Re-examination by [57] of the protein abundance data in the latter study (with corrected protein abundance estimates via alternate normalization strategies, using a range of housekeeping proteins) suggested that mRNA accounts for at least 56% of protein abundance and perhaps as high as 80%. The recent studies highlighted above lead us to conclude that data in the current study on differential gene expression likely predict changes in abundance of GPCRs at the protein level in tumors.

##### Supplementary Note 2: Correspondence of mRNA and protein expression of *GPRC5A*

The Human Protein Atlas (HPA) (www.proteinatlas.org) provides immunostaining data of normal tissue and tumors, quantifying protein abundance empirically as ‘high’, ‘medium’, ‘low’ and ‘not detected’. A principal challenge in quantifying GPCR protein abundance is the paucity of specific/selective, well-validated antibodies. The difficulties in GPCR protein detection underline the need for positive and negative controls (such as knockdowns/knockouts and overexpressing cells or tissues) for antibody validation. GPCRs are generally low-abundance proteins, making their detection challenging by proteomic methods.

Of GPCRs widely overexpressed in tumors (**Table S4**) that were also tested in HPA, we found that only GPRC5A currently had an antibody validated using controls indicated above (Atlas Antibodies, Cat#HPA007928). This antibody was also independently tested and validated in studies on pancreatic cancer cell lines [12], including knockdowns of *GPRC5A*. We thus compared protein abundance in normal and tumor tissue from HPA data that uses this well-validated antibody compared to our estimates for *GPRC5A* mRNA abundance.

Normal tissue shows low to moderate mRNA expression of *GPRC5A* except in lung tissue, in which this GPCR is highly expressed (**Fig S11A**), a result in agreement with antibody staining data from HPA (**Fig S11B**). Moreover, staining of multiple tumor replicates (**Fig S10D**) indicates a high frequency of positive staining in pancreatic, colon, stomach, cervical and lung tumors, consistent with *GPRC5A* mRNA expression (**Fig S11C**). Broadly stated, transcriptomic analysis suggested that GPRC5A is generally low-expressed in normal tissue besides the lung, but is widely expressed in a range of tumors. This same general pattern is reflected by *GPRC5A* protein abundance from HPA.

### SUPPLEMENTARY FIGURE 1

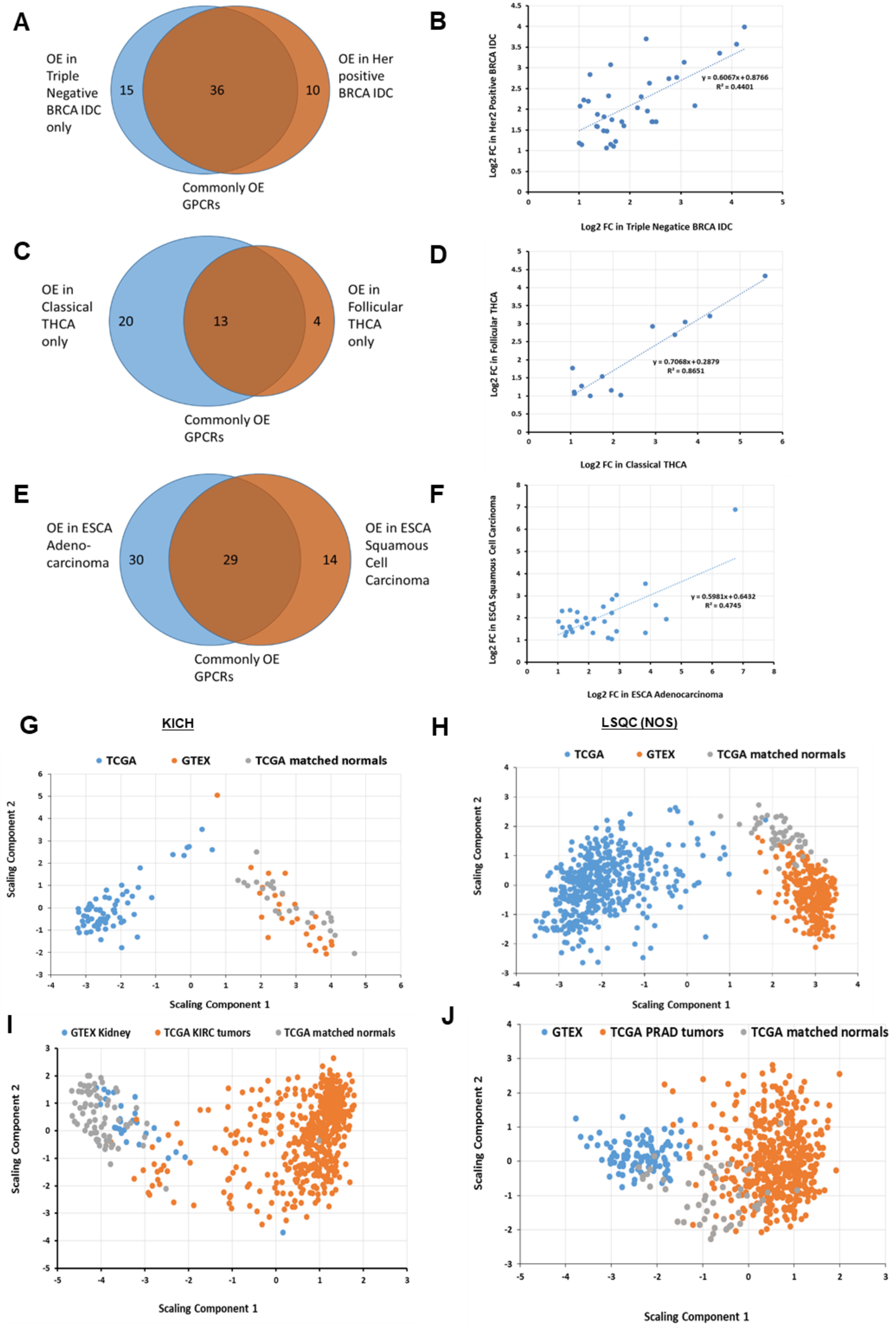

### SUPPLEMENTARY FIGURE 2

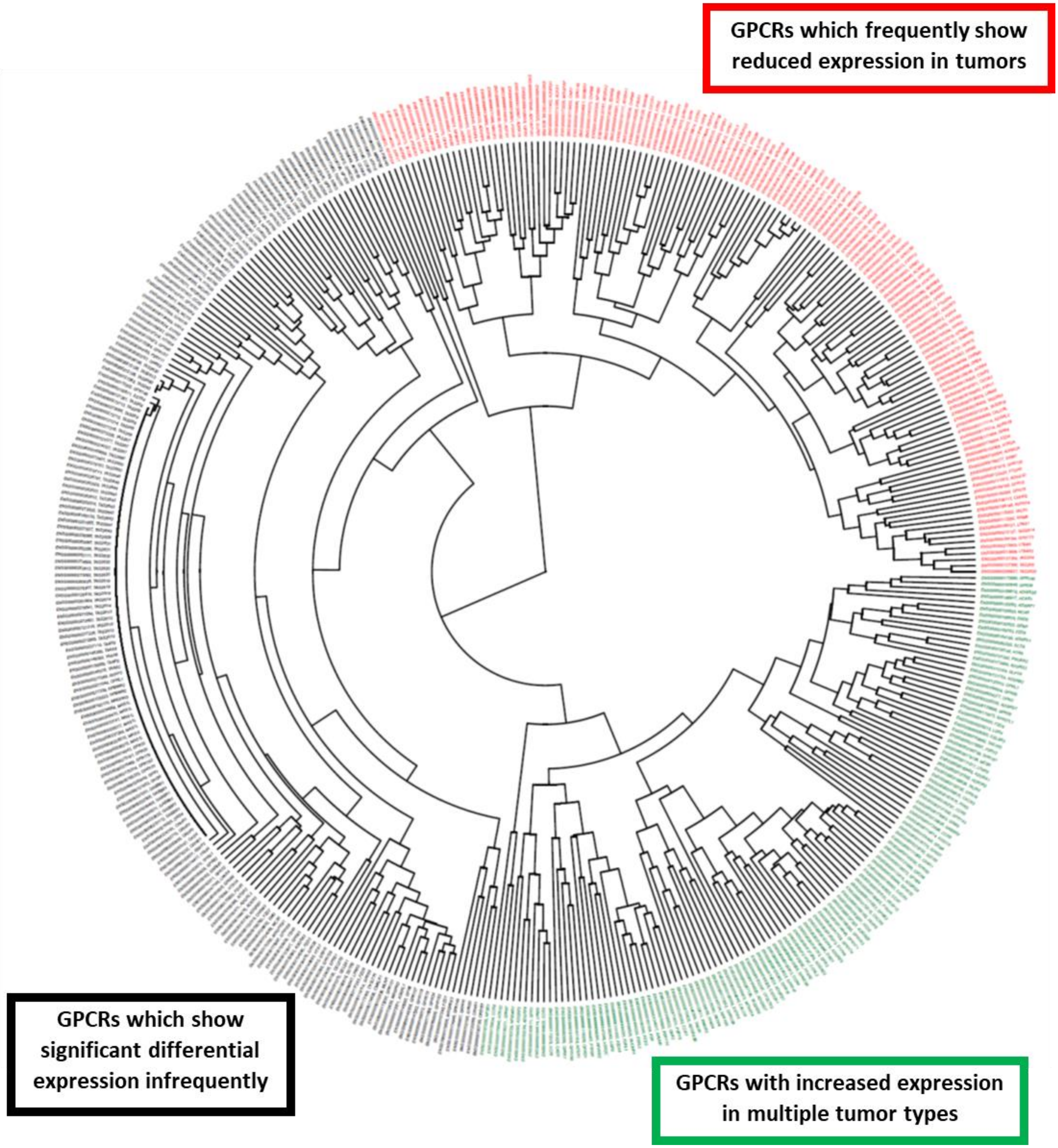

### SUPPLEMENTARY FIGURE 3

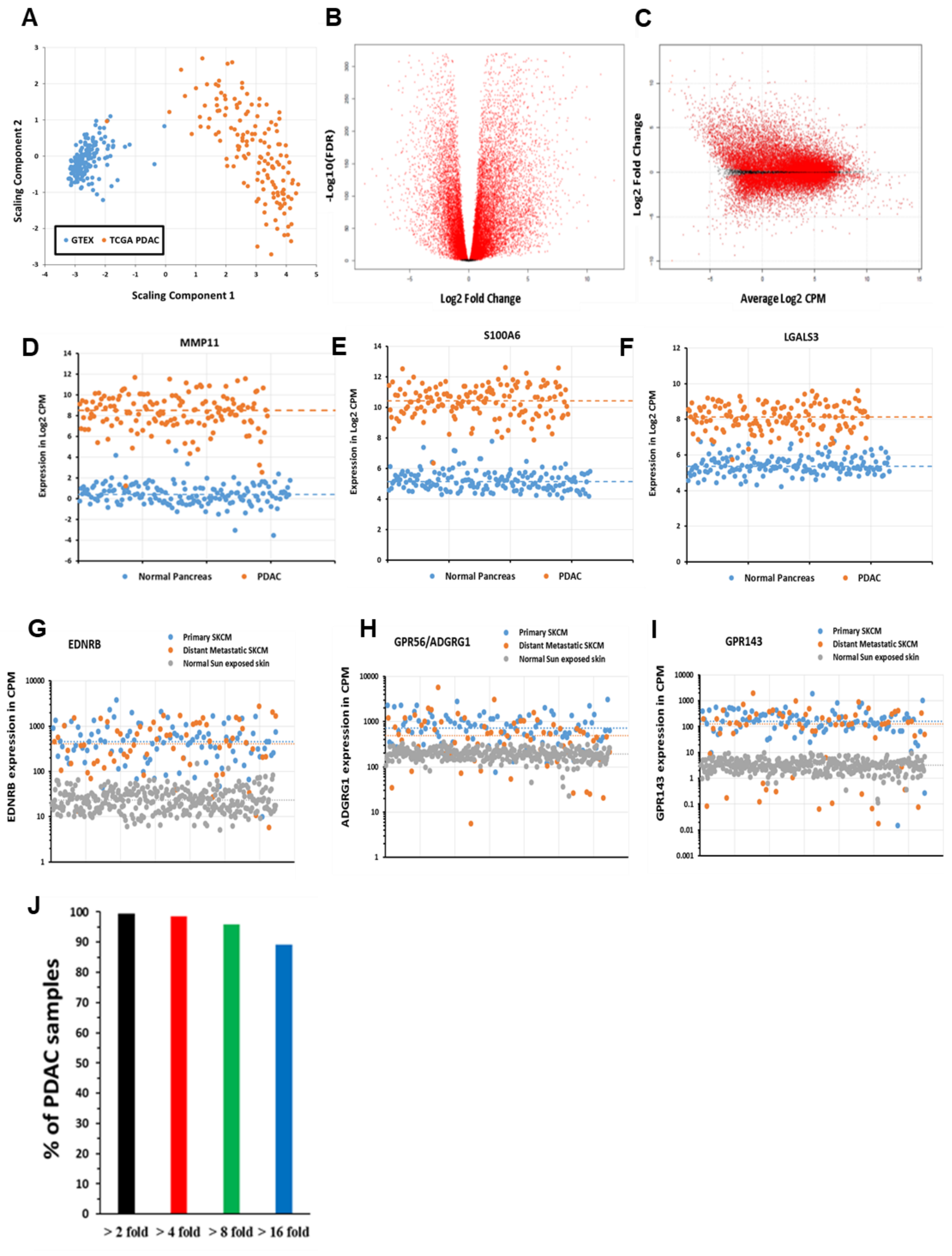

### SUPPLEMENTARY FIGURE 4

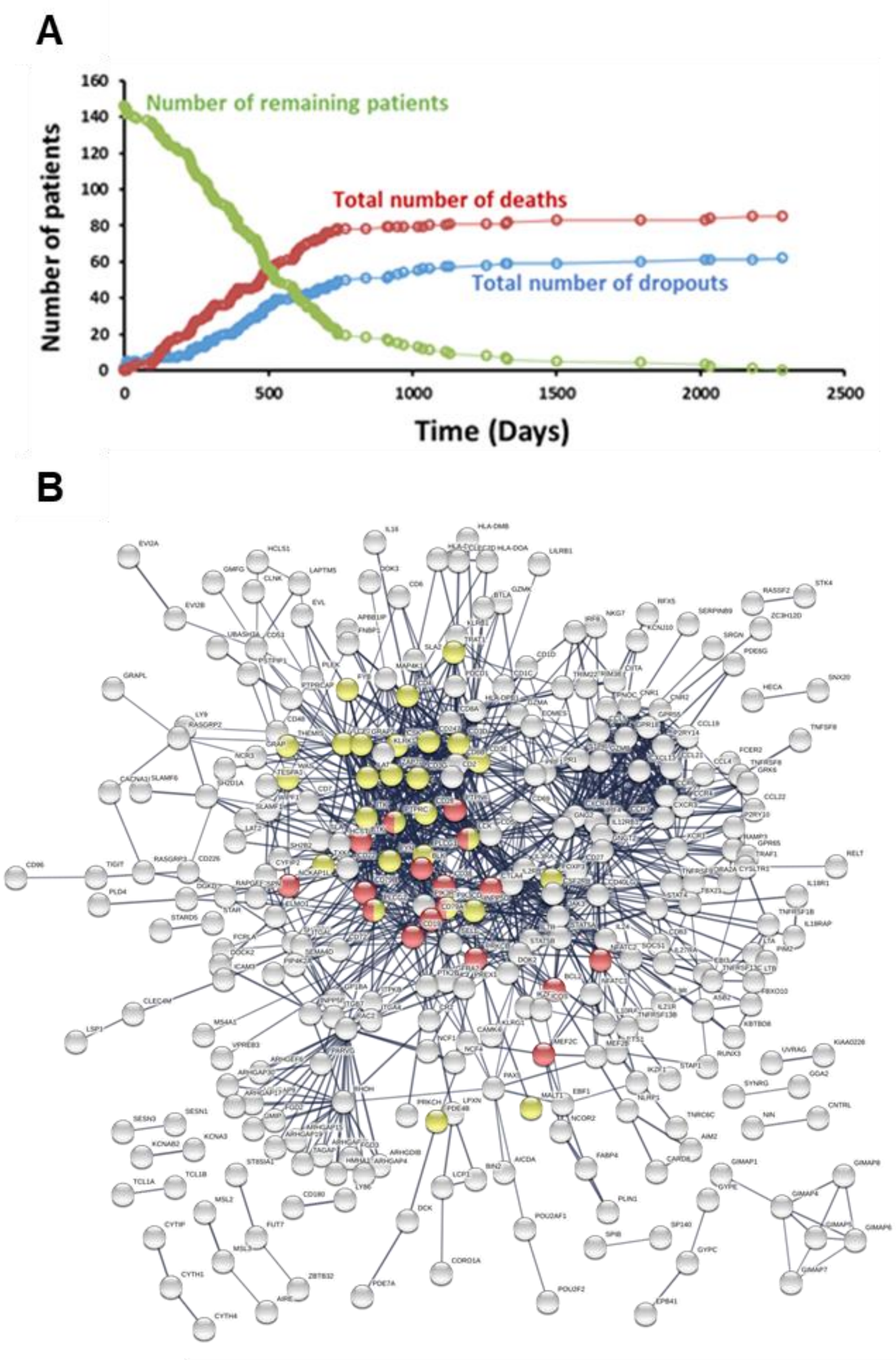


**Yellow: T cell receptor signaling pathway**

**Red: B cell receptor signaling pathway**

### SUPPLEMENTARY FIGURE 5

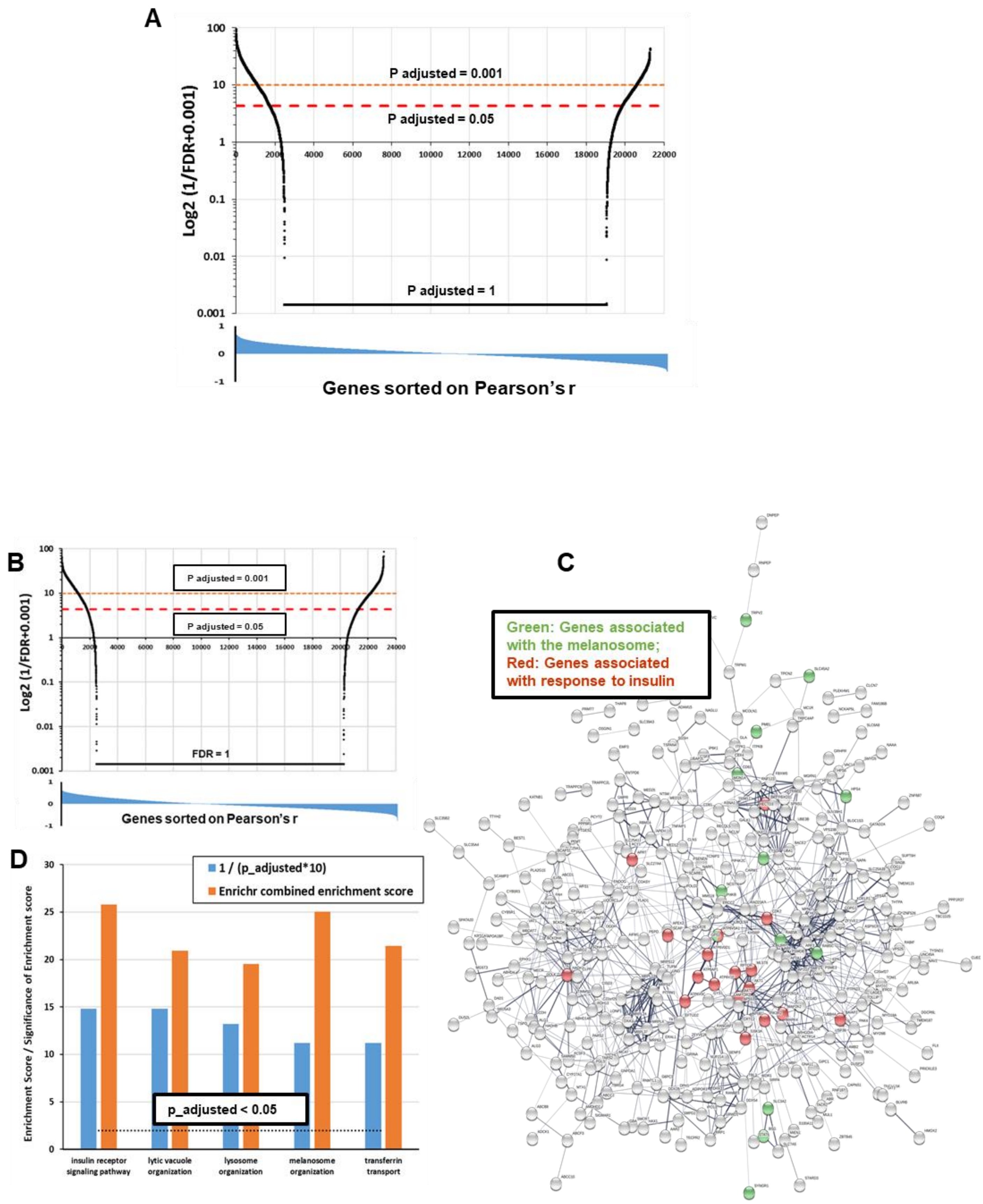

### SUPPLEMENTARY FIGURE 6

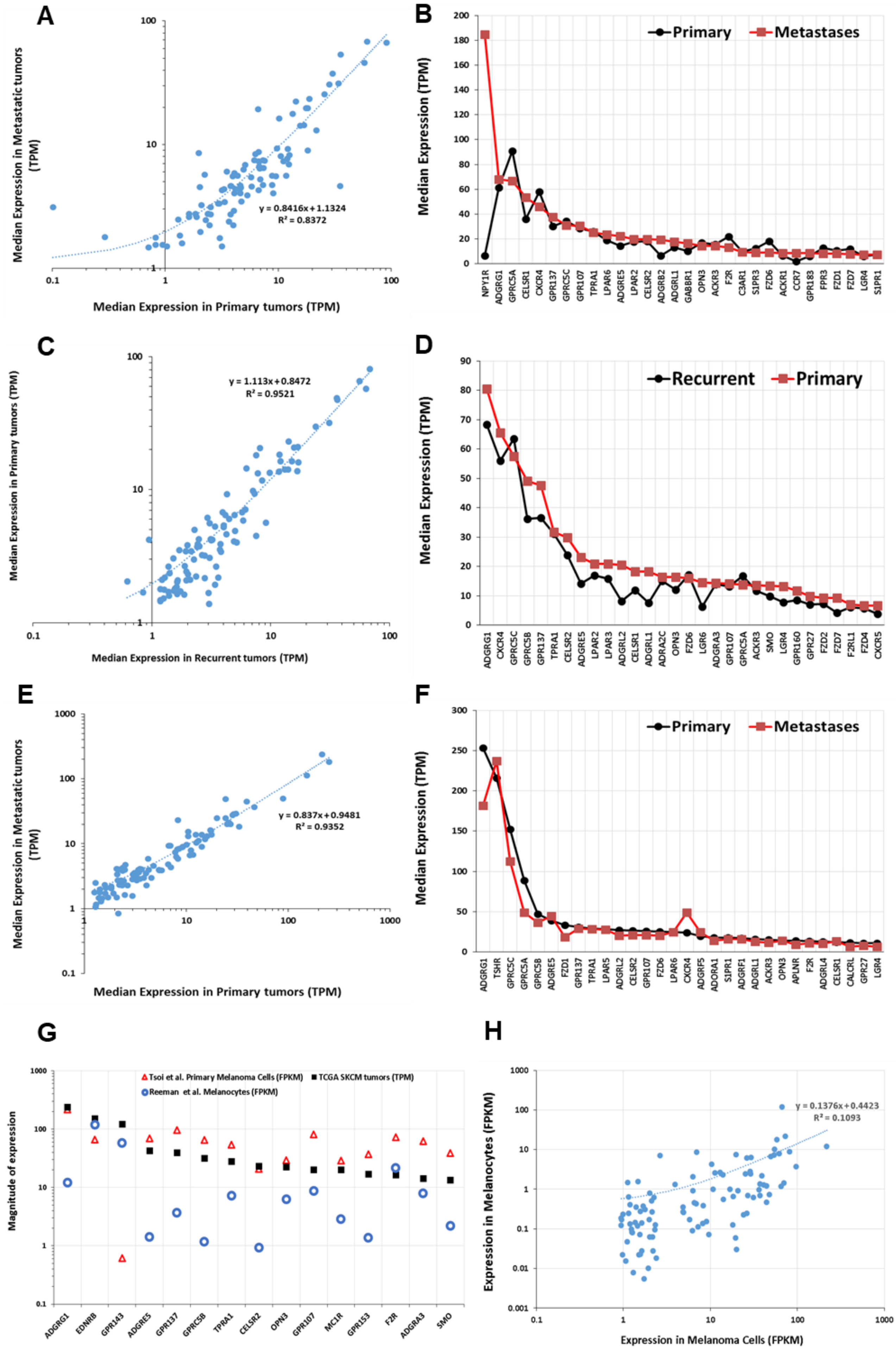

### SUPPLEMENTARY FIGURE 7

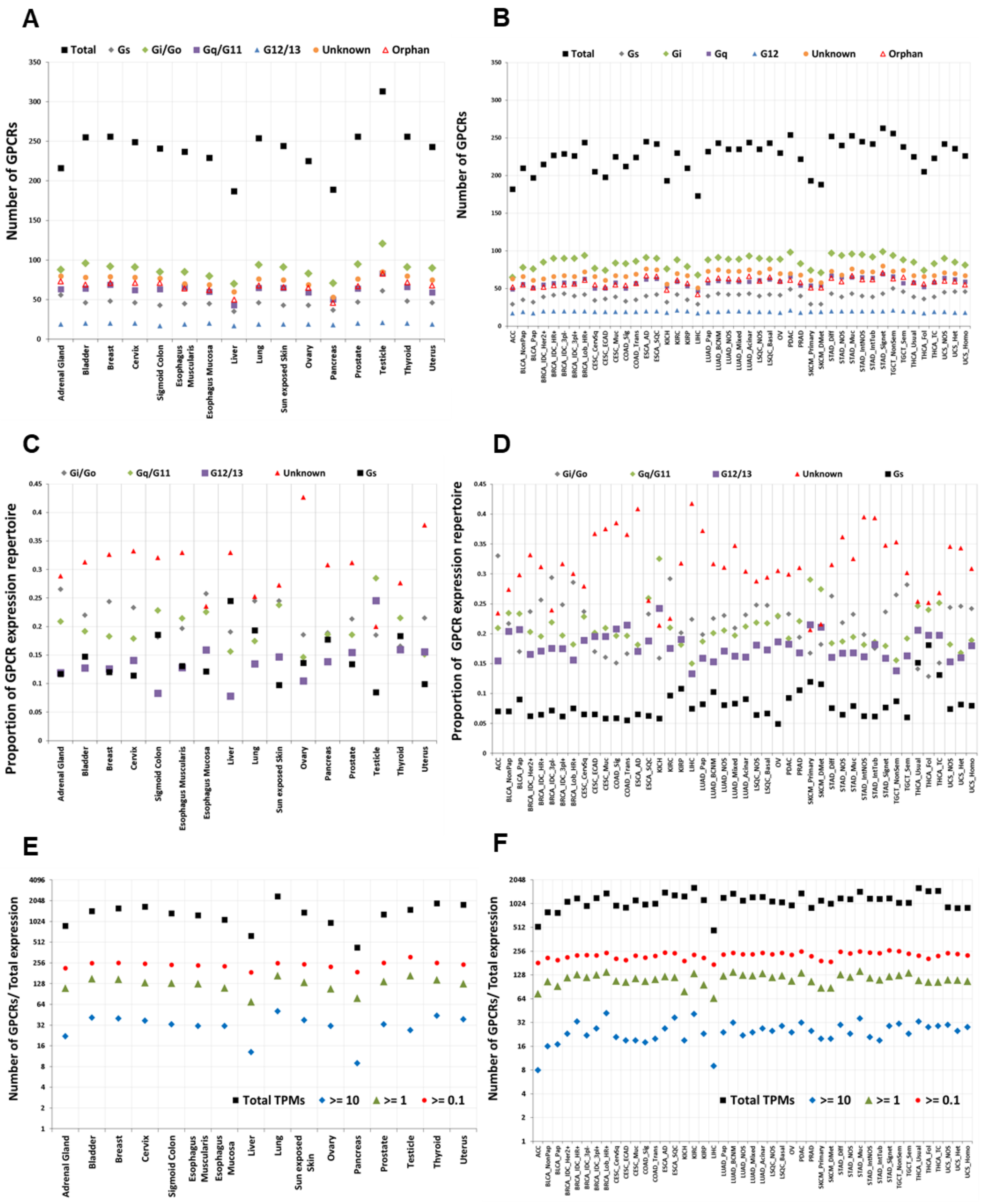

### SUPPLEMENTARY FIGURE 8

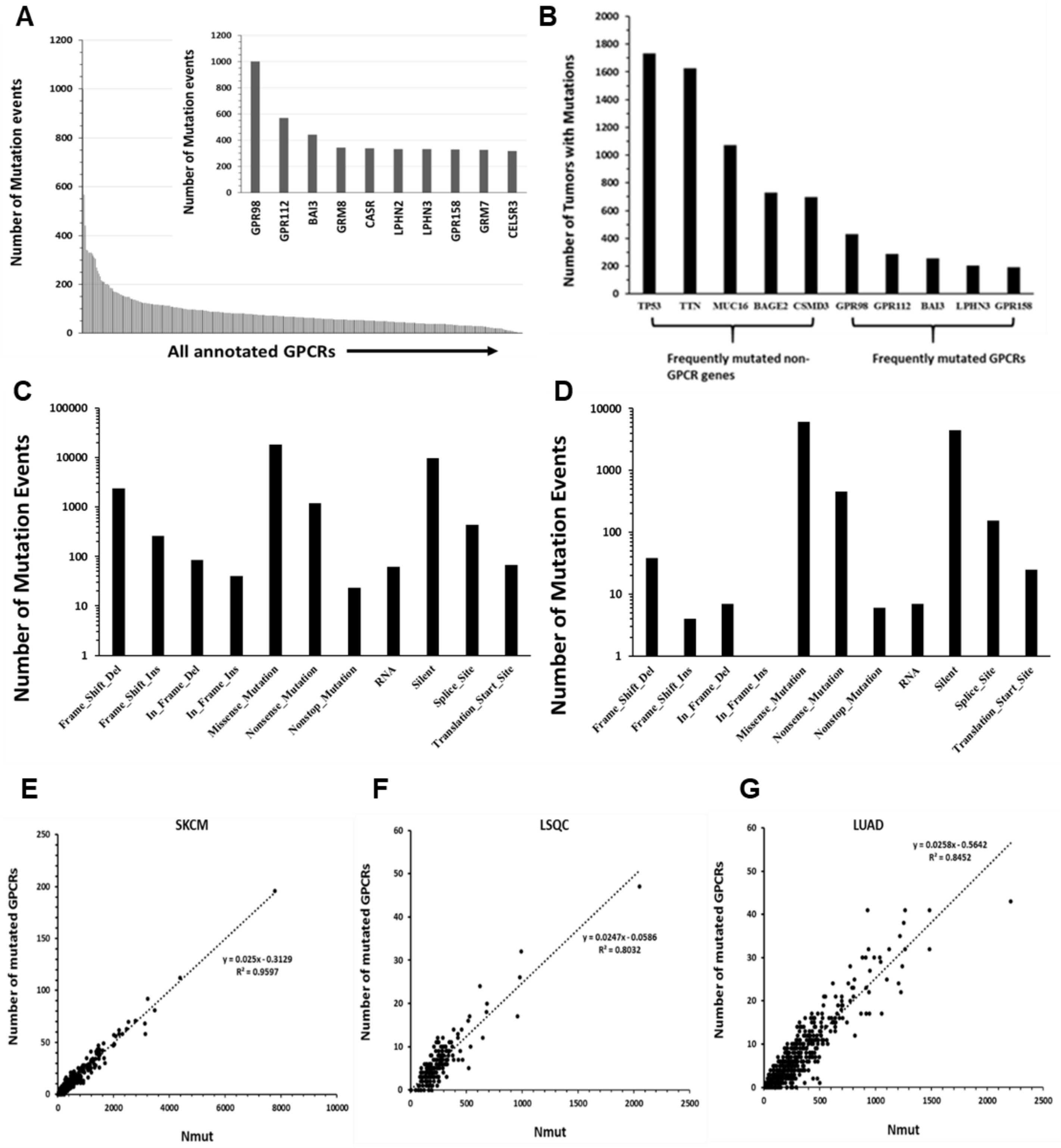

### SUPPLEMENTARY FIGURE 9

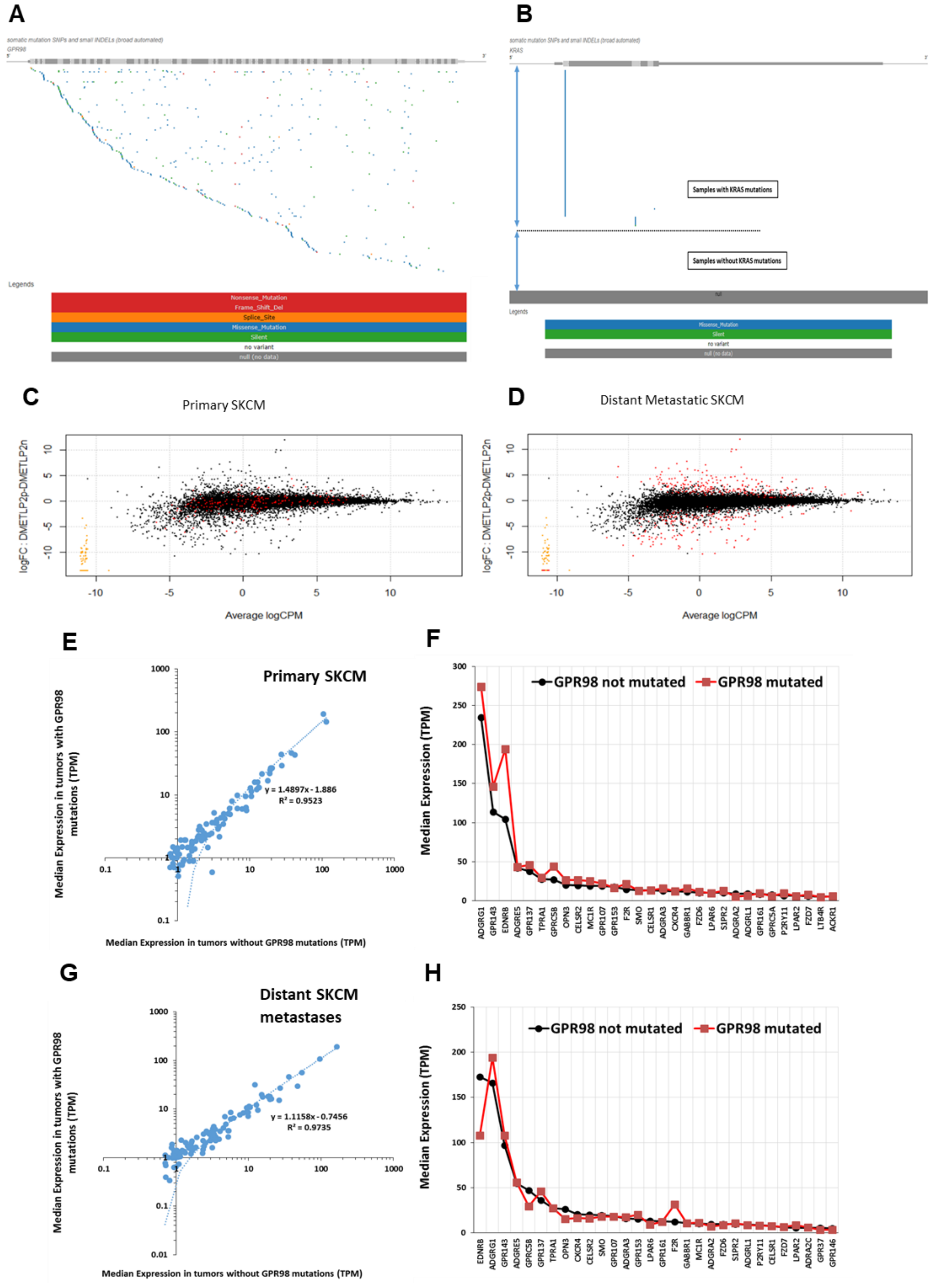

### SUPPLEMENTARY FIGURE 10

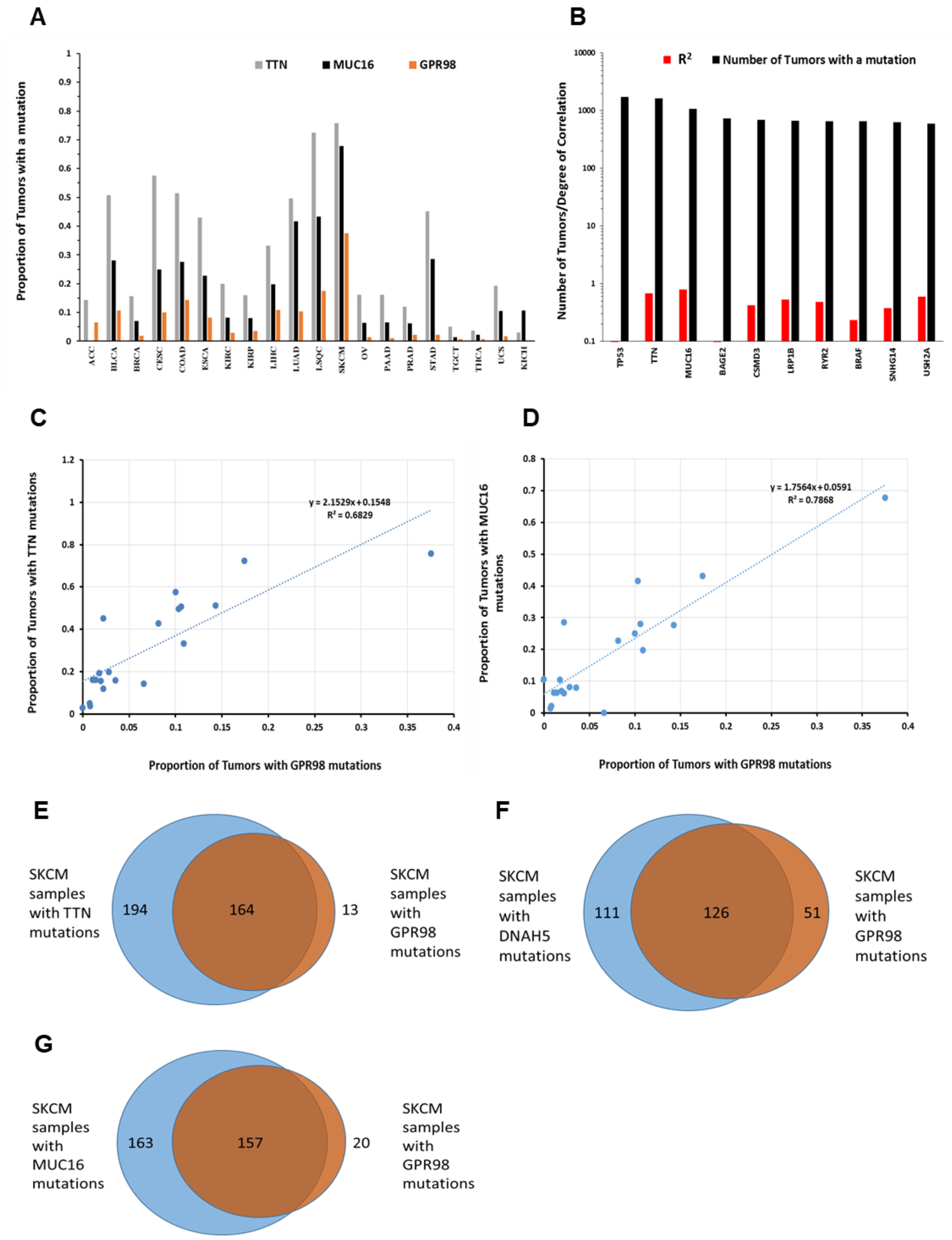

### SUPPLEMENTARY FIGURE 11

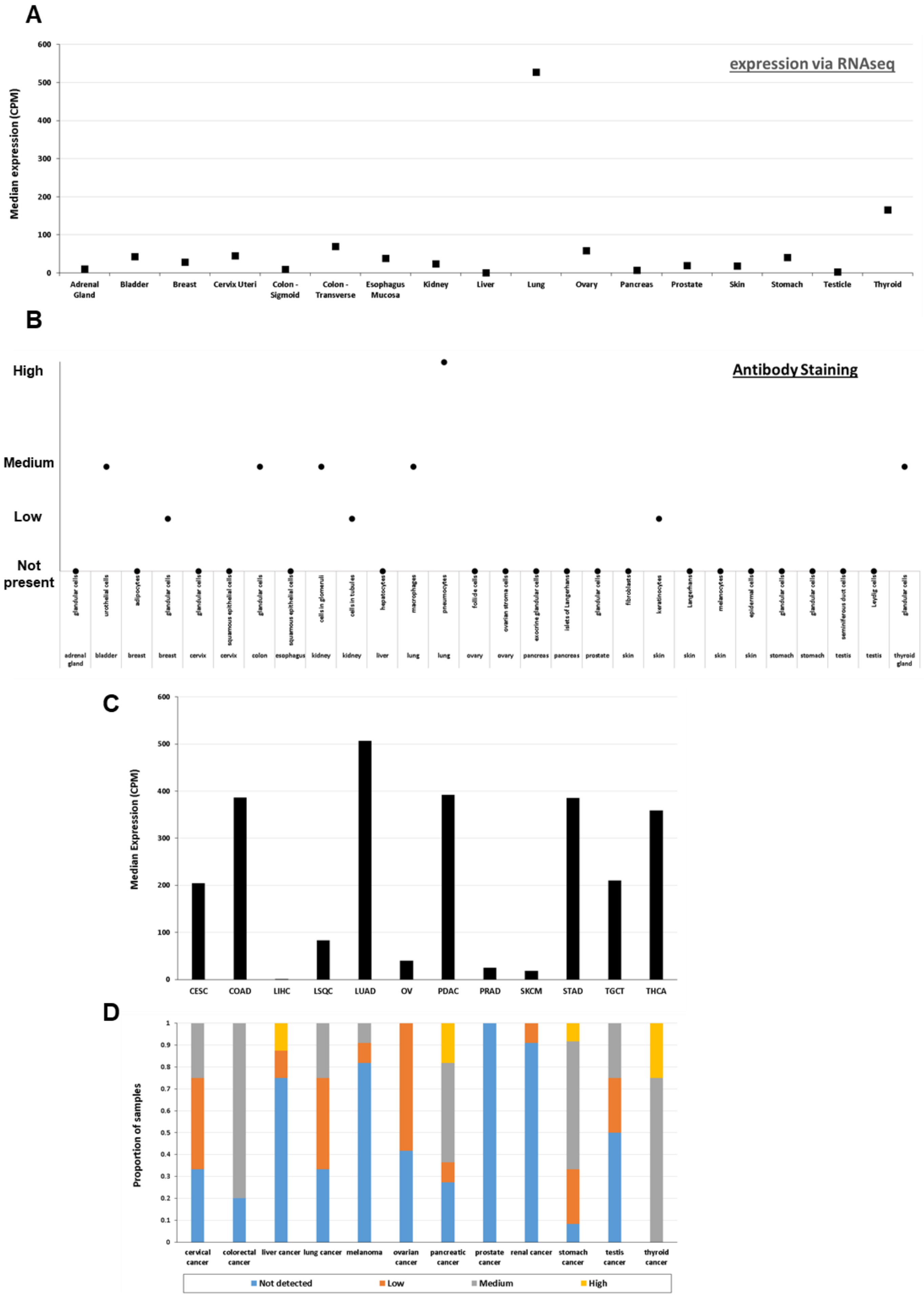

### SUPPLEMENTARY FIGURE 12

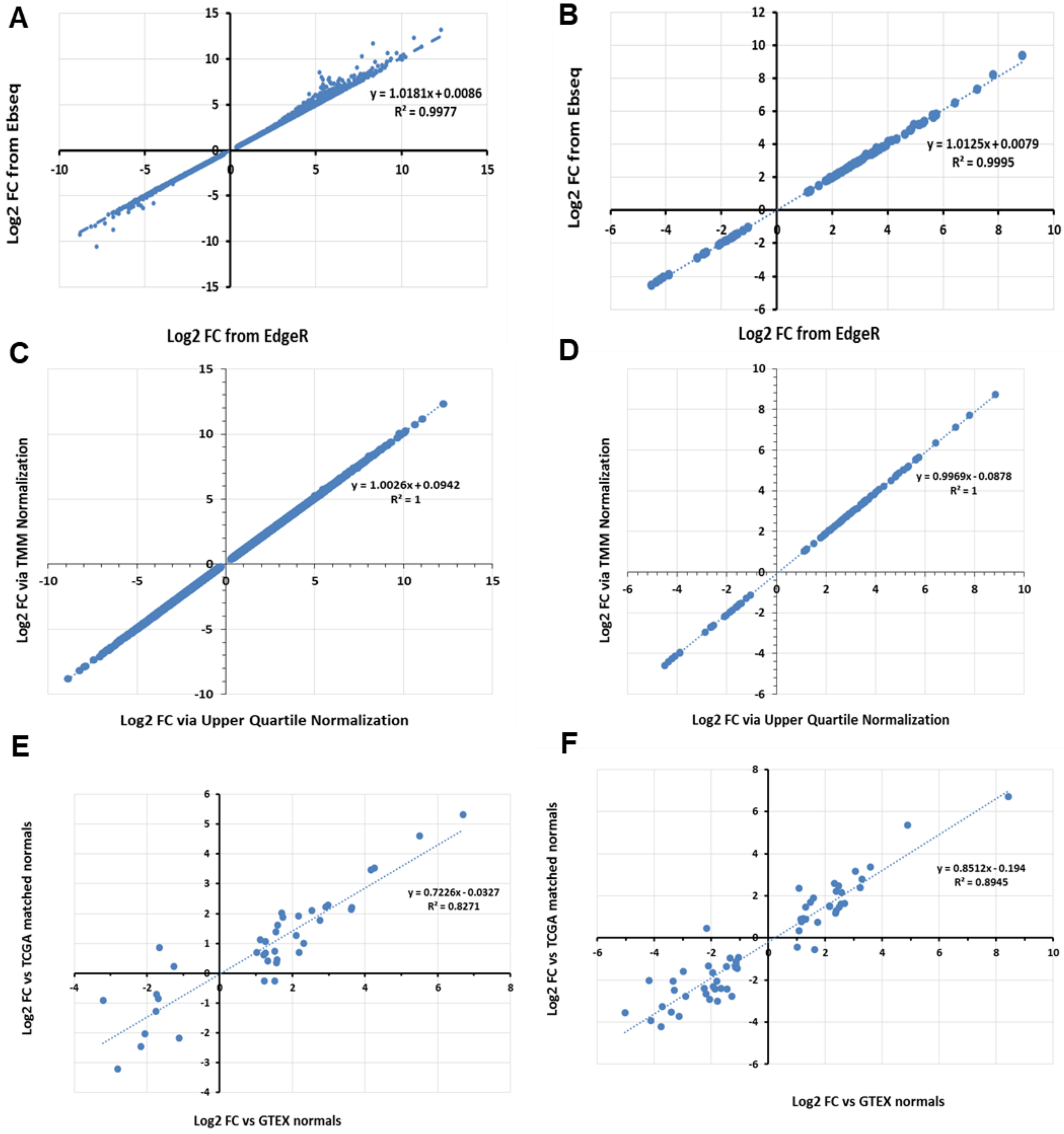

### SUPPLEMENTARY FIGURE 13

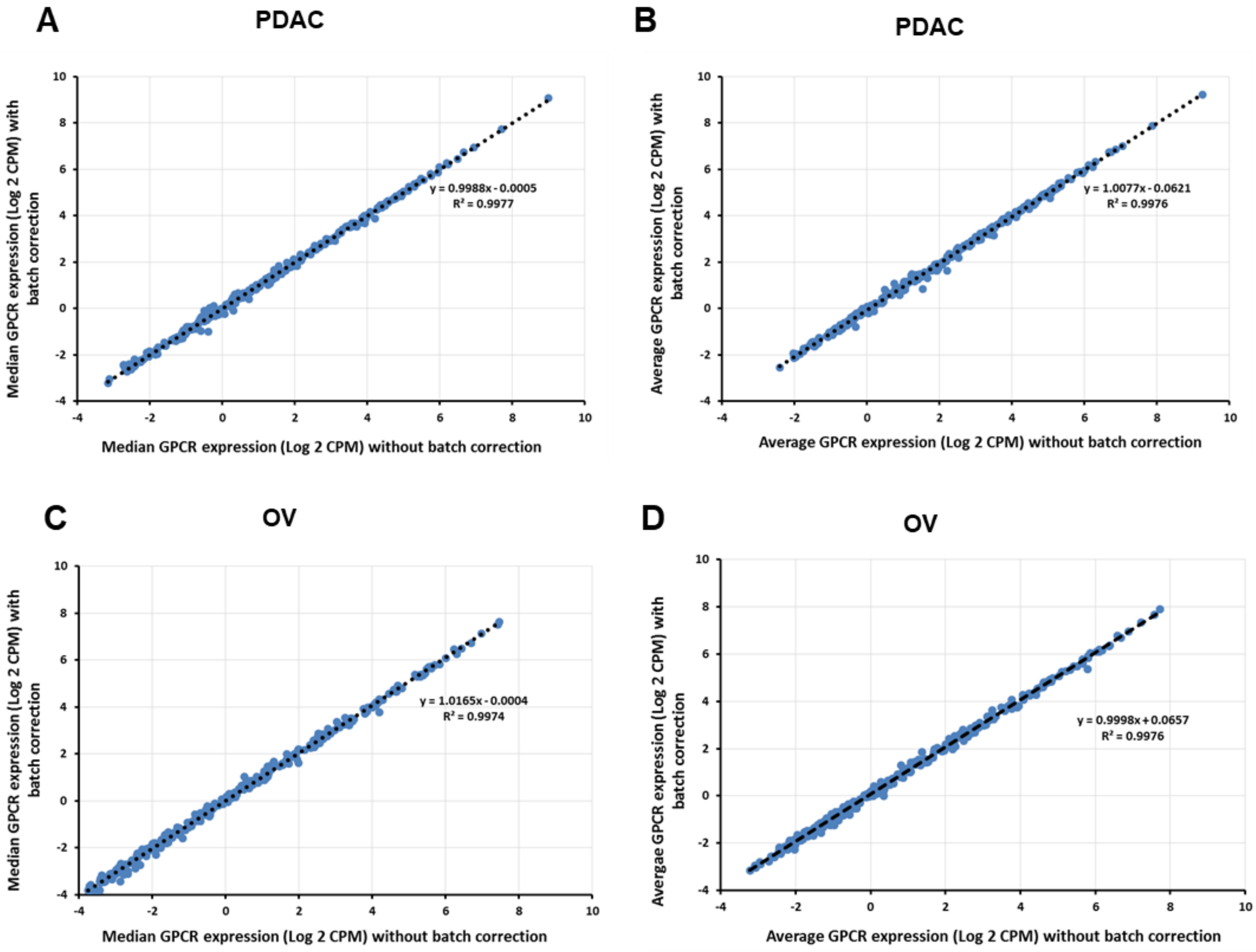
